# A new class of penicillin-binding protein inhibitors to address drug-resistant *Neisseria gonorrhoeae*

**DOI:** 10.1101/2024.12.27.630553

**Authors:** Tsuyoshi Uehara, Allison L. Zulli, Brittany Miller, Lindsay M. Avery, Steven A. Boyd, Cassandra L. Chatwin, Guo-Hua Chu, Anthony S. Drager, Mitchell Edwards, Susan G. Emeigh Hart, Cullen L. Myers, Gopinath Rongala, Annie Stevenson, Kyoko Uehara, Fan Yi, Bibo Wang, Zhenwu Liu, Mingyue Wang, Zhichao Zhao, Xinming Zhou, Haiyan Zhao, Caleb M. Stratton, Sandeepchowdary Bala, Christopher Davies, Rok Tkavc, Ann E. Jerse, Daniel C. Pevear, Christopher J. Burns, Denis M. Daigle, Stephen M. Condon

**Author notes:** These authors contributed equally (Allison L. Zulli, Brittany Miller).

## Abstract

β-Lactams are the most widely used antibiotics for the treatment of bacterial infections because of their proven track record of safety and efficacy. However, susceptibility to β-lactam antibiotics is continually eroded by resistance mechanisms. Emerging multidrug-resistant (MDR) *Neisseria gonorrhoeae* strains possessing altered *penA* alleles (encoding PBP2) pose a global health emergency as they threaten the utility of ceftriaxone, the last remaining outpatient antibiotic. Here we disclose a novel benzoxaborinine-based penicillin-binding protein inhibitor series (boro-PBPi) that is envisioned to address *penA-*mediated resistance while offering protection against evolution and expansion of β-lactamases. Optimization of boro-PBPi led to the identification of compound **21** (VNRX-14079) that exhibits potent antibacterial activity against MDR *N. gonorrhoeae* achieved by high affinity binding to the PBP2 target. Boro-PBPi/PBP2 complex structures confirmed covalent interaction of the boron atom with Ser310 and the importance of the β_3_-β_4_ loop for improved affinity. **21** elicits bactericidal activity, a low frequency of resistance, a good safety profile, suitable pharmacokinetic properties, and in vivo efficacy in a murine infection model against ceftriaxone-resistant *N. gonorrhoeae*. **21** is a promising anti-gonorrhea agent poised for further advancement.

## INTRODUCTION

For the treatment of bacterial infections, β-lactam antibiotics, including penicillins, cephalosporins, monobactams, and carbapenems, represent over 60% of the total antibiotic market, reflecting their proven safety and efficacy record over >70 years of clinical use^1^. However, bacterial pathogens are increasingly resistant to β-lactam drugs, mainly due to the rapid evolution of β-lactamase enzymes that chemically inactivate these clinically important agents^2, 3^. The therapeutic targets of β-lactams are the penicillin-binding proteins (PBPs), which are unique to bacteria, essential for cell wall synthesis, and indispensable for bacterial proliferation^4, 5^. The clinical success of β-lactams validates PBPs as extremely valuable antibacterial targets. To overcome β-lactamase-mediated resistance, combination therapies of BL with β-lactamase inhibitors (BLI) have been developed, but resistance to these β-lactam/BLI combinations is rapidly developing^6^.

An alternative approach that enables continued targeting of PBPs is to develop non-β-lactam PBP inhibitors that avoid degradation by β-lactamases^5^. Since serine β-lactamases and PBPs share similar active site residues and catalytic mechanism, PBP inhibition utilizing current β-lactamase inhibitor scaffolds that are refractory to inactivation by β-lactamases (e.g., diazabicyclooctanes (DBOs) and benzoxaborinine-based inhibitors) has been explored^5, 7^. Since non-β-lactam PBP inhibitors will likely evade the action of all known and future β-lactamases, these novel PBP inhibitors may circumvent the evolution of β-lactamases and reset the resistance clock for PBP targeting agents.

Gonorrhea, caused by *Neisseria gonorrhoeae,* is the second most prevalent sexually transmitted bacterial infection with an estimated 82.4 million cases worldwide in 2020^8^. If left untreated, gonorrhea can lead to serious health complications, including infertility in both men and women, pelvic inflammatory disease, and increased risk of HIV/AIDS^8^. Since the introduction of the sulfa drugs in the 1930s, *N. gonorrhoeae* has evolved under constant antimicrobial drug pressure. As such, owing to acquired resistance to nearly every antibiotic class including sulfa drugs, penicillins, tetracycline, ciprofloxacin, azithromycin, and cephalosporins, there has been a parallel evolution of therapeutic recommendations^9, 10, 11, 12, 13, 14^. Although gonorrhea is treatable and can be cured by current antibiotics, the emergence of *N. gonorrhoeae* resistant to antibiotics is making treatment of gonorrhea more challenging, forecasting future failings of outpatient therapy. Therefore, multidrug-resistant *N. gonorrhoeae* has been classified as an “urgent threat” by the Centers for Disease Control and Prevention (CDC) and the World Health Organization (WHO) has declared dual cephalosporin-resistant and fluoroquinolone-resistant *N. gonorrhoeae* as a “High Priority” pathogen for the discovery and development of new antibiotics^15, 16, 17^. The current last-line therapeutic option to treat uncomplicated gonorrhea is a single intramuscular (IM) dose (250–1000 mg) of the third-generation cephalosporin, ceftriaxone (CRO)^18^. Unfortunately, global CRO-resistant *N. gonorrhoeae* infections are increasing in frequency, foreshadowing an era of untreatable gonorrhea infections in the outpatient setting that would result in an unsustainable burden on healthcare systems^19, 20^. As there are no effective vaccines, antibiotic therapy remains the only option for treatment^21^.

Protein variants of *penA*, encoding penicillin-binding protein 2 (PBP2) (functionally equivalent to the cell division protein PBP3 in *Escherichia coli*), are the predominant mechanism of CRO-resistance in *N. gonorrhoeae* with the most urgent threat mediated by the mosaic *penA-60* allele (a "mosaic PBP2" refers to a variant of PBP2 protein composed of segments derived from several *Neisseria* species and are often associated with decreased susceptibility to cephalosporin antibiotics)^13, 20^. Notably, *N. gonorrhoeae* strains possessing *penA-60* have been spreading globally since 2015 when it was first discovered in the FC428 strain isolated in Japan^19, 22, 23, 24, 25, 26, 27^. In addition, *N. gonorrhoeae* can acquire genetic elements to produce an extended spectrum β-lactamase (ESBL) or carbapenemase which can hydrolyze CRO and other third generation cephalosporins^28^. Although production of the ESBL TEM-20 imposes a fitness cost in *N. gonorrhoeae*^29^, the recent report of an *N. gonorrhoeae* clinical isolate producing the ESBL CTX-M-1 serves as an early harbinger of the possible emergence of ESBL-mediated CRO resistance^30^.

Of the recent agents developed for the treatment of multi-drug-resistant *N. gonorrhoeae*, only zoliflodacin and gepotidacin, both of which target bacterial DNA replication (i.e., topoisomerases/gyrase inhibitors), have successfully completed phase 3 clinical evaluation for uncomplicated gonorrhea^31, 32, 33^. Although both agents are expected to gain approval, there continues to be a need for new anti-gonorrhea agents **—** particularly those with differing modes of action. Notably, while zoliflodacin-resistant strains have only been isolated in vitro, gepotidacin-resistant strains have already been identified within the community setting^34^. These observations portend emerging resistance to even these novel agents^35, 36^. Aside from the topoisomerase/gyrase inhibitors, single-dose IM-ertapenem (1 g) was recently found to be non-inferior to IM-CRO (500 mg) in a phase 3 trial^37^. Clearly, there is an urgent need for the discovery and development of additional safe and effective anti-gonorrhea therapies for use in the outpatient setting.

We have identified a series of non-β-lactam inhibitors of PBP transpeptidase function, i.e., PBPi, based on the α-alkylamido benzoxaborinine scaffold also present in the novel BLIs taniborbactam^38, 39^ and ledaborbactam etzadroxil (**Figure S1**)^40, 41^. Owing to the similarity within the β-lactamase and PBP active sites, boronate-based inhibitors can exploit several common binding elements, including the slowly reversible interaction of the active site serine hydroxyl group with the boron atom of the benzoxaborinine core^38, 39, 42, 43, 44^. Here we present a novel benzoxaborinine-based PBP inhibitor series (boro-PBPi) that exhibits activity against CRO-resistant, mosaic PBP2-containing *N. gonorrhoeae* variants. Moreover, we provide in vitro and in vivo data to support further preclinical development of boro-PBPi **21** (VNRX-14079) as a potential new agent for the treatment of *N. gonorrhoeae* infections, including those resistant to the current standard of care agent, CRO.

## RESULTS

### Identification of benzoxaborinine-based PBPi with stand-alone antibacterial activity

To adapt the BLI-focused benzoxaborinine template for use as a PBPi, we first introduced the aminothiazole-containing oxyimino-acetamide sidechain present in the third generation cephalosporins, cefdinir, and CRO (**Figure S2**), onto the benzoxaborinine warhead, generating boro-PBPi **1** (**Figure 1**). In vitro testing of **1** against a selected panel of Gram-negative organisms revealed modest single-agent activity against certain *E. coli* and *K. pneumoniae* strains (**Table S1**). Notably, **1** exhibited only 4-fold improved activity (minimal inhibitory concentration, MIC of 8 µg/mL) against the hyperpermeable *E. coli* BAS901C strain relative to the ATCC 25922 wild type strain, suggesting that its antibacterial activity might be limited by weak affinity for target PBPs (*E. coli* PBP3 IC_50_ of 113 µM). Importantly, **1** possessed modest activity against non-mosaic PBP2-producing *N. gonorrhoeae* ATCC 49226 (MIC of 16 µg/mL, relative to ceftriaxone MIC of 0.008 µg/mL) with an IC_50_ of 100 µM against wild type PBP2 (PBP2^WT^) isolated from the *N. gonorrhoeae* FA19 strain. However, **1** was inactive (MIC of >128 µg/mL) against the H041 strain producing the mosaic PBP2 that mediates CRO-resistance.

**Figure 1.**
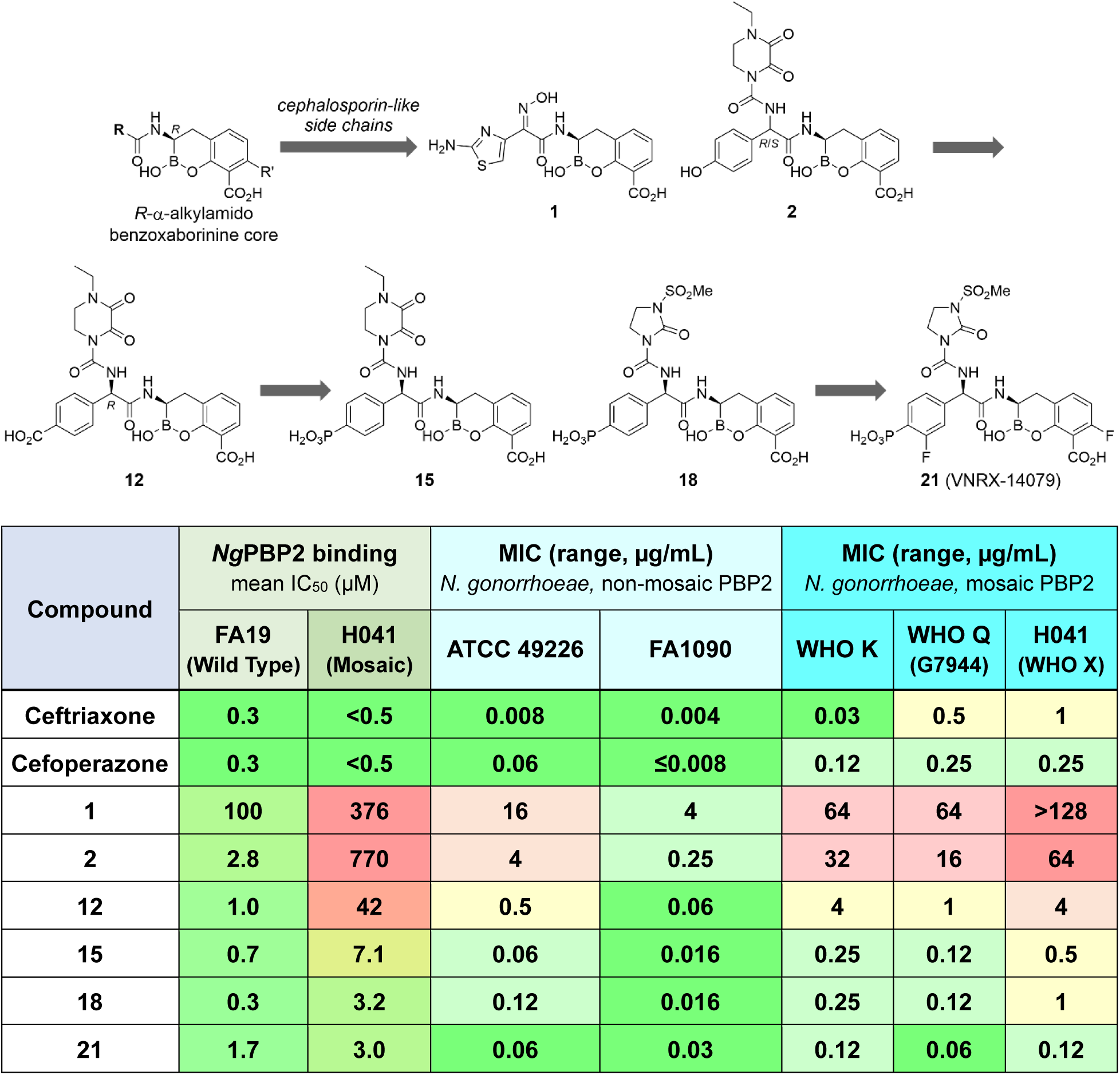
Evolution of boro-PBPi from BLI-containing benzoxaborinine core to ureido-containing lead compound 21 (VNRX-14079). The table shows PBP2 binding and MIC data for ceftriaxone, cefoperazone, and selected boro-PBPi.

### Incorporation of the ureido moiety for improved activity against *N. gonorrhoeae*

We next chose to replace the oxyimino amide sidechain with the ureido-based aryl glycine motif present in piperacillin and cefoperazone (**Figure S2**). The initial ureido-containing benzoxaborinines were prepared as near equimolar mixtures of the *R*,*R*- and *S*,*R*-diastereomers (**Supplementary Information**); when required, the desired *R*,*R*-diastereomers (*infra*) were isolated following preparative reversed-phase high performance liquid chromatography (RP-HPLC). The parent phenol-bearing boro-PBPi **2** (**Figure 1**), a benzoxaborinine linked to the ureido side chain of cefoperazone (**Figure S2**), showed moderate antibacterial activity against representative *N. gonorrhoeae* strains producing non-mosaic PBP2 and exhibited good in vitro binding to PBP2^WT^ (IC_50_ of 2.8 µM). However, poor binding affinity was observed for **2** against the mosaic PBP2 isolated from the CRO-resistant H041 strain (PBP2^H041^, IC_50_ of 770 µM). Correspondingly, across a panel of *N. gonorrhoeae* strains, MICs for **2** were higher in the mosaic PBP2-containing strains compared to the strains with non-mosaic PBP2 (**Figure 1**).

A series of phenol-bearing boro-PBPi was prepared and evaluated for PBP2^WT^ binding and antibacterial activity against both CRO-susceptible ATCC 49226 and CRO-resistant H041 strains – an abbreviated, yet informative list of analogs is provided in **Figure S3**. The introduction of a fluorine atom *ortho* to the parent phenol provided *R*,*R*-**3** as a single isomer after RP-HPLC-mediated separation of the diastereomers. In both CRO-susceptible and resistant strains, **3** exhibited comparable MICs to **2,** which had been tested as a ∼1:1 mixture of active and inactive diastereomers. *R*,*R*-**4** proved less active against ATCC 49226 than **3**, while exhibiting only modest activity against the CRO-resistant H041 strain (MIC of 16 µg/mL). Additional fluorinated analogs **5**–**9** revealed improved PBP2^WT^ binding relative to non-fluorinated **2** and had comparable or lower MIC values against ATCC 49226. Although most of these analogs plateaued at MIC of 16– 32 µg/mL when tested against CRO-resistant H041, trifluorinated **9** displayed a slightly improved MIC of 8 µg/mL against this mosaic PBP2-containing strain (**Figure S3**), suggesting that inclusion of fluoro groups may have a positive effect on PBP2 binding and/or periplasmic accumulation.

To better understand the relationship between fluoro substitution and PBP2 binding and/or antibacterial activity, the p*K*_a_ of the phenol proton for these modestly active analogs was calculated. While unsubstituted phenol has a p*K*_a_ of 10.0, the inductive effect of fluoro substitution lowers the p*K*_a_ of 2-fluoro- and 3-fluoro-phenol to 8.7 and 9.3, respectively. The 2,3,6-trifluoro-4-hydroxyphenyl glycine of **9** was calculated to have a p*K*_a_ (Ar-O<u>H</u>) of 6.7, which implies ionization of the phenol at physiological pH. These data suggest that **9** likely exists as a phenoxide at physiological pH, which may be contributing to the observed antibacterial activity of these fluorine-containing boro-PBPi.

### Benzoate- and phosphonate-containing benzoxaborinines improves boro-PBPi activity

Based on the hypothesis that a phenoxide anion might be beneficial at the *para* position of the aryl glycine moiety, a series of benzoate-containing boro-PBPi was investigated, with the expectation that a carboxylate residue might be similarly preferred. To this end, a suitable benzoate-based phenyl glycine intermediate (**42**) was prepared (**Supplementary Information**). The results from this set of benzoate-containing benzoxaborinines supported the hypothesis that an anion is preferred at the 4-position of the aryl glycine moiety. Both **10** and **11** displayed good activity against ATCC 49226 (MIC of 1 and 4 µg/mL, respectively), and importantly, both boro-PBPi exhibited activity against H041 (MIC of 8 µg/mL) at a level comparable to trifluorinated phenol **9** (**Figure S3**), with **11** showing measurable binding to mosaic PBP2^H041^ (**Figure S4**). Separation of the active isomer of **10**, i.e., *R*,*R*-**12**, allowed identification of the first benzoxaborinine-based PBPi with MIC of 4 µg/mL against H041 (**Figure 1**). Both fluorinated single isomers, *R*,*R*-**13** and *R*,*R*-**14**, also showed MICs of 4 µg/mL against H041 (**Figure S4**), further supporting this subseries as a promising lead benzoxaborinine PBPi for CRO-resistant gonococcal infection.

Seeking to identify alternatives to the putative phenoxide and carboxylate hydrogen bonding residues, a phosphonate group was incorporated onto the phenyl glycine moiety, preparing the non-fluorinated parent boro-PBPi *R*,*R*-**15** (**Figure 1**). Strikingly, the addition of a phosphonate residue to these boro-PBPi resulted in a marked improvement in mosaic PBP2^H0^^41^ binding and antibacterial activity against both CRO-susceptible and -resistant strains when compared to the phenol- or benzoate-containing boro-PBPi (**Figure 1** and **Figure S5**). The 2,3-dioxapiperidine-containing analogs **15**–**17** showed good binding to WT and mosaic PBP2 and excellent MICs of 0.06–0.12 and 0.5 µg/mL against ATCC 49226 and H041 strains, respectively.

Having discovered the importance of the terminal phosphonate group for PBP2 engagement and bactericidal activity, a series of boro-PBPi bearing the *N*-methylsulfonyl-2-imidazolinone motif present in mezlocillin was prepared (**Figures S2** and **S6**). These new boro-PBPi, represented by *R*,*R*-**18** (**Figure 1**), had similar PBP2 binding affinity and antibacterial activity as the 2,3-dioxapiperidine-containing analogs. The incorporation of multiple fluorine atoms (boro-PBPi **19**– **23**) led to *R*,*R*-**21** (VNRX-14079) that showed a 2–8-fold improvement in MICs relative to **18** in mosaic PBP2-producing strains. In addition, **21** exhibited 8-fold lower MICs than CRO against the CRO-resistant H041 and WHO Q strains that produce mosaic PBP2^H0^^41^ and PBP2^FC4^^28^ (encoded by *penA-60*), respectively.

### Structural evidence for high affinity binding of boro-PBPi to mosaic PBP2

Mutations in the transpeptidase (TPase) domain of PBP2 are firmly established as the primary determinant of cephalosporin resistance in *N. gonorrhoeae*^45^ and such mutations appear to act by restricting the conformational dynamics of the protein^46, 47^. We obtained the structures of benzoate-containing **12** and phosphonate-containing **15** complexes in the mosaic PBP2 transpeptidase domain isolated from the extended-spectrum-cephalosporin-decreased susceptibility strain 35/02, i.e., tPBP2^35/02^ (**Figure 2**)^47^. In both structures, essentially complete electron density is observed for the inhibitors, including a covalent interaction between the boro-PBPi boron atom and Ser310 of the tPBP2^35/02^ active site. In addition, the benzoxaborinine-associated carboxylate group is hydrogen bonded to Thr498 and Thr500, while Ser362 forms a hydrogen bond with the boronate ester oxygen atom. Furthermore, a hydrogen bond between Asn364 and the carbonyl group of the α-amido benzoxaborinine is observed. Notably, the *para-*carboxylate of **12** forms a hydrogen bond with Tyr543 in the β_5_-α_11_ loop (**Figure 2A**). Replacement of the benzoate moiety with a phosphonate residue in **15** leads to the formation of additional hydrogen bonds with Arg502 and His514, in addition to that with Tyr543 (**Figure 2B**).

**Figure 2.**
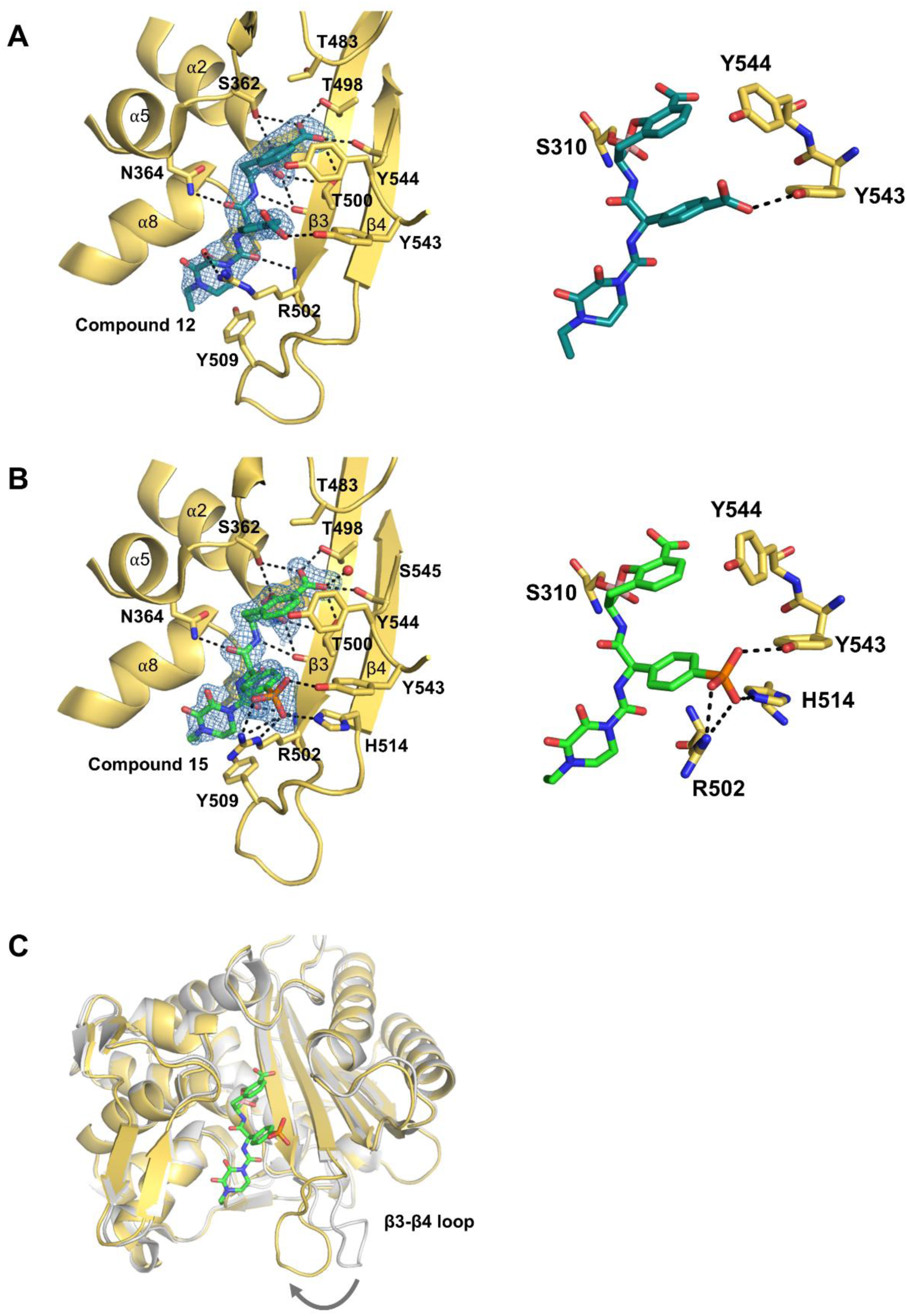
Crystal structures of tPBP2^35/02^ in complex with boro-PBPi 12 and 15. **A** and **B**) The crystal structures of mosaic *Ng* tPBP2^35/02^ bound to *R*,*R*-**12** and *R*,*R*-**15**, respectively. Right hand panels show the interactions made by the *para*-carboxylate in **12** and phosphonate of **15**. **C**) Superimposition of the crystal structures of *Ng* tPBP2^35/02^ in complex with **15** and *apo Ng* tPBP2^35/02^ (PDB: 6VBL) showing the movement of the β_3_-β_4_ loop toward the active site in the complex (arrow).

One of the most significant conformational changes observed upon boro-PBPi binding is the movement of the β_3_-β_4_ loop from a so-called “outbent” conformation as observed for *apo* tPBP2^35/02^ to an inward conformation closer to the active site (**Figure 2C**). The inward movement of the β_3_-β_4_ loop in the crystal structures in complex with the boro-PBPi is highly significant, as its failure to move in PBP2 variants derived from extended-spectrum-cephalosporin-resistant strains of *N. gonorrhoeae* is hypothesized to be an important contributor to CRO-resistance^47^. The observed binding modes of **12** and **15**, together with the movement of the β_3_-β_4_ loop, are similar to the binding modes of cefoperazone and the ureidopenicillins piperacillin and azlocillin observed in the tPBP2^H0^^41^ protein^48^. These β-lactams exhibit faster second-order acylation rates against tPBP2^H0^^41^ compared to the oxyimino-containing CRO^48^. These similar findings with the boro-PBPi are therefore consistent with the idea that higher-activity inhibitors work by overcoming the conformational barrier created by resistance mutations present in tPBP2 from resistant strains.

### Pharmacokinetic evaluation and in vivo efficacy of boro-PBPi

In advance of the murine efficacy studies^49^, the plasma protein binding (PPB) and pharmacokinetic (PK) profiles of **18** and **21** were determined in mice. In mouse plasma, PPB of **18** and **21** was 7% and 19%, respectively, whereas PPB in human plasma was 24% and 39%, respectively. As shown in **Figure S7**, boro-PBPi **18** (formulated at 100 mg/mL in PBS, pH 6.5-7.0) and **21** (formulated at 75 mg/mL in PBS, pH 6-7) exhibited similar plasma-time concentration profiles for each compound, when administered to mice by either intravenous (IV) bolus or subcutaneous (SC) injection. Boro-PBPi **18** and **21** were well tolerated up to the highest dose tested (1,000 and 300 mg/kg SC doses, respectively). Exposure, in terms of maximum concentrations (C_max_) and total overall exposures (AUC), was less than dose-proportional for **21** following SC administration from 10 to 300 mg/kg, while for **18**, exposure following SC exposure was less than dose proportional for C_max_ at ≥300 mg/kg and was generally dose-proportional for AUC_inf_ across all dose levels administered (**Tables S2 and S3**). The terminal half-life (t_½_) of **18** was approximately 4 h following SC dosing with good exposure at all doses. The terminal half-life of **21** was ∼4 h at ≥100 mg/kg, whereas the half-life at 10 and 30 mg/kg was calculated to be short at <0.9 h because the terminal half-life of the lower doses was not determined as concentrations at later timepoints were below the limit of detection.

Assuming all drugs that target PBPs display time-dependent antibacterial activity in vivo as demonstrated for β-lactam-based PBP inhibitors^50, 51^, we selected the boro-PBPi dosages of 0.25, 7.5, 50, and 200 mg/kg every 12 h (q12h or BID) for the in vivo efficacy study in the murine vaginal infection model with the CRO-susceptible FA1090 strain [agar MIC of 0.016 µg/mL for **18** and 0.008 µg/mL for CRO determined by CLSI guidelines^52^ (**Table 1**)]. In vivo efficacy of **18** was observed at ≥7.5 mg/kg SC q12h, as PK simulations predicted that these doses would provide unbound boro-PBPi concentrations in plasma above the MIC for at least 40% of the 24 h treatment duration (i.e., ≥40% *f*T>MIC); bacterial growth stasis was observed at 0.25 mg/kg SC q12h (NB: ≤13% PK-simulated *f*T>MIC) (**Figure S8**). The efficacious doses of **18** were similar to the efficacy achieved by CRO at 1 mg/kg (IV, single dose).

**Table 1.** Agar MIC of boro-PBPi and comparator agents against *N. gonorrhoeae* reference strains.

| Strain | PBP2 allele | 12 | 15 | 18 | 21 | CRO | ZOLI | GEPO | AZM | CIP | PEN |
| --- | --- | --- | --- | --- | --- | --- | --- | --- | --- | --- | --- |
| **ATCC 49226** | *penA-22* | 0.5 | 0.06 | 0.06 | 0.03 | 0.016 | 0.12 | 0.25 | 0.25 | 0.004 | **0.5** |
| **FA19** | *penA-15* | ≤0.016 | 0.008 | 0.008 | 0.008 | 0.004 | 0.03 | 0.12 | 0.06 | ≤0.002 | ≤0.016 |
| **FA1090** | *penA-1* | 0.06 | 0.016 | 0.016 | 0.016 | 0.008 | 0.03 | 0.12 | 0.06 | 0.004 | 0.06 |
| **WHO F** | *penA-15* | ≤0.016 | ≤0.004 | 0.008 | 0.008 | ≤0.002 | 0.06 | 0.12 | 0.06 | 0.004 | ≤0.016 |
| **WHO G** | *penA-2* | 0.25 | 0.03 | 0.06 | 0.03 | 0.016 | 0.12 | 1 | 0.12 | **0.12** | **0.5** |
| **WHO M** | *penA-2* | 0.5 | 0.06 | 0.06 | 0.06 | 0.016 | 0.06 | 1 | 0.25 | **2** | **>64** |
| **MS11** | *penA-22* | 0.5 | 0.03 | 0.06 | 0.03 | 0.016 | 0.12 | 0.5 | 0.25 | 0.004 | **0.5** |
| **WHO L** | *penA-7* | 8 | 1 | 1 | 0.5 | 0.25 | 0.12 | 4 | 0.25 | **16** | **2** |
| **WHO K** | Mosaic *penA-10* | 4 | 0.25 | 0.5 | 0.12 | 0.12 | 0.12 | 0.25 | 0.25 | **>16** | **2** |
| **H041** | Mosaic *penA-37* | 16 | 1 | 1 | 0.25 | **1** | 0.12 | 0.5 | 0.25 | **>16** | **2** |
| **WHO Z** | Mosaic *penA-64* | 2 | 0.25 | 0.25 | 0.06 | **0.5** | 0.12 | 0.25 | 1 | **>16** | **2** |
| **WHO Q (G7944)** | Mosaic *penA-60* | 4 | 0.5 | 0.25 | 0.06 | **0.5** | 0.06 | 1 | **>4** | **16** | **1** |
| **CDC-0197** | Mosaic *penA-34* | 2 | 0.25 | 0.25 | 0.06 | 0.06 | 0.06 | 0.25 | **>4** | **16** | **1** |
| **F89** | Mosaic *penA-42* | >16 | >8 | >8 | >8 | **2** | 0.12 | 0.5 | 0.25 | **16** | **2** |

Agar MIC (µg/mL)
Shown in **red** are resistant/non-susceptible MIC values based on the CLSI guideline<sup>77</sup> where MIC breakpoints are defined only for ceftriaxone, azithromycin, ciprofloxacin, and penicillin G.
Abbreviations: CRO, ceftriaxone; ZOLI, zoliflodacin; GEPO, gepotidacin; AZM, azithromycin; CIP, ciprofloxacin; PEN G, penicillin G.

In vivo efficacy of **21** was evaluated in the murine vaginal infection model using the CRO-resistant H041 strain (agar MIC of 0.25 µg/mL for **21** and 1 µg/mL for CRO) as previously described^51^ **(Figure S9**). Four dosing groups were selected to provide %*f*T>MIC for **21**, ranging from 20% to 100% when corrected for 19% mouse plasma protein binding. In this model, **21** showed good efficacy against the CRO-resistant H041 strain by demonstrating a statistically significant reduction in the percentage of culture-positive mice relative to vehicle control (*p* <0.0001) (**Figure 3** and **Table S5**). The 150 mg/kg SC twice-a-day (q12h) and three-times-a-day (q8h) regimens resulted in 90% and 100% bacterial clearance by day 2, respectively, whereas 200 mg/kg SC single dose (q24h) achieved 100% clearance by day 3. Results with 10 mg/kg SC q12h were also significant with 80% clearance by day 4 relative to vehicle control (p <0.01). Bacterial bioburden (CFU/mL) in vaginal swab suspension was consistent with bacterial clearance (% mice cured of infection) in all treatment groups (**Figure 3**). Based on the PK data and simulations, the in vivo breakpoint of **21** for clearance of infections by H041 was a single dose of 200 mg/kg, which corresponds to a predicted *f*T/MIC of 44% on day 1 and is consistent with that observed for **18** against FA1090 (≥40% *f*T>MIC). Notably, neither the evolution of spontaneous gonococcal mutants nor a significant change in vaginal polymorphonuclear leukocytes (PMNs) was observed between groups during this study (**Figure S10**). No adverse effects were observed in mice treated with any dose of **21**.

**Figure 3.**
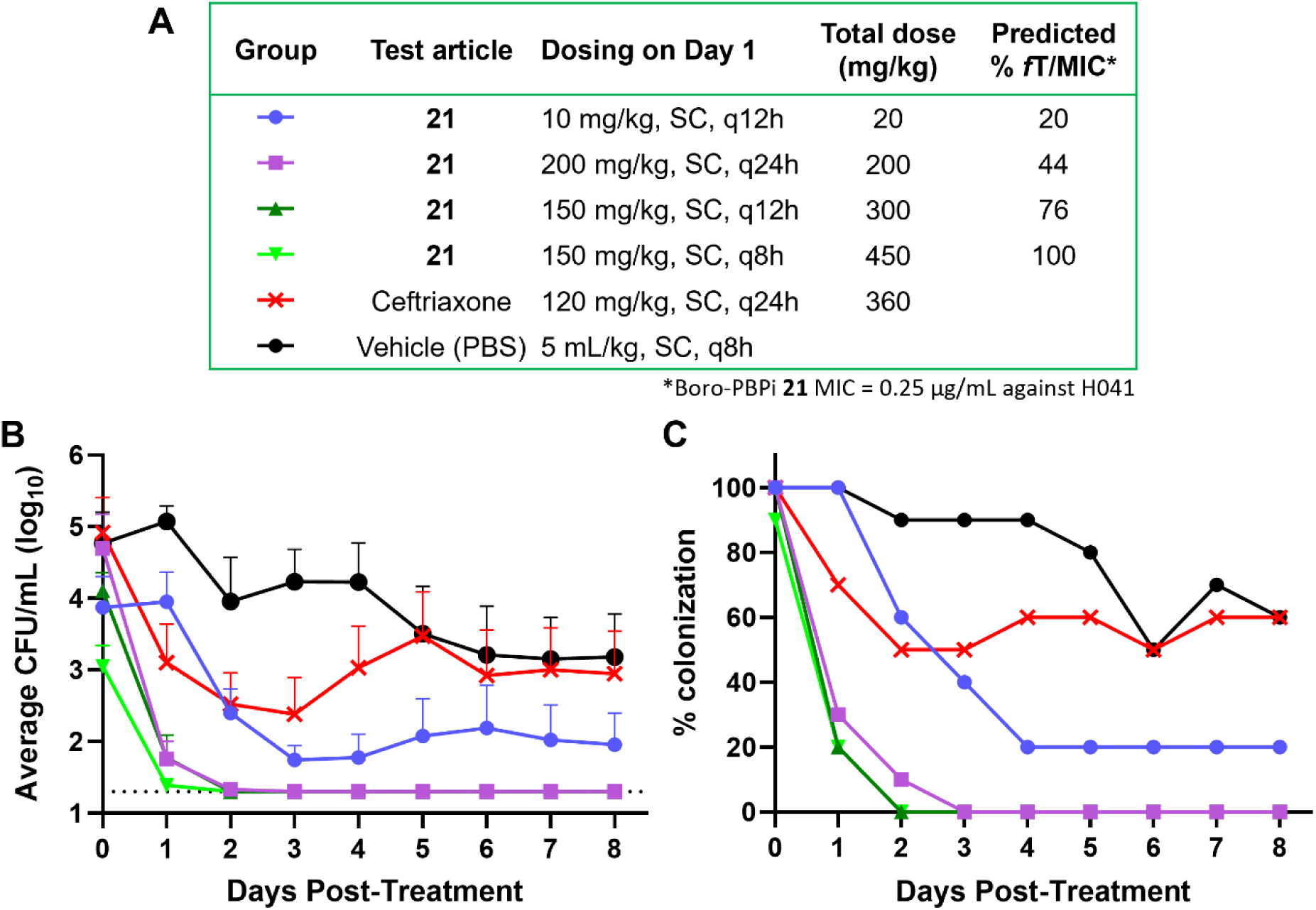
In vivo efficacy of boro-PBPi 21 in the murine vaginal infection model with ceftriaxone-resistant *Neisseria gonorrhoeae* H041. **A)** The dosing scheme to examine in vivo efficacy of **21** against the ceftriaxone-resistant H041-STM^R^ strain in the murine vaginal infection model is shown with predicted % *f*T/MIC. **21** was dosed only on day 1. Vaginal swab samples from infected mice were cultured to confirm H041 infection on day 0 prior to administration of test compound or controls on day 0 (green box). Vaginal cultures were collected on days 1–8 post treatment to determine the bacterial burden (CFU/mL). **B)** The percentage of mice infected over time with H041, and the average bacterial burden recovered over the course of the experiment. Over 85% of vehicle control mice remained colonized through day 5, with 60% colonized by day 8. SC-administered CRO, used as a positive control, yielded unanticipated underperformance relative to traditional intraperitoneal (IP) injection^51^. The horizontal dashed line denotes the limit of detection (20 CFU/mL). **C**) The average bacterial burden recovered over the course of the experiment. Error bars indicate standard error of the mean. All bacterial burden data are presented in **Table S5**.

### Microbiological activity of boro-PBPi 21

**21** showed comparable MICs to CRO against strains producing non-mosaic PBP2 and lower MICs against mosaic PBP2-producing strains, including the CRO-non-susceptible strain WHO Q, which belongs to the globally expanding FC428 lineage (**Tables 1** and **S7**). The WHO M strain that produces TEM-1 β-lactamase was susceptible to **21**, which is consistent with the stability of benzoxaborinines towards β-lactamase inactivation. Two strains, WHO L (containing non-mosaic PBP2) and F89 (containing mosaic PBP2), were the least susceptible strains to **21**. WHO L carries a PBP2 A501V variant (**Table S8**) that reduces CRO susceptibility^53, 54^ and likely also affects activity of **21**. The F89 strain has mosaic PBP2 (*penA*-42) which is identical to PBP2 (*penA*-34) of the CDC-0197 strain but with an additional A501P alteration (**Table S8**). The difference in MIC of boro-PBPi against *N. gonorrhoeae* F89 and CDC-0197 strains indicates that the A501P mutation impacts the activity of multiple boro-PBPi including **21**. Notably, F89 and closely related strains were isolated in France and Spain in 2010 and 2011^55, 56^, but since then, no clinical isolate carrying a PBP2 A501P variant has been reported likely owing to the fitness costs of producing this mosaic PBP2^57^.

Zoliflodacin and gepotidacin, two novel topoisomerase inhibitors, are the most advanced agents in clinical development for the treatment of gonorrhea. Zoliflodacin, which had comparable activity to **21** in the mosaic PBP2-producing strains (except F89 as stated above), was the most active comparator agent (**Table 1**). Gepotidacin was slightly less active than zoliflodacin against the strain panel. As *N. gonorrhoeae* have evolved resistance to all current or former antibiotic regimens, resistance to these new agents is anticipated as well. In fact, resistance to these inhibitors mediated by topoisomerase mutations has already been reported^34, 35, 36^. Since the molecular target of boro-PBPi like **21** is PBP2, no cross-resistance to the topoisomerase inhibitors zoliflodacin or gepotidacin is expected. To confirm this, we isolated mutants with reduced susceptibility to zoliflodacin from WHO K (all mutants possessing GyrB D429N) and showed no cross-resistance to **21** in these mutants (**Table S9**).

Boro-PBPi were tested against the CDC panel of *N. gonorrhoeae* clinical isolates (n = 44) that include isolates with diverse antimicrobial susceptibilities to a variety of known drugs used to treat gonorrhea^58^. **21** showed MIC_50_ and MIC_90_ of 0.06 and 0.12 µg/mL, respectively, which were equivalent to CRO and slightly more active than zoliflodacin (**Tables 2 and S10**). As expected, based on the primary strain panel data, **18** showed a 4-fold higher MIC_50_ and MIC_90_ than **21**, while the MIC_50_/MIC_90_ of benzoate-containing **12** was 8-fold higher than **18**. In time-kill assays using three strains (ATCC 49226, WHO Q, and H041), both **21** and CRO reduced CFU/mL by ≥2 log_10_ in 6 h and below limit of detection at 24 h relative to untreated controls (i.e., 10^8^ CFU/mL by 6 h) (**Figure S11**), confirming similar levels of bactericidal activity of **21** to CRO. The frequencies of resistance (FoR) to **21** and CRO were determined using ATCC 49226, WHO Q, and H041 on agar containing 4× and 16× MIC of each agent. No colonies arose on agar containing either **21** or CRO by 24 h, establishing FoR as <10^-9^ against all three strains (**Table S11**).

**Table 2.** MIC_50_/MIC_90_ and MIC distribution of boro-PBPi and competitor compounds against 44 CDC isolates.

| Compound | MIC<sub>50</sub> | MIC<sub>90</sub> | Number of strains at agar MIC (µg/mL) |  |  |  |  |  |  |  |  |  |
| --- | --- | --- | --- | --- | --- | --- | --- | --- | --- | --- | --- | --- |
|  |  |  | ≤0.016 | 0.03 | 0.06 | 0.12 | 0.25 | 0.5 | 1 | 2 | 4 | ≥8 |
| **Ceftriaxone** | 0.06 | 0.12 | 4 | 2 | 25 | 12 | 1 |  |  |  |  |  |
| **12** | 2 | 4 |  |  | 2 | 1 | 1 |  | 2 | 22 | 15 | 1 |
| **15** | 0.25 | 0.5 | 3 | 1 |  | 3 | 21 | 16 |  |  |  |  |
| **18** | 0.25 | 0.5 | 3 | 1 |  | 3 | 16 | 21 |  |  |  |  |
| **21** | 0.06 | 0.12 | 3 | 1 | 21 | 18 | 1 |  |  |  |  |  |
| **Zoliflodacin** | 0.12 | 0.12 |  |  | 8 | 32 | 4 |  |  |  |  |  |
| **Gepotidacin** | 0.5 | 1 |  |  |  |  | 7 | 21 | 16 |  |  |  |
| **Azithromycin** | 0.5 | 1 |  |  |  | 1 | 18 | 17 | 4 |  | 1 | 3 |
| **Ciprofloxacin** | 16 | 16 | 5 |  |  |  |  |  | 1 | 7 | 2 | 29 |
| **Penicillin G** | 2 | 2 |  |  |  | 3 |  |  | 12 | 25 | 2 |  |

MIC values against each strain are shown in **Table S10**.

### Boro-PBPi 21 exhibits promising pharmaceutical properties

**21** showed no hemolytic activity when tested at 1 mg/mL in human red blood cells, no cytotoxicity at 256 µg/mL against 3 mammalian cell lines (HeLa, MRC-5, and 3T3), no mitochondrial toxicity (IC_50_ >100 µM) in a Glu/Gal assay using SKOV-3 human ovarian cancer cells, and no chromosomal aberrations (IC_50_ >300 µM) in the micronucleus assay using CHO-K1 cells with and without liver S9 fraction pre-incubation (**Tables 3 and S12**). When tested at 30 µM, **21** showed no noteworthy inhibition (≥10% decrease of baseline activity) of seven CYP enzymes (CYP1A2, CYP2D6, CYP3A4, CYP2E1, CYP2B6, CYP2C9, and CYP2C8) and low level inhibition (decrease of 34.6% of baseline activity) of CYP 2C19 and three mammalian serine proteases (decreases of 17.8%, 9.3% and 7.6% of baseline activity for chymotrypsin, trypsin and thrombin A, respectively). In addition, **21** at 30 µM had no noteworthy binding to the hERG channel, decreasing binding of the tracer by 8.34%. The results suggest favorable antibacterial selectivity, low potential drug-drug interactions, and minimal cardiac channel-mediated toxicity.

**Table 3.** Safety and selectivity of boro-PBPi 21.

| **In vitro ADMET assay** | **Boro-PBPi 21 results** |
| --- | --- |
| **Solubility (formulated)** | 75 mg/mL in PBS adjusted to pH 6.5 - 7 |
| **Plasma protein binding (% bound)** | 39% (human), 19% (mouse) |
| **Hemolysis** | >1,000 µg/mL |
| **Cytotoxicity** | CC<sub>50</sub> >256 µg/mL (HeLa, MRC-5, 3T3) |
| **Mitochondrial toxicity** | No toxicity observed at 100 µM |
| **CYP inhibition** | IC<sub>50</sub> >30 µM for 8 CYP enzymes<br>(~35% inhibition of CYP2C19 observed) |
| **Micronucleus assay (mammalian chromosomal aberration assay)** | No micronucleus observed up to 300 µM |
| **hERG cardiac channel assay** | No inhibition at 30 µM |
| **Protease inhibition** | Negligible inhibition of 3 human proteases at 30 µM<br>(18% inhibition for chymotrypsin observed) |

A small-molecule therapeutic agent to treat *N. gonorrhoeae* infections is recommended to be oral or intramuscular administration within the outpatient setting^59^. Several boro-PBPi were evaluated in the Caco-2 monolayer assay to measure the potential for human gastrointestinal tract permeability^60^. None of the boro-PBPi tested, including several ester prodrugs, demonstrated sufficient passage across Caco-2 cell monolayers, indicating that oral administration of these boro-PBPi is impractical (data not shown). We therefore evaluated the potential for IM administration of **21** in male Sprague Dawley rats (**Figure S12**). Based on the pharmacokinetic results, the IM bioavailability (F_IM_) of **21** was estimated to be at least 66%, supporting the potential for IM administration of **21** for the treatment of gonorrhea.

## DISCUSSION

Boron-containing compounds have become increasingly attractive in drug discovery because of their unique binding properties to biological targets, their favorable pharmacokinetic properties, and low overall toxicity^61^. Here, we describe the identification of **21** (VNRX-14079), a novel non-β-lactam benzoxaborinine PBPi, that binds to both non-mosaic and mosaic PBP2 and exhibits potent standalone in vitro antibacterial activity and in vivo efficacy against CRO-resistant *N. gonorrhoeae*, as well as a favorable safety, antibacterial selectivity, and pharmacokinetic profile for intramuscular administration.

During the discovery phase of this program, we first identified **2**, a prototype boro-PBPi containing the ureido side chain of cefoperazone. Notably, **2** showed weaker antibacterial activity than cefoperazone against multiple *N. gonorrhoeae* strains. This reduced activity may be a consequence of the reversible nature of the covalent bond between **2** and the catalytic Ser310 of PBP2, whereas cefoperazone forms an irreversible acyl-enzyme complex with PBP2 that requires a separate hydrolysis step to regenerate the enzyme. We subsequently identified benzoate-containing **12** and aryl phosphonate-containing **15** as well as **18** with increased interactions to PBP2 and antibacterial activity. The preference of the phosphonate-containing boro-PBPi is partly related to the ability of the terminal phosphonate residue to interact with PBP2 residues Arg502, His514 and Tyr543, further stabilizing the boro-PBPi/PBP2 complex relative to **2**. X-ray crystallographic analysis of boro-PBPi/tPBP2 complexes indicates that, following binding of the benzoxaborinines **12** and **15,** the β_3_-β_4_ loop occupies an inward conformation closer to the active site of mosaic tPBP2^35/02^. Inward movement of the β_3_-β_4_ loop is postulated to promote efficient acylation of β-lactams to PBP2^48, 62^. The importance of β_3_-β_4_ loop movement for potent boro-PBPi activity is also supported by the observation that Ala501 mutations — known to cause ordering of the β_3_-β_4_ loop^54^ — reduced boro-PBPi activity in the WHO L (A501V mutation) and F89 (A501P mutation) strains. Importantly, antibacterial activities of these ureido boro-PBPi were less impacted by altered PBP2 variants (e.g., mosaic PBP2) than oxyimino-acetamide boro-PBPi (e.g., compound **1**), which is consistent with the observation that ureidopenicillins and ureidocephalosporins are more potent inhibitors of mosaic PBP2 than the oxyimino-acetamide cephalosporin CRO^48^.

Although the acquisition of an Ala501 mutation in PBP2 (i.e., A501P in F89) might lead to boro-PBPi resistance, recent studies have shown that the A501P mutation is associated with reduced biological fitness in vitro and in vivo^57^. Not surprisingly, recurrence of the A501P mutation of PBP2 has not been reported for over a decade. By contrast, the PBP2 variant of highest concern is *penA-60*, which has been observed globally in many CRO-resistant clinical isolates. This variant is not mutated at position 501 and overall contains similar mutations as PBP2 from H041. Unlike the PBP2 A501P variant, only minor fitness defects are observed for *penA-60*-possessing strains, which have likely contributed to the rapid spread of these CRO-resistant strains^63^. The MIC of **21** against WHO Q, which carries the *penA-60* allele, was 8-fold lower than CRO (MIC of 0.06 µg/mL for **21**, relative to 0.5 µg/mL for CRO). In addition, like CRO, **21** showed a low frequency of resistance (<4.17 × 10^-9^) and rapid cell killing at 4× MIC in WHO Q. Thus, **21** is active against CRO-resistant *N. gonorrhoeae* strains carrying the *penA-60* allele, while retaining favorable drug-like properties akin to CRO.

Overall, boro-PBPi can exhibit potent standalone antibacterial activity, good safety and selectivity profiles, and excellent in vivo efficacy. **21** is a promising new agent to treat *N. gonorrhoeae* infections including CRO-resistant infections. Return on investment and other financial barriers imposes a real-world limitation to development of novel antibacterial agents such as boro-PBPi **21**. An unfortunate consequence of the current antibiotic market that prevents further advancement of highly needed new antibacterial drugs^64^. Despite the current market situation, **21** is an outstanding candidate that could replace CRO for outpatient treatment of gonorrhea.

## METHODS

### Compounds and chemical synthesis

Boro-PBPi compounds were synthesized at Venatorx Pharmaceuticals, Inc. and BioDuro-Sundia as described in Supplementary Information. Ceftriaxone and azithromycin were purchased from USP. Ciprofloxacin was purchased from Sigma. Cefoperazone was purchased from Alfa Aesar. Zoliflodacin and gepotidacin were purchased from MedChemExpress. Molecular Operating Environment (MOE, Chemical Computing Group, Montreal, Canada) was used for computational chemistry and inhibitor design.

### Bacterial strains and growth conditions

The strains used are listed in **Table S13**. FA1090, H041 (WHO X), MS11, and F89 were obtained from Dr. Ann Jerse (Uniformed Services University of the Health Sciences). The *N. gonorrhoeae* WHO reference strains (WHO F, WHO G, WHO L, WHO M, WHO K, WHO L, WHO Y, and WHO Z)^65^ were obtained from the CDC. WHO Q (G7944, NCTC 14208)^66^ was obtained from the National Collection of Type Cultures. FA19 was obtained from Dr. Robert A. Nicholas (University of North Carolina). A set of 50 CDC strains (AR bank #165-214) were obtained from the CDC, of which 6 strains were confirmed not to be *N. gonorrhoeae* and excluded from the study. All *N. gonorrhoeae* strains were grown at 37 °C in a 5% CO_2_ humidified environment.

### Minimum inhibitory concentration (MIC) assays

To determine the ability of test compounds to inhibit the growth of bacterial strains, MIC assays were performed using both liquid- and agar-based microdilution methods as described previously with modifications^52, 67, 68^. Briefly, cryo-preserved bacterial cultures of clinical strains were streaked for isolation on Chocolate Agar [72 g/L (2×) GC Agar Base (Remel #R453502) and 2% (2×) hemoglobin (Remel #R451402) was autoclaved separately at 121 °C for 20 min to sterilize. Once cooled to ∼50 °C, the 2× GC Agar Base and 2× hemoglobin solutions were combined at a 1:1 volume ratio, and 1% IsoVitaleX™ Enrichment (BD # 211876) was added to the solution] and grown for ∼24 h. Freshly grown colonies were used for inoculum preparation. In the liquid broth-based MIC assay, 2-fold serial dilutions of test compounds were conducted in a 96-well plate with a final volume of 75 μL per well at 2-fold the final desired concentration in Fastidious broth (FB, prepared according to Cartwright *et al.* with an adjustment in final pH to 7.2 ± 0.2)^67^. For preparation of assay inoculum, a direct suspension was prepared by aseptically swabbing 10-15 colonies from agar plates into culture tubes containing 2 mL of fresh sterile saline. After the dilution plates were set up, direct suspensions were then diluted in a cuvette containing sterile saline and the optical density was measured at 600 nm (OD_600_). Direct suspensions were diluted in FB to make assay inocula (∼1 × 10^6^ CFU/mL) and 75 μL of assay inocula was added to the 96-well plates prepared with compound dilutions, yielding a starting bacterial concentration at 5 × 10^5^ CFU/mL. The plates were incubated for ∼24 h. The broth MIC was read visually as the lowest concentration well that completely inhibited bacterial growth.

In the agar-based method, GC agar was used to determine MIC. After the initial strain incubation for ∼24 h, bacterial colonies were re-streaked for isolation on fresh Chocolate Agar and grown for 18–24 h for use in the assay. To prepare GC agar, GC agar base (Remel #R453502) dissolved in water was autoclaved at 121°C to sterilize, and once cooled to ∼50°C, it was supplemented with IsoVitaleX™ Enrichment (BD # 211876) and 1M filter-sterilized ferric nitrate (Fisher #S25320) to achieve 1% and 12 µM final concentrations, respectively. The determined amount of test compound stock solution was pipetted directly to a 100 × 15 mm petri dish for each desired test concentration and 20 mL of the prepared GC agar (before solidification) was added to the petri dish containing test compound. Agar plates were swirled to dissolve compound and vented for solidification in the biological safety cabinet (BSC) until dry. For preparation of assay inoculum, a direct suspension was made by aseptically swabbing several colonies from the agar plates and suspending them into 2 mL of sterile cation-adjusted Mueller Hinton broth (CAMHB). Direct suspensions were then diluted with sterile CAMHB and adjusted to an OD_600_ of 0.1 (0.5 McFarland standard equivalent) to prepare the inocula. The Steer’s replicator was used to plate up to 32 spots of 2 µL inocula on a single agar plate from a separate 32-well plate containing inocula (500 µL each). The agar plates were vented in the BSC until dry, inverted, and incubated for 24 h. The agar MIC was read as the lowest concentration that completely inhibits bacterial growth.

### Time-kill assay

The time-kill assay for *N. gonorrhoeae* was performed as described^69^ with the following modifications. The bacterial inoculum for the assays was prepared by suspending colonies of *N. gonorrhoeae* grown on Chocolate Agar for ∼24 h in sterile saline. The cell suspension was adjusted to an OD_600_ of 0.1 and was diluted 1:500 in FB (final cell density ∼1 × 10^5^ colony forming units (CFU)/mL, 90 µL each) in a 96-well plate, which was then incubated with shaking at 200 RPM for 4 h to allow the bacteria to reach log phase of growth. After the initial growth phase, 10 µL of appropriate 10× drug dilutions (4× and 16× MIC) were added to each desired well for a total volume of 100 µL. As an untreated control, 10 µL of FB was added. At time -4 h (time of inoculation), 0 h (time of compound addition), 2 h, 4 h, 6 h, and 24 h samples were taken and CFU/mL in each sample was measured. Growth curves were analyzed by plotting the log_10_CFU/mL versus time using GraphPad Prism version 10.3. 2.

### Frequency of resistance (FoR)

*N. gonorrhoeae* strains (ATCC 49226, H041, and WHO Q) were grown for ∼24 h on Chocolate Agar and colonies were suspended in phosphate buffered saline (PBS) to make a cell suspension at ∼5 × 10^8^ CFU/mL. A 0.1 mL aliquot of the cell suspension was spread on 10 GC agar plates (100 mm diameter) supplemented with 1% IsoVitaleX™, 12 µM ferric nitrate and test compounds (4× and 16× MIC) for ATCC 49226 and H041. Chocolate Agar was used for WHO Q because the strain grew poorly on GC agar. The viable cell count in each suspension was determined by plating serial 10-fold dilutions onto the corresponding agar. The plates were incubated for 24 h and the number of colonies that arose were counted visually and the spontaneous mutation frequency was calculated.

### Isolation of zoliflodacin-resistant mutants and mutation identification

Zoliflodacin-resistant mutants were isolated by selection of WHO K (MIC = 0.12 µg/mL) on GC agar supplemented with 4× MIC of zoliflodacin. Colonies arising on agar were streaked on Chocolate Agar and a single colony was tested with broth MIC to confirm elevated MIC of zoliflodacin. Short-read genome-sequencing was performed on parent and isolated mutants (GENEWIZ, South Plainfield, NJ, USA). Genome analysis from FASTQ files provided by GENEWIZ was performed using Geneious Prime version 2022.1.1 (Biomatters Inc., San Diego, CA, USA). The reads were trimmed with BBDuK Adapter/Quality Trimming Version 38.84 (Brian Bushnell), yielding ∼10 million reads. Read mapping was performed with the Geneious Mapper using the WHO K reference genome (Genbank ID: GCF_900087865.2) to give >100 mean coverage of the entire chromosome followed by Geneious SNP analysis, resulting in the identification of the GyrB(D429N) mutation present in each mutant.

### Expression and purification of *N. gonorrhoeae* PBP2

In the *N. gonorrhoeae* PBP2 inhibition assay, constructs comprising the transpeptidase (TP) domain of PBP2 from the FA19 and H041 strains (tPBP2^FA19^ and tPBP2^H041^) were used. The cloning, expression, and purification of these PBP2 constructs was described previously^46, 70^. Importantly, these TP domains of PBP2 are acylated by Bocillin™-FL at the same rate as for full-length PBP2^70^.

### Measurement of in vitro inhibition of *N. gonorrhoeae* PBP2

The inhibitory potency towards wild-type PBP2^FA19^ was assessed by determining the inhibitor concentration that was required to reduce PBP2^FA19^ binding to fluorescently labelled penicillin V, Bocillin™-FL (Thermo Fisher Scientific) by 50% (IC_50_), using a fluorescence polarization (FP) competitive equilibrium binding assay^71^. To establish assay conditions for competition binding, enzyme titration/saturation binding experiments were initially performed. Bocillin™-FL was prepared at 0.2 µM in assay buffer (50 mM Hepes-NaOH pH 8.0), 300 mM NaCl and 10% (v/v) glycerol), and saturation binding performed by mixing 40 µL of PBP2 ranging in concentrations from 0–24 µM with 40 µL of the 0.2 µM Bocillin™-FL solution, in individual wells of black 384-well microplates. Immediately upon mixing, fluorescence was monitored using a Cytation™ 3 plate reader (BioTek) with an FP cube containing polarizing filters with 485 nm excitation and 520 nm emission. The instrument gain was set to achieve minimum fluorescence values of 50–60 and each measurement was the average of 3 flashes. Fluorescence was measured continuously for up to 120 min, and the response was stabilized within 10 min, with a dose-dependence on PBP2^FA19^ concentration. For competition binding assays, two-fold serial dilutions of compounds were prepared in assay buffer and mixed with Bocillin™-FL (final concentration of 0.1 µM) in black 384-well microplates. PBP2^FA19^ was added to achieve a final concentration of 0.25 µM and fluorescence was immediately measured at 1-min intervals for up to 10 min. Fluorescence values at the reaction endpoint (typically within 8 min) were normalized to the maximal response, then plotted against inhibitor concentration and the data fit to a four-parameter inhibitor-response curve to derive the half-maximal inhibitory concentration, IC_50_.

The interaction between PBP2^H041^ and Bocillin™-FL did not produce an adequate response in the FP assay. Therefore, inhibitory potency towards tPBP2^H041^ was assessed by determining the concentration of compound required to inhibit the binding of Bocillin™-FL to PBP2^H041^ by 50% (IC_50_) with an SDS-PAGE-based competition binding assay. Two-fold serial dilutions of each compound were prepared in assay buffer and mixed with 1 µM tPBP2^H041^, then incubated at ambient temperature for 60 min. Bocillin™-FL (1 µM) was added, and reaction mixtures were further incubated for 60 min before resolution by SDS-PAGE using 10% NuPAGE Bis-Tris Mini Protein Gels (Invitrogen). The amount of Bocillin™-FL incorporated into tPBP2^H041^ at each inhibitor concentration was detected by fluorescence scanning of PAGE gels with an Azure 600 Imager (Azure Biosystems, Inc.) and quantitated using ImageJ software^72^. Data were normalized to the maximum fluorescence, then plotted against inhibitor concentration and fit to a four-parameter inhibitor-response curve to derive the half-maximal inhibitory concentration, IC_50_.

### X-ray crystallography

The crystal structure of the transpeptidase domain of PBP2 derived from the cephalosporin reduced susceptibility strain 35/02 of *N. gonorrhoeae* (tPBP2^35/02^) has been reported previously^47^. The protein was purified and concentrated to 13 mg/mL in Tris-HCl, pH 7.8, with 10% glycerol and 500 mM NaCl and then crystallized in the same way as reported previously. These crystals occupy the P2_1_2_1_2_1_ space group with cell dimensions *a*=50.5, *b*=61.2, and *c*=108.8 Å and one molecule in the asymmetric unit.

Complexes of tPBP2^35/02^ with *R*,*R*-**12** and *R*,*R*-**15** were generated by soaking crystals with 1 µL of 10 mM boro-PBPi dissolved in PBS for 4 h, followed by flash-freezing in liquid nitrogen. To obtain the tPBP2^35/02^/*R*,*R*-**15** complex structure, a mixture of the *S,R-* and *R,R*-diastereomers was used for soaking and only *R,R*-**15** in the electron density was observed. Diffraction data were collected at a wavelength of 1.00 Å on an Eiger-16M detector at the SER-CAT 22-ID beamline at the Advanced Photon Source in Argonne, IL, USA. 200° of data were collected in 0.25° oscillations with an exposure time of 0.2 s/frame and a crystal-to-plate distance of 200 mm. Data were processed using HKL2000^73^ and structures solved by refinement against the *apo* structure of tPBP2^35/02^. Bound inhibitors were modeled into the |F_o_|-|F_c_| difference electron density map, followed by iterative cycles of model building and refinement using COOT 0.9.8.95^74^ and PHENIX 1.18.2-3874^75^. The crystallographic data collection and model refinement statistics are found in **Table S14.**

### Pharmacokinetic analysis of boro-PBPi 18 in mice

**18** was formulated at the maximum concentration of 100 mg/mL in PBS with pH adjusted between 6 and 7 by NaOH. The pharmacokinetic study was conducted at the BioDuro-Sundia DMPK group (Jiangsu, China) using female BALB/c mice (7–9 weeks old). **18** was administered once intravenously at 3 mg/kg or subcutaneously at 10, 30, 100, 300 and 1,000 mg/kg (3 mice per group). Blood microsamples (30 µL each) were collected at the timepoints of 0.0083, 0.25, 0.5, 1, 2, 4, 8 and 24 h post-dose. The plasma of each blood sample was prepared using K_2_EDTA and centrifugation. The concentration of boro-PBPi in the plasma was measured using the LC-MS/MS system with a Waters™ Acquity Ultra Performance LC and a Sciex API 6500 controlled by Analyst Software 1.7.1. Each plasma sample was diluted in 5% trichloroacetic acid in water followed by centrifugation. The supernatant was diluted 2-fold with 50% acetonitrile and loaded onto an Avantor® ACE 5 C4 column (50 mm × 2.1 mm). The mobile phase (MP) A was 5 mM ammonium acetate and 0.05% formic acid in water and MP B was 0.1% formic acid in acetonitrile. **18** was eluted in a linear gradient from 5% to 95% MP B over 1.2 min at a 0.6 mL/min flow rate and the elution was subjected to the MS/MS analysis using multiple reaction monitoring with a positive ion mode at 611.10/429.00. Buspirone at 5 ng/mL and tolbutamide at 50 ng/mL in 5% trichloroacetic acid were used as an internal standard. The lower limit of detection for **18** was 2 ng/mL.

### Pharmacokinetic analysis of boro-PBPi 21 in mice

**21** was formulated at the maximum concentration at 30 mg/mL in PBS with pH adjusted between 6 and 7 by NaOH. The pharmacokinetic study was conducted at the BioDuro-Sundia DMPK group using CD-1 mice (7–9 weeks old). The pharmacokinetic study of **21** was conducted similarly to that of **18**. The bioanalytical method to quantify **21** was the LC-MS/MS system with a SCIEX AD system and a SCIEX Triple Quad 7500 controlled by SCIEX OS 2.0.0.45330 software. Each sample prepared from plasma was loaded on to an ACQUITY UPLC® Peptide BEH C18 column (1.7 μm, 300 Å, 50 mm × 2.1 mm). The MP A was 1% formic acid in water and MP B is 1% formic acid in acetonitrile. **21** was eluted at a 0.5 mL/min flow rate in linear gradients from 5% to 55% of MP B over 0.1 min and to 95% of MP B over 0.8 min, and the elution was subjected to the MS/MS analysis using multiple reaction monitoring with a positive ion mode at 647.4/465.0. Tolbutamide at 0.5 ng/mL in 5% trichloroacetic acid was used as an internal standard. The lower limit of detection for **21** was 2 ng/mL.

The parameters derived from non-compartmental PK analysis were the plasma concentration at time 0 following IV administration (C_0_), plasma half-life (t_1/2_), maximum plasma concentration (*C_max_*), the area under the concentration vs. time curve from t = 0 to infinity (AUC_inf_), volume of distribution at steady-state (*V_SS_*), clearance (CL), time to maximum plasma concentration (T_max_), and bioavailability from the subcutaneous dose (F_SC_). In each of these instances, variability was related to concentrations below the lower limit of quantification (BLOQ) which restricted characterization of the terminal phase in at least one mouse per group (**Table S3**).

### Efficacy assessment of boro-PBPi 18 in the FA1090 vaginal infection model

Efficacy analysis was performed by Eurofins Pharmacology Discovery Services (Taipei, Taiwan) through the NIAID preclinical services, using ovariectomized and 17β-estradiol-treated female BALB/c mice^76^. The facility and procedures have been accredited by the Association for Assessment and Accreditation of Laboratory Animal Care International and assured from the Office of Laboratory Animal Welfare. The protocol was approved by the Institutional Animal Care and Use Committee. Groups of five immunocompetent female ovariectomized BALB/c mice aged 5-6 weeks were used for the efficacy study. Animals were received from BioLASCO Taiwan and acclimated for three days. After acclimation, an ovariectomy was performed on animals aged 4 weeks. The period of surgical recovery and acclimation was at least 7 days. Animals were treated with subcutaneous injection of meloxicam (20 mg/kg) if signs of pain or distress are observed during this period. Prior to infection, animals were subcutaneously injected with estradiol solution at 0.23 mg/mouse 2 days before infection (day -2) and on the day of infection (day 0). To minimize the indigenous vaginal bacteria, animals were treated twice daily (BID) with streptomycin (1.2 mg/mouse) and vancomycin (0.6 mg/mouse) by intraperitoneal injection along with trimethoprim sulfate at 0.4 mg/mL supplied in the drinking water. These antibiotic treatments started two days prior to infection and continued daily until the end of study. On day 0, animals were inoculated intravaginally with *N. gonorrhoeae* FA1090 (ATCC 700825, 0.02 mL [2.56 × 10^6^ CFU] per mouse) under anesthesia by intraperitoneal injection of pentobarbital (80 mg/kg), followed by rinsing the vagina with 50 mM Hepes (pH 7.4, 30 μL). **18** was administered subcutaneously twice-a-day with a 12 h interval (q12h) starting at 2 h post-infection for one day (5 mice per group). At 2 or 26 h after infection, animals were sacrificed with CO_2_ asphyxiation for harvest of vaginal lavage fluid. For the vaginal infected animals, vaginal lavage was performed twice with 200 μL GC broth containing 0.05% saponin to recover vaginal bacteria. The lavage samples from each animal were pooled in a total volume of 500 μL. Bacterial burden in the lavage fluids was determined by performing 10-fold serial dilutions and plating 0.1 mL of each onto Chocolate agar plates. The colony forming unit value (CFU) per mL lavage fluid (CFU/mL) was calculated.

### Efficacy assessment of boro-PBPi 21 in the H041 vaginal infection model

In vivo efficacy of **21** was tested in female NCI BALB/c mice as described previously^51^. Using a simulation based on the mouse plasma protein binding (19%) and the PK profile in male CD-1 mice, four dosing regimens for **21** were selected to vary %*f*T>MIC from 20% to 100% in this efficacy study. Mice in diestrus or anestrus stage of estrous cycle were randomized into 6 groups and pretreated with 17-β estradiol as well as antibiotics (streptomycin and trimethoprim) to suppress the overgrowth of the commensal microbiota and to support *N. gonorrhoeae* colonization. The pretreated mice were vaginally infected with H041 (1-2 × 10^4^ CFU/mouse) two days before treating the mice with **21** via subcutaneous injection, either once or alternate dosing regimens. Control groups were given ceftriaxone (CRO, 120 mg/kg SC, TID) or the vehicle control (PBS). Vaginal mucus was quantitatively cultured for *N. gonorrhoeae* for eight consecutive days post-treatment. A portion of the swab was also inoculated onto heart infusion agar (HIA) to monitor the presence of facultative aerobic commensal microbiota. Vaginal polymorphonuclear leukocyte (PMN) influx was assessed on each culture day by cytological examination of stained vaginal smears and reported as the percentage of PMNs among 100 vaginal cells. Efficacy was measured by comparing differences in the clearance rate and average bacterial burden over 8 days following treatment with **21**, CRO, or the compound vehicle PBS.

### In vitro safety and selectivity assays for boro-PBPi 21

The assays that examined plasma protein binding, cytotoxicity, hemolysis, mitochondrial toxicity, and chromosomal aberrations (micronucleus assay) as well as several in vitro inhibition assays (CYP450s and human proteases) or binding (hERG) assays for **21** were performed using established methods and controls that behaved as expected. The detailed description of the methods and raw data are found in **Table S12**.

### Pharmacokinetic analysis of boro-PBPi 21 in Sprague Dawley rats

The PK study was performed in male Sprague Dawley (SD) rats after a single intravenous (IV) or intramuscular (IM) dose at QPS, LLC (Newark, DE). All dosing formulations of **21** were clear freshly prepared solutions. For IM dose preparation, **21** was dissolved in vehicle (45:55 vol/vol 1N NaOH:1.6× PBS) with vortex, sonication and stirring to achieve 75 mg/mL at pH ∼6.8. A 0.6 mg/mL IV dose formulation was prepared by diluting the IM formulation with 1× PBS, pH 7.4 (final pH ∼7.1) and filtered through a 0.22 µm filter (Millex®-GV). Male Sprague Dawley rats (weight ∼270 g) were administered **21** via IM and IV routes. Blood samples (0.3 – 0.4 mL) were collected via tail vein at pre-dose (0), 0.083, 0.25, 0.5, 2, 4, 8 and 24 h post-dose from IV-dosed rats (n = 3) and pre-dose (0), 0.5, 1, 2, 4, 8 and 24 h pose-dose from IM-dosed rats (n = 3). All blood samples were collected into tubes containing K_2_EDTA on wet ice until centrifuged at 4°C, for 3 min, at 6,000 RPM (∼3,800 × *g*) within 40 min of blood collection. Terminal blood samples were collected in 10 mL Vacutainer tubes and centrifuged at 4,000 RPM (∼3,300 × *g*) for 15 min at 4°C. All plasma samples were snap frozen on dry ice and stored at approximately -70°C until bioanalysis to determine the concentration of **21** in plasma. The pharmacokinetic data (**Figure S12**) were evaluated using noncompartmental analysis (Phoenix® WinNonlin®, version 8.3, Certara USA Inc., Princeton, New Jersey) to determine IM bioavailability (F_IM_) in rats. Average concentrations from three rats per dose group were included. Following 3 mg/kg IV administration of **21** in rats, terminal half-life, *R*-squared, and adjusted *R*-squared values were 0.63 h, 0.98, and 0.98, respectively, while 30 mg/kg IM administration provided t_½_ = 0.95 h with *R*^2^ = 0.93, and adjusted *R*^2^ = 0.89. The F_IM_ was calculated by dividing the dose-normalized area under the concentration-time curve (AUC) extrapolated to infinity based on the last observed concentration (AUCINF_D_obs in h*kg*ng/mL/mg) when administered IM by the AUCINF_D_obs when administered IV (1030.81/1558.40 = 66%).

## Acknowledgements

We thank Robert Nicholas, Eric Brown, and Jean-Denis Docquier for providing strains. BioDuro-Sundia, Eurofins Panlabs Discovery Services Taiwan, Ltd., and QPS were the contract research organizations that supported this work, including compound synthesis, mouse PK, DMPK, in vivo efficacy, and rat PK. We also thank Michael Barbachyn, Alice Erwin, and Su Chiang (CARB-X) for supportive discussion, and our current and former colleagues at Venatorx for preliminary data generation and helpful discussions. Research reported in this publication was supported by the National Institute of Allergy and Infectious Diseases of the National Institutes of Health under Award Number R01 AI141239 and powered by CARB-X under Research Subaward Agreement Number 4500003206. The content is solely the responsibility of the authors and does not necessarily represent the official views of the National Institutes of Health. Diffraction data were collected at Southeast Regional Collaborative Access Team (SER-CAT) 22-ID beamline at the Advanced Photon Source, Argonne National Laboratory. SER-CAT is supported by its member institutions (https://www3.ser.aps.anl.gov/contact-us#TITLE_SER_CAT_Memberships), equipment grants (S10_RR25528, S10_RR028976 and S10_OD027000) from the National Institutes of Health, and funding from the Georgia Research Alliance. This research used resources of the Advanced Photon Source, a U.S. Department of Energy (DOE) Office of Science user facility operated for the DOE Office of Science by Argonne National Laboratory under Contract No. DE-AC02-06CH11357.

## Author contributions

T.U. A.Z., B.M., L.M.A., S.A.B., C.L.C., G.H.C., A.S.D., M.E., S.G.E.H., C.L.M., G.R., A.S., K.U., F.Y., D.C.P, C.J.B., D.M.D., and S.M.C. contributed to compound synthesis and data generation of antibacterial activity, in vitro PBP binding, ADME, toxicity, pharmacokinetics, in vivo efficacy study, and/or rat PK. B.W., Z.L., M.W., Z.Z., X.Z., and H.Z. contributed to compound synthesis. T.U., S.A.B., C.M.S., S.B., and C.D. contributed structural data generation and analyses. R.T., and A.E.J. contributed to in vivo efficacy study in the H041 infection model. T.U., A.L.Z., B.M., C.L.M., D.M.D., and S.M.C. wrote the manuscript with contributions from other authors.

## Data availability

The data that support this study are included in this published article and its supplementary information or are available from the corresponding authors on reasonable request. Structural data have been deposited to the PDB with the accession codes of 9MD0 (tPBP2^35/02^-**12**) and 9MCZ (tPBP2^35/02^-**15**).

