## Supplementary information for "A new class of penicillin-binding protein inhibitors to address drug-resistant *Neisseria gonorrhoeae*"

‡These authors contributed equally.

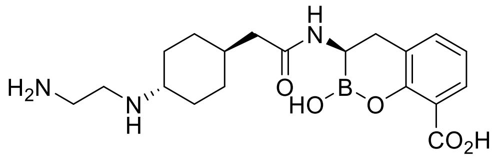

taniborbactam

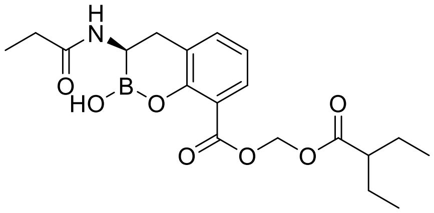

ledaborbactam etzadroxil

**Figure S1.** The chemical structures of the BLI taniborbactam and ledaborbactam etzadroxil.

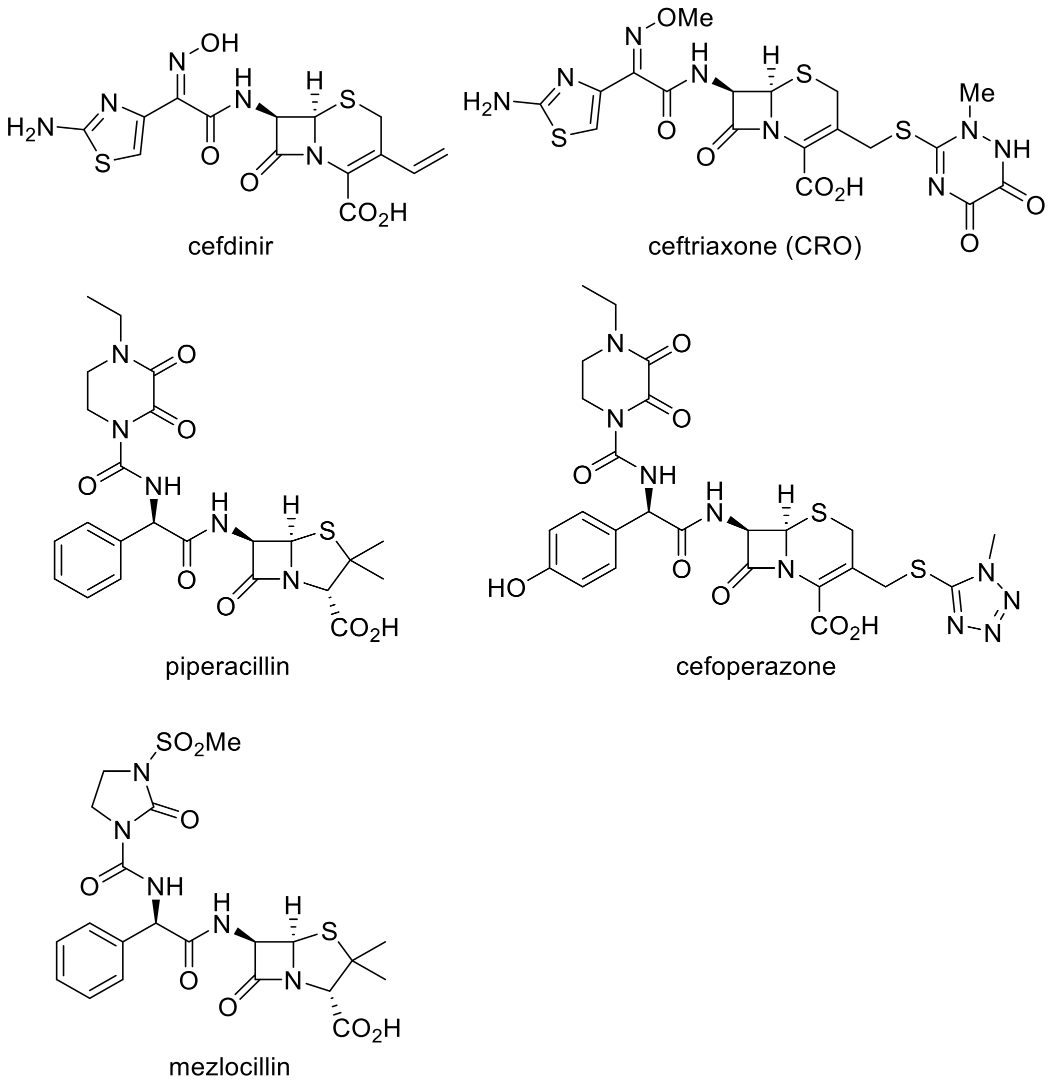

**Figure S2.** The chemical structures of cefdinir, ceftriaxone (CRO), piperacillin, cefoperazone, and mezlocillin.

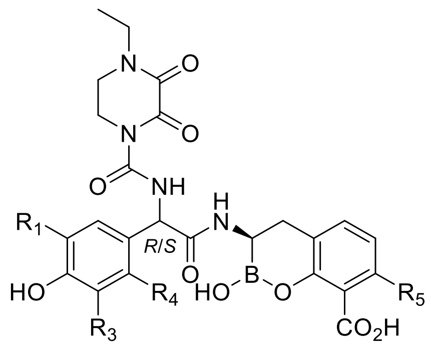

| Compound | R1 | R3 | R4 | R5 | $IC_{50}$, μM | MIC, μg/mL | |
|---|---|---|---|---|---|---|---|
| | | | | | WT *Ng* $PBP2^{WT}$ | ATCC 49226 | H041 (WHO X) |
| **cefoperazone** | NA | NA | NA | NA | 0.3 | 0.06 | 0.25 |
| **2** | H | H | H | H | 2.8 | 4 | 64 |
| *R,R*-**3** | F | H | H | H | ND | 2 | 32 |
| *R,R*-**4** | F | H | H | F | ND | 16 | 16 |
| **5** | H | H | F | H | 1.7 | 8 | 32 |
| **6** | H | F | F | H | 0.7 | 2 | 32 |
| **7** | F | F | H | H | 0.3 | 8 | 32 |
| **8** | F | H | F | H | 1.8 | 8 | 32 |
| **9** | F | F | F | H | 0.8 | 1 | 8 |

**Figure S3**. Wild type (WT) PBP2 binding and minimal inhibitory concentration (MIC) data for selected phenol-containing benzoxaborinines.

*Single diastereomer. ND, not determined. $PBP2^{WT}$, PBP2 from the FA19 strain.

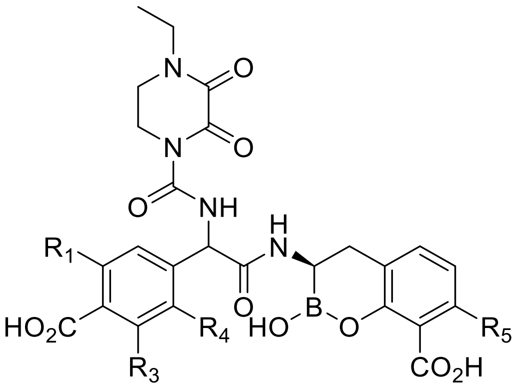

| **Compound** | **R1** | **R3** | **R4** | **R5** | **$IC_{50}$(*Ng* PBP2), µM** | | **MIC, µg/mL** | |
|---|---|---|---|---|---|---|---|---|
| | | | | | **$PBP2^{WT}$** | **$PBP2^{H041}$** | **ATCC 49226** | **H041 (WHO X)** |
| **10** | H | H | H | H | 1.2 | ND | 1 | 8 |
| **11** | H | H | H | F | ND | 395 | 4 | 8 |
| *R*,*R*-**12** | H | H | H | H | 1.0 | 42 | 0.5 | 4 |
| *R*,*R*-**13** | F | H | H | H | 1.2 | ND | 1 | 4 |
| *R*,*R*-**14** | H | F | F | H | 0.8 | 33 | 0.5 | 4 |

**Figure S4**. PBP2 binding and MIC data for selected benzoate-containing benzoxaborinines.

*Single diastereomer. ND, not determined.

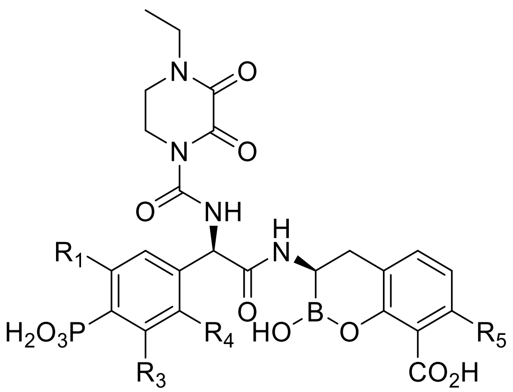

| Compound | R1 | R3 | R4 | R5 | $IC_{50}$(*Ng* PBP2), µM | | MIC, µg/mL | |
|---|---|---|---|---|---|---|---|---|
| | | | | | **PBP2$^{WT}$** | **PBP2$^{H041}$** | **ATCC 49226** | **H041 (WHO X)** |
| *R,R*-**15** | H | H | H | H | 0.7 | 7.1 | 0.06 | 0.5 |
| *R,R*-**16** | F | H | H | H | 0.3 | 2.5 | 0.06 | 0.5 |
| *R,R*-**17** | H | H | H | F | 1.4 | 10.0 | 0.12 | 0.5 |

**Figure S5**. PBP2 binding and MIC data for selected 2,3-dioxapiperidine-containing benzoxaborinines.

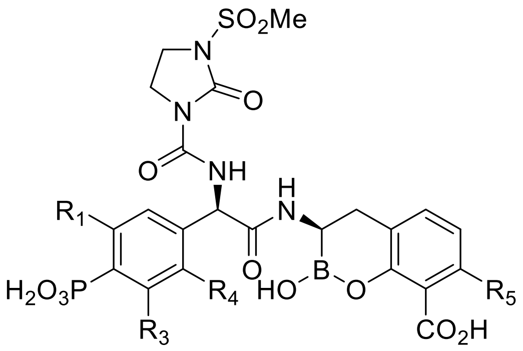

| Compound | R1 | R3 | R4 | R5 | $IC_{50}$(*Ng* PBP2), µM | | MIC, µg/mL | |
|---|---|---|---|---|---|---|---|---|
| | | | | | $PBP2^{WT}$ | $PBP2^{H041}$ | ATCC 49226 | H041 (WHO X) |
| *R,R*-**18** | H | H | H | H | 0.3 | 3.2 | 0.12 | 1 |
| *R,R*-**19** | F | H | H | H | 0.6 | 4.7 | 0.06 | 0.5 |
| *R,R*-**20** | F | F | H | H | 1.3 | 2.1 | 0.12 | 0.5 |
| *R,R*-**21** | F | H | H | F | 1.7 | 3.0 | 0.06 | 0.12 |
| *R,R*-**22** | H | F | F | H | 0.5 | 3.7 | 0.12 | 1 |
| *R,R*-**23** | H | F | F | F | 2.6 | ND | 0.06 | 0.25 |

**Figure S6**. PBP2 binding and MIC data for selected *N*-methylsulfonyl-2-imidazolinone-containing benzoxaborinines.

ND, not determined.

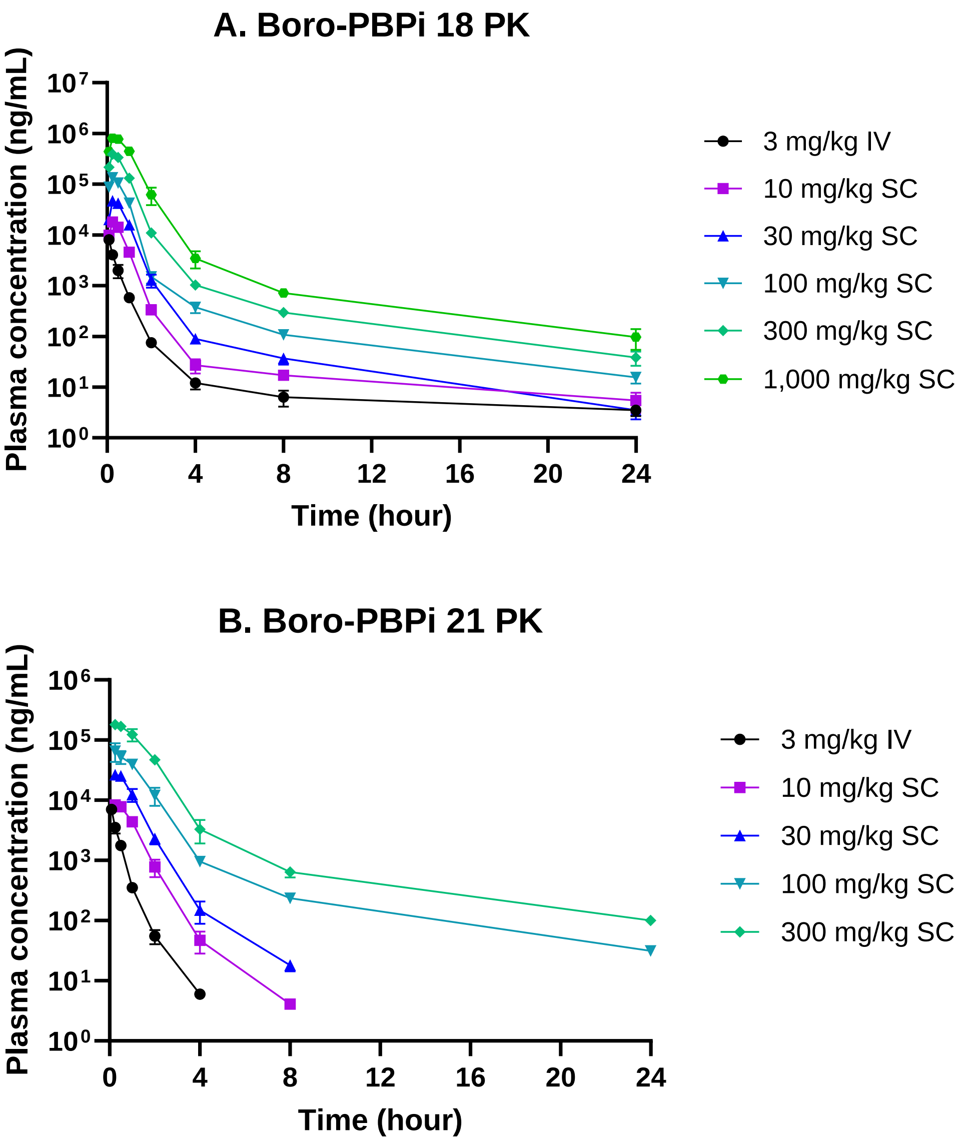

**Figure S7**. Plasma pharmacokinetics of boro-PBPi **18** (A) and **21** (B) following intravenous (IV) and subcutaneous (SC) doses in mice (n = 3/group). The error bars show standard deviations. The pharmacokinetic parameters for **18** and **21** are shown in **Table S2** and **Table S3,** respectively.

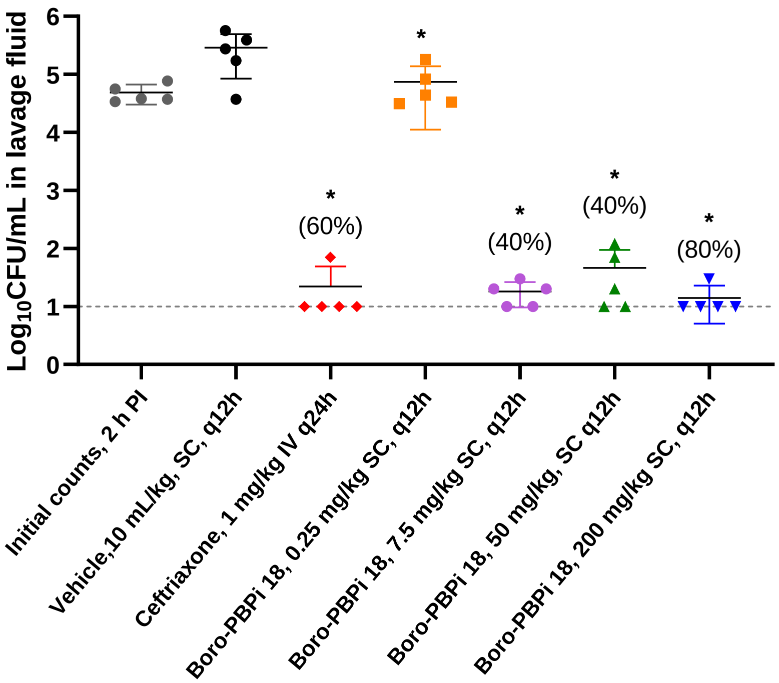

| Group | Test article description | Route, schedule | mg/kg each dose | mg/kg/day | End point | Predicted *f*T>MIC (%) |
|---|---|---|---|---|---|---|
| 1 | Baseline counts | - | - | - | 2 h | |
| 2 | Vehicle | SC, BID | - | - | 26 h | |
| 3 | Ceftriaxone | IV, QD | 1 | 1 | 26 h | |
| 4 | **18** | SC, BID | 200 | 400 | 26 h | 100 |
| 5 | **18** | SC, BID | 50 | 100 | 26 h | 100 |
| 6 | **18** | SC, BID | 7.5 | 15 | 26 h | 40 |
| 7 | **18** | SC, BID | 0.25 | 0.5 | 26 h | 13 |

**5 mice each group**
[a] Simulated concentrations remained above 4× MIC

**Figure S8.** In vivo efficacy result for boro-PBPi **18** in the murine vaginal infection model with ceftriaxone-susceptible *N. gonorrhoeae* FA1090 (ATCC 700825).

Significance was calculated relative to the CFU/mL with the vehicle at 26 h and assessed by one-way ANOVA ($F (6, 28) = 8.503$). The significant difference to the vehicle ($p < 0.05$) is marked as asterisks (*). The percent (%) shown in the figure is the percentage of animals with bacterial counts below limit of detection (LOD). The CFU/mL data are listed in **Table S4**.

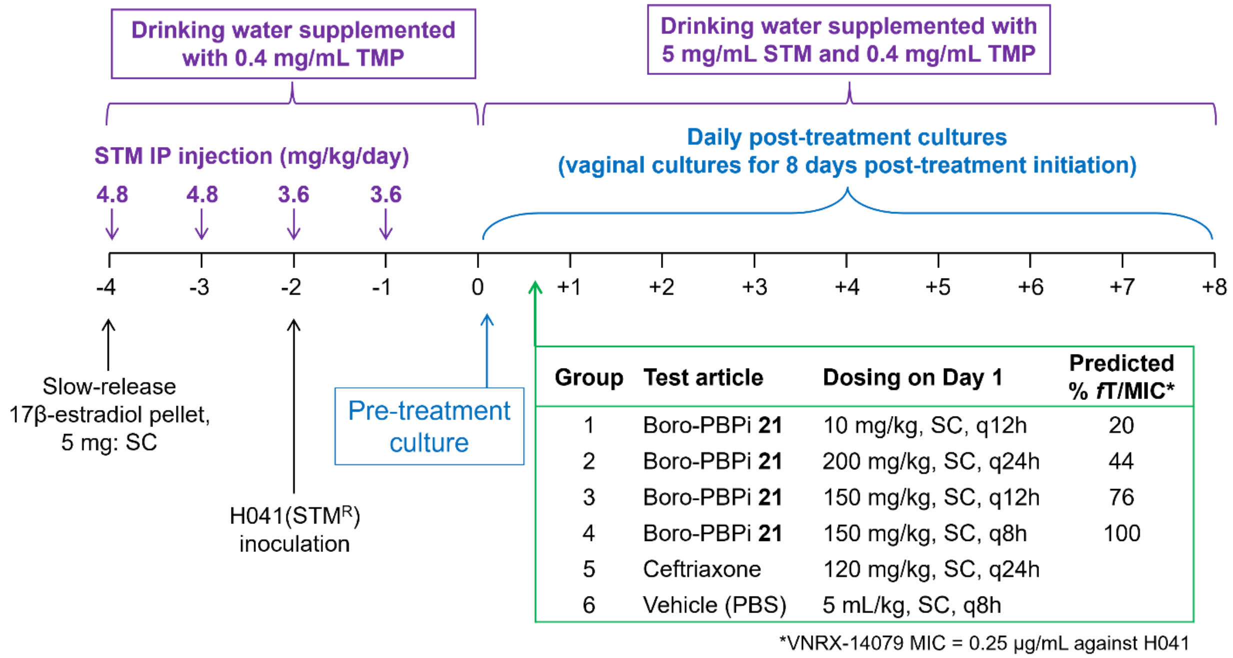

| Group | Test article description | Route, schedule | mg/kg each dose | mg/kg/day | Total dose (mg/kg) | End point | Predicted % *f*T/MIC (MIC 0.25) |
|---|---|---|---|---|---|---|---|
| 1 | **21** | SC, BID (q12h) | 10 | 20 | 20 | 1–8 days | 20 |
| 2 | **21** | SC, OD (q24h) | 200 | 200 | 200 | 1–8 days | 44 |
| 3 | **21** | SC, BID (q12h) | 150 | 300 | 300 | 1–8 days | 76 |
| 4 | **21** | SC, TID (q8h) | 150 | 450 | 450 | 1–8 days | 100 |
| 5 | Ceftriaxone | SC, TID (q8h) | 120 | 360 | 360 | 1–8 days | |
| 6 | Vehicle | SC, OD | - | - | - | 1–8 days | |
| **Total number of animals: 60 (8-10 mice each group)** | | | | | | | |

**Figure S9.** In vivo efficacy study design for boro-PBPi **21** in the murine vaginal infection model with ceftriaxone-resistant *N. gonorrhoeae* strain H041.

Abbreviations: IP, intraperitoneal; TMP, trimethoprim; SC, subcutaneous; STM, streptomycin.

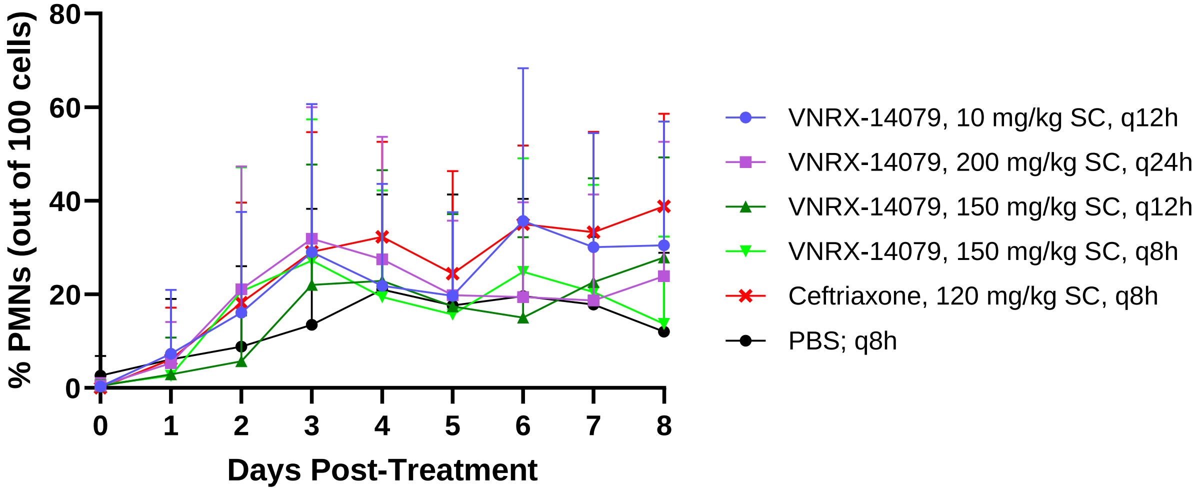

**Figure S10**. No difference in vaginal polymorphonuclear leukocytes (PMNs) over time in samples from in vivo efficacy study for boro-PBPi **21**.

PMNs were counted in samples taken from the in vivo murein efficacy studies for **21** using the H041 strains. The data were plotted above. The comparison of groups using the two-way ANOVA with repeated measures with Bonferroni post-hoc analysis did not show any significant differences (all p-values >0.05). The degrees of freedom and the F values calculated from the comparison of groups were F (5, 54) = 0.7214.

The raw data are shown in **Table S6**.

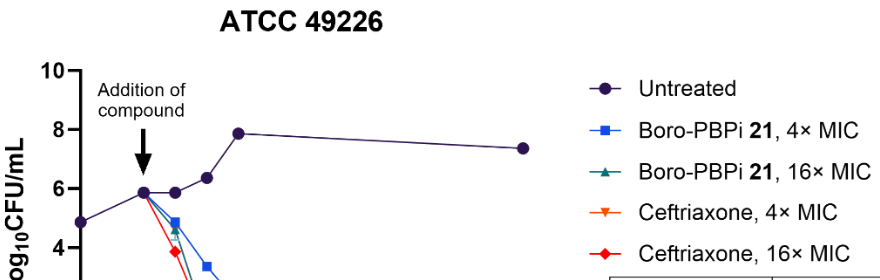

| Compound | Broth MIC (μg/mL) |
|---|---|
| Boro-PBPi **21** | 0.06 |
| Ceftriaxone | 0.004 |

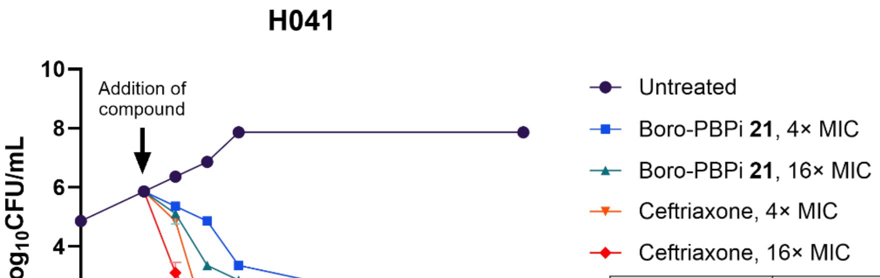

| Compound | Broth MIC (μg/mL) |
|---|---|
| Boro-PBPi **21** | 0.12 |
| Ceftriaxone | 1 |

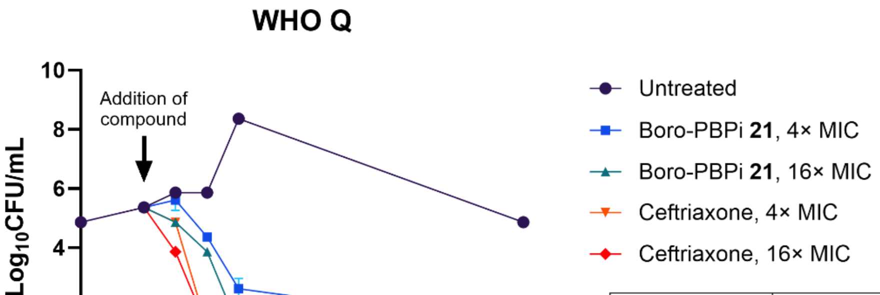

| Compound | Broth MIC (μg/mL) |
|---|---|
| Boro-PBPi **21** | 0.06 |
| Ceftriaxone | 0.25 |

**Figure S11**. In vitro bactericidal activity of boro-PBPi **21** and ceftriaxone in *N. gonorrhoeae* strains containing non-mosaic PBP2 (ATCC 49226) and mosaic PBP2 (WHO Q and H041). The MICs shown were determined when the time-kill experiments were run. LLOD: lower limit of detection.

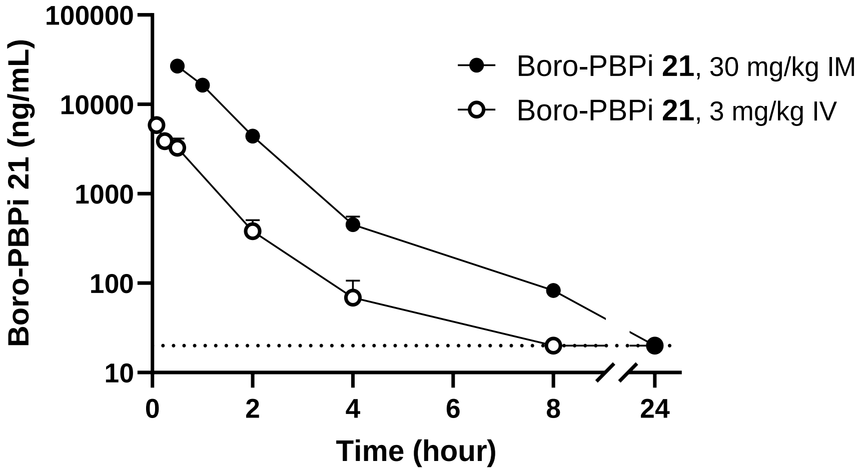

**3 mg/kg intravenous (IV) administration** (5 mL/kg dose)

| | Observed concentration (ng/mL) | | | | | | | |
|---|---|---|---|---|---|---|---|---|
| **Animal #** | **0.083 h** | **0.25 h** | **0.5 h** | **2 h** | **4 h** | **8 h** | **24 h** | **Predose** |
| 1 | 6229 | 4490 | 2742.1 | 457.7 | 70.7 | BQL | BQL | BQL |
| 2 | 5966 | 3949.5 | 2766.7 | 236.3 | 30.1 | BQL | BQL | BQL |
| 3 | 5402 | 3169.4 | 4295 | 450.2 | 106.1 | BQL | BQL | BQL |
| **Average (ng/mL)** | **5866** | **3870** | **3268** | **381** | **69** | **BQL** | **BQL** | **BQL** |

**30 mg/kg intramuscular (IM) administration** (0.4 mL/kg dose, one injection of 0.2 mL/kg each side of the left and right thigh muscle)

| | Observed concentration (ng/mL) | | | | | | |
|---|---|---|---|---|---|---|---|
| **Animal #** | **0.5 h** | **1 h** | **2 h** | **4 h** | **8 h** | **24 h** | **Predose** |
| 4 | 28035 | 17650 | 4061.9 | 377.2 | 72.5 | BQL | BQL |
| 5 | 24123 | 15463 | 4569.8 | 403.2 | 91.6 | BQL | BQL |
| 6 | 28280 | 16243 | 4589.1 | 570.8 | 84.4 | BQL | BQL |
| **Average (ng/mL)** | **26813** | **16452** | **4407** | **450** | **83** | **BQL** | **BQL** |

**Figure S12.** Plasma pharmacokinetics of boro-PBPi **21** in rats.

Dotted line: BQL, below quantitation level (concentration lower than 20 ng/mL).

The bioanalysis condition was described in **Table S15**, **Table S16**, and **Table S17**.

### CHEMISTRY EXPERIMENTAL DETAILS

**Compound 1: The preparation of *Z*-(*R*)-3-(2-(2-aminothiazol-4-yl)-2-(hydroxyimino)-acetamido)-2-hydroxy-3,4-dihydro-2H-benzo[e][1,2]oxaborinine-8-carboxylic acid:**

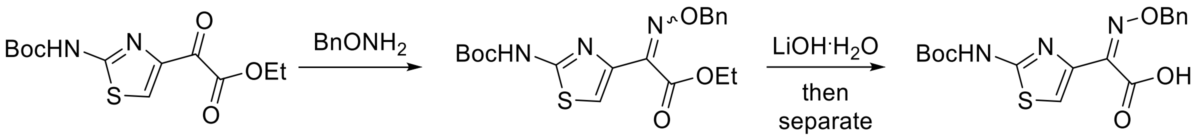

**22** **23** **24**

**Scheme S1.** The preparation of *O*-benzyl-protected oxyiminocarboxylic acid **24**.

Step 1: **Ethyl-2-((benzyloxy)imino)-2-(2-((*tert*-butoxycarbonyl)amino)thiazol-4-yl)acetate (23)**. To a solution of **22** (11 g, 36.7 mmol) in EtOH (500 mL) was added *O*-benzylhydroxylamine hydrochloride (10 g, 62.5 mmol) and the reaction mixture was stirred at ambient temperature. After 18 h, the reaction mixture was concentrated and the residue was dissolved in dichloromethane (DCM), washed with saturated aqueous $NaHCO_3$, dried over anhydrous $Na_2SO_4$, filtered and concentrated. The crude product was purified by flash silica gel chromatography (hexane/EtOAc, 20:1–2:1) to afford **23** as an inseparable mixture of *Z*- (major) and *E*- (minor) isomers, (14.7 g, 36.3 mmol, quant.). Mass spectrum, ESI-MS *m/z* 406 $(M + H)^+$, calc'd for $C_{19}H_{24}N_3O_5S$.

Step 2: **(*Z*)-2-((benzyloxy)imino)-2-(2-((*tert*-butoxycarbonyl)amino)thiazol-4-yl)acetic acid (24)**. To a solution of **23** (14.7 g, 36.3 mmol) in tetrahydrofuran (THF, 200 mL) and water (200 mL) was added LiOH hydrate (840 mg, 20 mmol) and the reaction mixture was stirred at ambient temperature. After 2 h, LC/MS analysis showed complete hydrolysis of the less-hindered *E*-isomer; the reaction mixture was diluted with water and extracted with diethyl ether. The combined ether extracts were concentrated, and the residue was dissolved in THF (150 mL), MeOH (150 mL), and water (150 mL), treated with LiOH hydrate (4.41 g, 105 mmol), and stirred at ambient temperature. After 2 days, the reaction mixture was concentrated to remove the organic solvents then acidified with 1 N HCl (pH 3). The solid was collected by filtration, washed with water, and dried *in vacuo* to yield *Z*-**24** as a tan solid (9.3 g, 70% yield). Mass spectrum, ESI-MS *m/z* 378 (M + H)$^+$, calc'd for $C_{17}H_{20}N_3O_5S$.

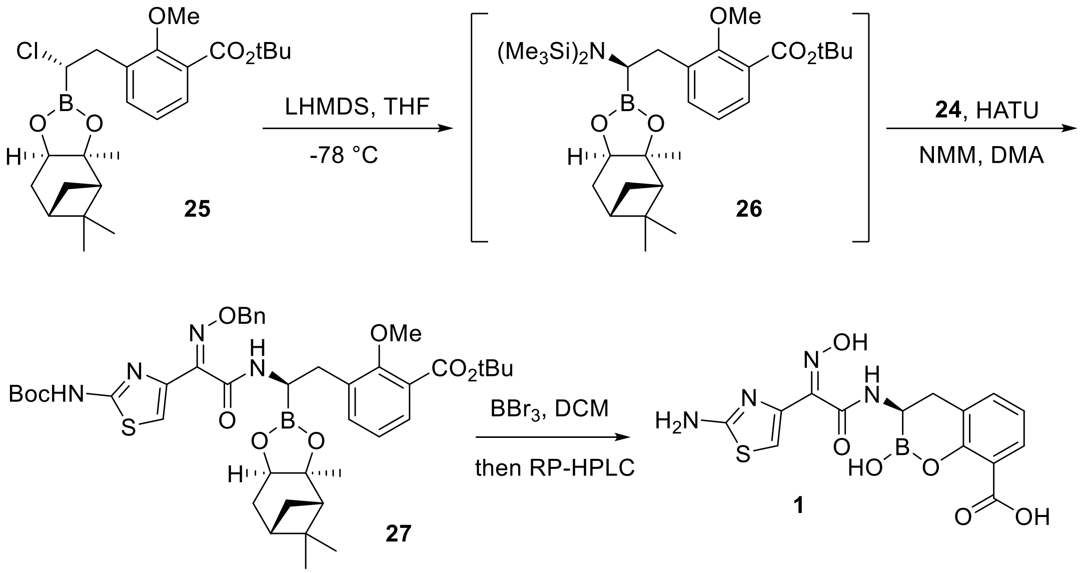

**Scheme S2.** The preparation of oxyimino-containing benzoxaborinine **1**.

Step 3: ***tert*-Butyl-3-((*R*)-2-((*Z*)-2-((benzyloxy)imino)-2-(2-((*tert*-butoxycarbonyl)amino)-thiazol-4-yl)acetamido)-2-((3a*S*,4*S*,6*S*,7a*R*)-3a,5,5-trimethylhexahydro-4,6-methanobenzo-[d][1,3,2]-dioxaborol-2-yl)ethyl)-2-methoxybenzoate (27)**. To a solution of **25** (1.35 g, 3.0 mmol) in THF (9 mL) at −78°C was added dropwise a solution of lithium hexamethyldisilazide (LHMDS, 3.0 mL, 1 M in THF, 3.0 mmol). Following the addition, the cold bath was removed, and the reaction mixture was slowly warmed to ambient temperature. After 18 h, the resulting solution of **26** (*ca.* 3 mmol) was used without further purification.

To a mixture of **24** (3.3 mmol) and HATU (3.3 mmol) in DMA (9 mL) was added NMM (3.6 mmol). After 90 min, the previously prepared solution of **26** (3.0 mmol) was added. The resulting mixture was stirred for 2.5 h, diluted with EtOAc, washed successively with water and brine, dried over anhydrous $Na_2SO_4$, filtered, and concentrated. The crude residue was purified by flash silica gel chromatography (20−100% EtOAc/hexanes) to give **27** as an off-white-colored solid (50−70% yield). Mass spectrum, ESI-MS *m/z* 789 $(M + H)^+$, calc'd for $C_{41}H_{54}BN_4O_9S$ .

Step 4: ***Z*-(*R*)-3-(2-(2-aminothiazol-4-yl)-2-(hydroxyimino)acetamido)-2-hydroxy-3,4-dihydro-2H-benzo[e][1,2]oxaborinine-8-carboxylic acid (1).** To a solution of **27** (0.4 mmol) in anhydrous DCM (15 mL) at -78°C under argon was added $BBr_3$ (1.0 M in DCM, 2.4-4 mmol) in a dropwise fashion and the reaction mixture was slowly warmed to 0°C. After 1–2 h, the reaction mixture was quenched with water (2 mL) and methanol (20 mL), concentrated to remove DCM, purified by preparative RP-HPLC using a Waters XBridge™ C18 column (5-40% acetonitrile (ACN) in water containing 0.1% TFA over 10 min; Flow rate: 45 mL/min). The product-containing fractions were lyophilized to dryness to give **1** as a white-colored solid (20–40% yield). $^1H$ NMR (MeOH-$d_4$): δ 7.89–7.78 (m, 1H), 7.40 (m, 1H), 7.04–6.83 (m, 1H), 6.13 (s, 1H), 3.45 (m, 1H), 3.04 (m, 2H) ppm. Mass spectrum, ESI-MS *m/z* 377 $(M + H)^+$, calc'd for $C_{14}H_{14}BN_4O_6S$.

**Compound 2: The preparation of (3*R*)-3-(2-(4-ethyl-2,3-dioxopiperazine-1-carboxamido)-2-(4-hydroxyphenyl)-acetamido)-2-hydroxy-3,4-dihydro-2H-benzo[e][1,2]oxaborinine-8-carboxylic acid:**

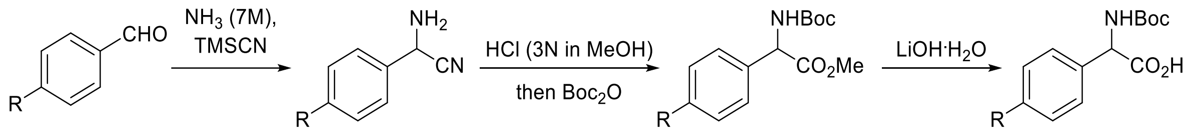

**28** R = OMe
**29** R = I
**30** R = OMe
**31** R = I
**32** R = OMe
**33** R = I
**34** R = OMe
**35** R = I

**Scheme S3.** The synthesis of racemic aryl glycine intermediates **34** and **35**.

Step 1: **2-((tert-butoxycarbonyl)amino)-2-(4-methoxyphenyl)acetic acid (34)** was prepared using the general procedures described for the preparation of *rac*-**51** (*vide infra*). Mass spectrum, ESI-MS *m/z* 282 $(M + H)^+$, calc'd for $C_{14}H_{20}NO_5$.

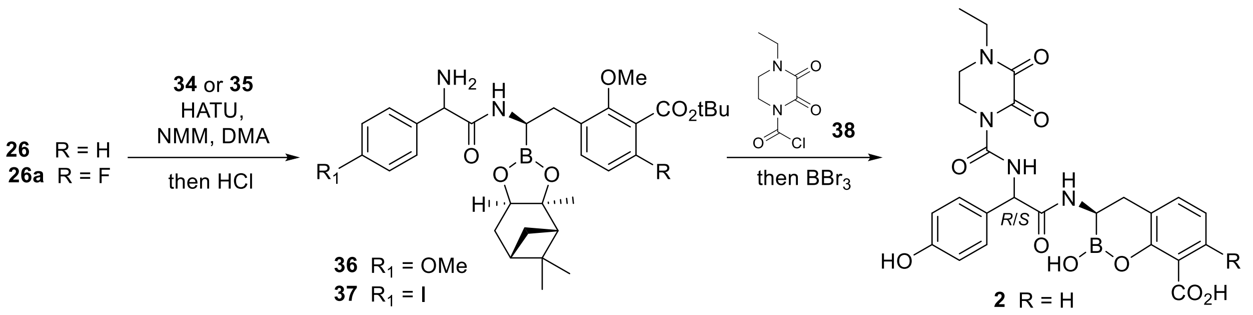

**Scheme S4.** Preparation of phenol-based boro-PBPi **2**.

Step 2: **2-(((*R*)-2-(3-(*tert*-butoxycarbonyl)-2-methoxyphenyl)-1-((3a*S*,4*S*,6*S*,7a*R*)-3a,5,5-trimethylhexa-hydro-4,6-methanobenzo[d][1,3,2]dioxaborol-2-yl)ethyl)amino)-1-(4-methoxyphenyl)-2-oxoethan-1-aminium (36)** was prepared using the general procedures described for the preparation of **43** (*vide infra*). Mass spectrum, ESI-MS *m/z* 593 $(M + H)^+$, calc'd for $C_{33}H_{46}BN_2O_7$.

Step 3: **(3*R*)-3-(2-(4-ethyl-2,3-dioxopiperazine-1-carboxamido)-2-(4-hydroxyphenyl)-acetamido)-2-hydroxy-3,4-dihydro-2H-benzo[e][1,2]oxaborinine-8-carboxylic acid (2)**. To a solution of **36** (0.5 g, 0.9 mmol) in DCM (16 mL) was cooled to 0°C. DIPEA (0.47 mL, 2.8 mmol) was added followed by **38** (0.3 g, 1.4 mmol) and the reaction mixture was warmed to ambient temperature. After 0.5 h, the reaction mixture was poured into water and extracted with DCM. The layers were separated and the organic phase was dried over anhydrous $Na_2SO_4$, filtered, and concentrated to give the ureido intermediate which was used in the next step without further purification [Mass spectrum, ESI-MS *m/z* 761 $(M + H)^+$, calc'd for $C_{40}H_{54}BN_4O_{10}$].

The ureido intermediate was treated with $BBr_3$ in DCM as described for the preparation of **1** and the final product was purified by RP-HPLC [Waters™ XBridge C18 column (5-40% ACN in $H_2O$ containing 0.1% TFA over 10 min; Flow rate: 45 mL/min)]. The product-containing fractions were combined and lyophilized to dryness to afford **2** as a white-colored solid (20–40% yield). $^1H$ NMR (MeOH-$d_4$): δ 9.42–9.20 (m, 1H), 7.82 (m, 1H), 7.38 (m, 1H), 7.05 (m, 1H), 6.80 (m, 2H),

6.50 (m, 2H), 5.30 (m, 1H), 3.98 (m, 2H), 3.60 (m, 2H), 3.50 (m, 2H), 3.32 (m, 1H), 2.98–2.80 (m, 2H), 1.20 (m, 3H) ppm. Mass spectrum, ESI-MS *m/z* 525 $(M + H)^+$, calc'd for $C_{24}H_{26}BN_4O_9$.

Boro-PBPi **3** through **9** were prepared following the general methods outlined for the synthesis of **2**.

**Compound 3: (*R*)-3-((*R*)-2-(4-ethyl-2,3-dioxopiperazine-1-carboxamido)-2-(3-fluoro-4-hydroxyphenyl)acetamido)-2-hydroxy-3,4-dihydro-2H-benzo[e][1,2]oxaborinine-8-carboxylic acid.** $^1$H NMR (MeOH-*d*$_4$): δ 9.25 (m, 1H), 7.80 (m, 1H), 7.38 (m, 1H), 7.00 (m, 1H), 6.75 (m, 2H), 6.55 (m, 1H), 5.38 (m, 1H), 3.98 (m, 2H), 3.62 (m, 2H), 3.50 (m, 2H), 3.30 (m, 1H), 2.99–2.75 (m, 2H), 1.20 (m, 3H) ppm. Mass spectrum, ESI-MS *m/z* 543 $(M + H)^+$, calc'd for $C_{24}H_{25}BFN_4O_9$.

**Compound 4: (*R*)-3-((*R*)-2-(4-ethyl-2,3-dioxopiperazine-1-carboxamido)-2-(3-fluoro-4-hydroxyphenyl)acetamido)-7-fluoro-2-hydroxy-3,4-dihydro-2H-benzo[e][1,2]oxaborinine-8-carboxylic acid.** $^1$H NMR (MeOH-*d*$_4$): δ 9.30 (m, 1H), 7.10 (m, 1H), 6.85 (m, 2H), 6.70 (m, 2H), 5.30 (m, 1H), 3.99 (m, 2H), 3.62 (m, 2H), 3.50 (m, 2H), 3.20 (s, 1H), 2.85–2.65 (m, 2H), 1.20 (m, 3H) ppm. Mass spectrum, ESI-MS *m/z* 561 $(M + H)^+$, calc'd for $C_{24}H_{24}BF_2N_4O_9$.

**Compound 5: (3*R*)-3-(2-(4-ethyl-2,3-dioxopiperazine-1-carboxamido)-2-(2-fluoro-4-hydroxyphenyl)acetamido)-2-hydroxy-3,4-dihydro-2H-benzo[e][1,2]oxaborinine-8-carboxylic acid.** $^1$H NMR (MeOH-*d*$_4$): δ 9.50–9.30 (m, 1H), 7.80 (m, 1H), 7.38 (m, 1H), 7.10–7.05 (m, 1H), 6.85–6.75 (m, 1H), 6.45–6.38 (m, 2H), 5.62 (m, 1H), 3.98 (m, 2H), 3.60 (m, 2H), 3.50 (m, 2H), 3.32 (m, 1H), 2.98–2.85 (m, 2H), 1.20 (m, 3H) ppm. Mass spectrum, ESI-MS *m/z* 543 $(M + H)^+$, calc'd for $C_{24}H_{25}BFN_4O_9$.

**Compound 6: (3*R*)-3-(2-(2,3-difluoro-4-hydroxyphenyl)-2-(4-ethyl-2,3-dioxopiperazine-1-carboxamido)acetamido)-2-hydroxy-3,4-dihydro-2H-benzo[e][1,2]oxaborinine-8-carboxylic acid.** $^1$H NMR (MeOH-$d_4$): δ 9.59–9.38 (m, 1H), 7.88–7.68 (m, 1H), 7.36–7.14 (m, 1H), 7.04–6.83 (m, 1H), 6.61–6.31 (m, 2H), 5.70–5.62 (m, 1H), 4.05–3.93 (m, 2H), 3.70–3.61 (m, 2H), 3.56–3.49 (m, 2H), 3.36 (s, 1H), 3.01–2.84 (m, 2H), 1.22–1.18 (m, 3H) ppm. Mass spectrum, ESI-MS *m/z* 561 $(M + H)^+$, calc'd for $C_{24}H_{24}BF_2N_4O_9$.

**Compound 7: (3*R*)-3-(2-(3,5-difluoro-4-hydroxyphenyl)-2-(4-ethyl-2,3-dioxopiperazine-1-carboxamido)acetamido)-2-hydroxy-3,4-dihydro-2H-benzo[e][1,2]oxaborinine-8-carboxylic acid.** $^1$H NMR (MeOH-$d_4$): δ 9.57–9.30 (m, 1H), 7.76 (dd, $J$ = 10.8, 4.5 Hz, 1H), 7.32–7.09 (m, 1H), 7.00–6.80 (m, 1H), 6.62–6.59 (m, 2H), 5.36–5.33 (m, 1H), 4.05–3.93 (m, 2H), 3.69–3.59 (m, 2H), 3.56–3.47 (m, 2H), 3.39–3.31 (m, 1H), 3.00–2.78 (m, 2H), 1.25–1.16 (m, 3H) ppm. Mass spectrum, ESI-MS *m/z* 561 $(M + H)^+$, calc'd for $C_{24}H_{24}BF_2N_4O_9$.

**Compound 8: (3*R*)-3-(2-(2,5-difluoro-4-hydroxyphenyl)-2-(4-ethyl-2,3-dioxopiperazine-1-carboxamido)acetamido)-2-hydroxy-3,4-dihydro-2H-benzo[e][1,2]oxaborinine-8-carboxylic acid.** $^1$H NMR (MeOH-$d_4$): δ 9.62–9.41 (m, 1H), 7.85–7.77 (m, 1H), 7.36–7.16 (m, 1H), 7.04–6.85 (m, 1H), 6.79–6.63 (m, 1H), 6.61–6.49 (m, 1H), 5.71–5.62 (m, 1H), 4.07–3.96 (m, 2H), 3.68–3.65 (m, 2H), 3.58–3.50 (m, 2H), 3.38–3.37 (m, 1H), 3.00 (d, $J$ = 2.8 Hz, 1H), 2.94–2.86 (m, 1H), 1.25–1.20 (m, 3H) ppm. Mass spectrum, ESI-MS *m/z* 561 $(M + H)^+$, calc'd for $C_{24}H_{24}BF_2N_4O_9$.

**Compound 9: (3*R*)-3-(2-(4-ethyl-2,3-dioxopiperazine-1-carboxamido)-2-(2,3,5-trifluoro-4-hydroxyphenyl)acetamido)-2-hydroxy-3,4-dihydro-2H-benzo[e][1,2]oxaborinine-8-carboxylic acid.** $^1$H NMR (MeOH-$d_4$): δ 9.67–9.45 (m, 1H), 7.77–7.73 (m, 1H), 7.33–7.14 (m, 1H), 7.01–6.83 (m, 1H), 6.71–6.39 (m, 1H), 5.71–5.68 (m, 1H), 4.00–3.98 (m, 2H), 3.66–3.62 (m,

2H), 3.58–3.46 (m, 2H), 3.36–3.32 (m, 1H), 3.01–2.81 (m, 2H), 1.21–1.18 (m, 3H) ppm. Mass spectrum, ESI-MS *m/z* 579 $(M + H)^+$, calc'd for $C_{24}H_{23}BF_3N_4O_9$.

**Compound 10: The preparation of (3*R*)-3-(2-(4-carboxyphenyl)-2-(4-ethyl-2,3-dioxopiperazine-1-carboxamido)-acetamido)-2-hydroxy-3,4-dihydro-2H-benzo[e][1,2]oxaborinine-8-carboxylic acid:**

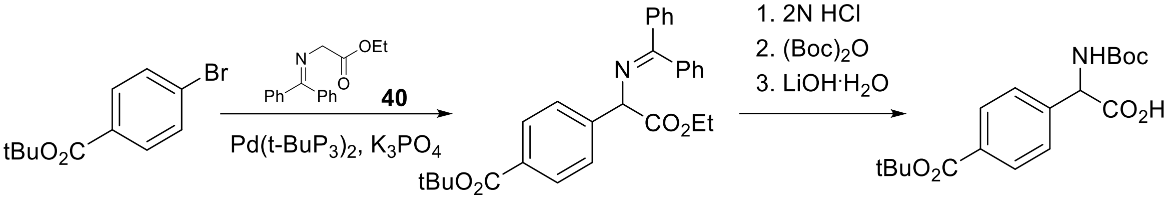

**39** **41** **42**

**Scheme S5.** The synthesis of racemic, benzoate-containing aryl glycine **42**.

Step 1: ***tert*-Butyl 4-(1-((diphenylmethylene)amino)-2-ethoxy-2-oxoethyl)benzoate (41)**. To a solution of *tert*-butyl 4-bromo-benzoate (**39**, 7 g, 27 mmol) in toluene (80 mL) was added Pd(t-$BuP_3)_2$ (2 g, 3.9 mmol), $K_3PO_4$ (17 g, 81 mmol), and ethyl *N*-(diphenylmethylene)glycinate (**40**, 10 g, 38 mmol) at ambient temperature and the reaction mixture was warmed to 145°C under an argon atmosphere. After 18 h, the reaction mixture was concentrated to dryness and the crude residue was purified by flash silica gel chromatography (EtOAc/petroleum ether, 12:1) to afford **41** (1.8 g, 15%) as a brown-colored oil. Mass spectrum, ESI-MS m/z: 444 $(M + H)^+$, calc'd for $C_{28}H_{30}NO_4$.

Step 2: **2-((*tert*-Butoxycarbonyl)amino)-2-(4-(*tert*-butoxycarbonyl)phenyl)acetic acid (42)**. A solution of **41** (2.3 g, 5 mmol) in 2N HCl in diethyl ether (20 mL) was maintained at ambient temperature. After 4 h, water (1 mL) was added. After 2 min, the reaction mixture was dried over anhydrous $Na_2SO_4$, then filtered. The filtrate was diluted with EtOAc (60 mL) then concentrated to provide the crude HCl salt (2.4 g) as a brown oil. Mass spectrum, ESI-MS *m/z* 280 $(M + H)^+$, calc'd for $C_{15}H_{22}NO_4$. To the crude HCl salt (2.4 g, 8 mmol) in THF (20 mL) was added DIPEA (3 mL, 17 mmol) and $Boc_2O$ (3.5 g, 16 mmol) and the reaction mixture was maintained at ambient temperature. After 18 h, the reaction mixture was concentrated to dryness and the crude residue was purified by flash silica gel chromatography (20:1 petroleum ether/EtOAc) to give the *N*-Boc-protected ester as a yellow-colored oil (1.6 g, 49%). Mass spectrum, ESI-MS *m/z* 380 $(M + H)^+$, calc'd for $C_{20}H_{30}NO_6$. To the *N*-Boc-protected ester (1.6 g, 4 mmol) in THF (15 mL) was added 2N LiOH (15 mL, 30 mmol) and the reaction mixture was stirred at ambient temperature. After 4 h, the reaction mixture was acidified to pH 3 with aqueous HCl, extracted with EtOAc (3 × 50 mL), dried over anhydrous $Na_2SO_4$, filtered, and concentrated. The crude acid was purified by flash silica gel chromatography (1:1 EtOAc/petroleum ether) to give **42** (1 g, 67%) as a white-colored solid. Mass spectrum, ESI-MS *m/z* 352 $(M + H)^+$, calc'd for $C_{18}H_{26}NO_6$.

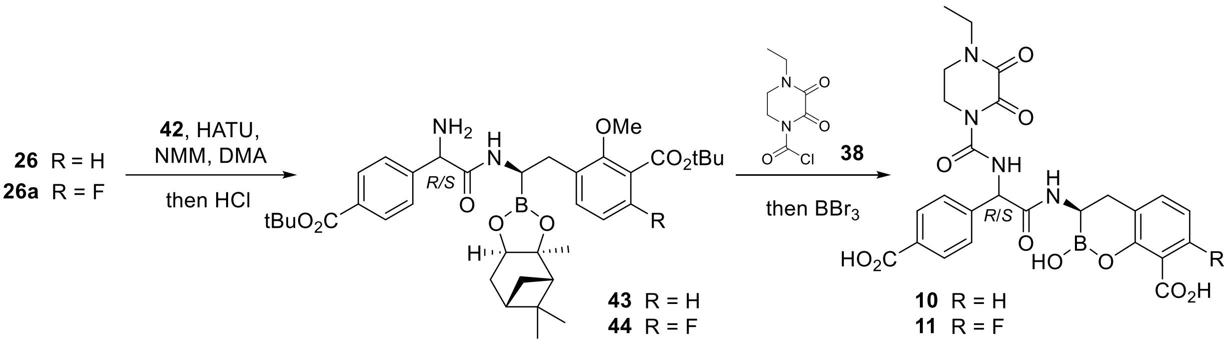

**Scheme S6.** Preparation of benzoate-based boro-PBPi **10** and **11**.

Step 3: **(3*R*)-3-(2-(4-carboxyphenyl)-2-(4-ethyl-2,3-dioxopiperazine-1-carboxamido)-acetamido)-2-hydroxy-3,4-dihydro-2H-benzo[e][1,2]oxaborinine-8-carboxylic acid (10).** Prepared from the general methods described for the preparation of **2**. $^1H$ NMR (MeOH-*d₄*): δ 9.62–9.40 (m, 1H), 7.85–7.70 (m, 3H), 7.35 (m, 1H), 7.05 (m, 2H), 6.78 (m, 1H), 5.59 (m, 1H), 4.00 (m, 2H), 3.62 (m, 2H), 3.50 (m, 2H), 3.32 (s, 1H), 2.99 -2.82 (m, 2H), 1.20 (m, 3H) ppm. Mass spectrum, ESI-MS *m/z* 553 $(M + H)^+$, calc'd for $C_{25}H_{26}BN_4O_{10}$.

Boro-PBPi **11** through **14** were prepared following the general methods outlined for the synthesis of **2**.

**Compound 11: (3*R*)-3-(2-(4-carboxyphenyl)-2-(4-ethyl-2,3-dioxopiperazine-1-carboxamido)acetamido)-7-fluoro-2-hydroxy-3,4-dihydro-2H-benzo[e][1,2]oxaborinine-8-carboxylic acid.** $^1$H NMR (MeOH-$d_4$): δ 9.70–9.42 (m, 1H), 7.99–7.82 (m, 2H), 7.25–7.10 (m, 2H), 6.75–6.62 (m, 1H), 6.30 (m, 1H), 5.58 (m, 1H), 4.00 (m, 2H), 3.62 (m, 2H), 3.50 (m, 2H), 3.20 (s, 1H), 2.80 (m, 2H), 1.20 (m, 3H) ppm. Mass spectrum, ESI-MS *m/z* 571 $(M + H)^+$, calc'd for $C_{25}H_{25}BFN_4O_{10}$.

**Compound 12: (*R*)-3-((*R*)-2-(4-carboxyphenyl)-2-(4-ethyl-2,3-dioxopiperazine-1-carboxamido)acetamido)-2-hydroxy-3,4-dihydro-2H-benzo[e][1,2]oxaborinine-8-carboxylic acid.** $^1$H NMR (MeOH-$d_4$): δ 9.69 (d, *J* = 5.5 Hz, 1H), 7.81 (d, *J* = 8.3 Hz, 2H), 7.72 (dd, *J* = 7.9, 1.2 Hz, 1H), 7.19–7.07 (m, 3H), 6.79 (t, *J* = 7.6 Hz, 1H), 5.69–5.55 (m, 1H), 4.04–4.01 (m, 2H), 3.74–3.61 (m, 2H), 3.54 (q, *J* = 7.2 Hz, 2H), 3.43–3.38 (m, 1H), 2.95–2.83 (m, 2H), 1.22 (t, *J* = 7.2 Hz, 3H) ppm. Mass spectrum, ESI-MS *m/z* 553 $(M + H)^+$, calc'd for $C_{25}H_{26}BN_4O_{10}$.

**Compound 13: (*R*)-3-((*S*)-2-(4-carboxy-3-fluorophenyl)-2-(4-ethyl-2,3-dioxopiperazine-1-carboxamido)acetamido)-2-hydroxy-3,4-dihydro-2H-benzo[e][1,2]oxaborinine-8-carboxylic acid.** $^1$H NMR (MeOH-$d_4$): δ 7.84–7.74 (m, 2H), 7.34 (d, *J* = 5.9 Hz, 1H), 7.02 (t, *J* = 7.6 Hz, 1H), 6.94 (dd, *J* = 10.9, 1.5 Hz, 1H), 6.85 (dd, *J* = 8.2, 1.4 Hz, 1H), 5.62 (s, 1H), 4.01 (t, *J* = 5.8 Hz, 2H), 3.70–3.68 (m, 2H), 3.60–3.52 (m, 3H), 3.00 (d, *J* = 3.1 Hz, 2H), 1.24 (t, *J* = 7.2 Hz, 3H) ppm. Mass spectrum, ESI-MS *m/z* 571 $(M + H)^+$, calc'd for $C_{25}H_{25}BFN_4O_{10}$.

**Compound 14: (*R*)-3-((*R*)-2-(4-carboxy-2,3-difluorophenyl)-2-(4-ethyl-2,3-dioxopiperazine-1-carboxamido)acetamido)-2-hydroxy-3,4-dihydro-2H-benzo[e][1,2]oxaborinine-8-carboxylic acid.** $^1$H NMR (MeOH-$d_4$): δ 7.76 (dd, *J* = 7.9, 1.7 Hz, 1H), 7.60–7.50 (m, 1H), 7.34

(dd, *J* = 7.3, 1.5 Hz, 1H), 7.02 (t, *J* = 7.6 Hz, 1H), 6.74 (t, *J* = 7.0 Hz, 1H), 5.90 (d, *J* = 1.3 Hz, 1H), 4.02–3.99 (m, 2H), 3.65 (t, *J* = 5.7 Hz, 2H), 3.56–3.50 (m, 2H), 3.36–3.32 (m, 1H), 2.98 (d, *J* = 3.1 Hz, 2H), 1.21 (t, *J* = 7.2 Hz, 3H) ppm. Mass spectrum, ESI-MS *m/z* 589 $(M + H)^+$, calc'd for $C_{25}H_{24}BF_2N_4O_{10}$.

**Compound 15: The Preparation of (*R*)-3-((*R*)-2-(4-Ethyl-2,3-dioxopiperazine-1-carboxamido)-2-(4-phosphonophenyl)acetamido)-2-hydroxy-3,4-dihydro-2H-benzo[e][1,2]oxaborinine-8-carboxylic acid:**

Step 1: **2-Amino-2-(4-iodophenyl)acetonitrile (31)**. To a solution of **29** (1 g, 4 mmol) in methanolic ammonia (7 N $NH_3$/methanol, 23 mL) at 0°C was added trimethylsilyl cyanide (0.75 mL, 6 mmol) and the reaction mixture was warmed to 45°C. After 7 h, the reaction mixture was concentrated to give crude **31**, which was used without further purification.

Step 2: **Methyl 2-((*tert*-butoxycarbonyl)amino)-2-(4-iodophenyl)acetate (33)**. Crude **31** (4 mmol) was dissolved in 3 N HCl in MeOH (14 mL) and warmed to 70°C. After 18 h, the reaction mixture was concentrated and used without further purification. The crude residue was suspended in THF (20 mL) and cooled at 0°C. Triethylamine (1.8 mL, 12.0 mmol) was added followed by di-*tert*-butyl dicarbonate (1.4 g, 6.0 mmol) and the reaction mixture was warmed to ambient temperature. After 1 h, the reaction mixture was concentrated to dryness. The crude product was purified by flash silica gel chromatography (10% EtOAc/hexanes) to give compound **33** as a light tan-colored solid (1.0 g, 85% yield). Mass spectrum, ESI-MS *m/z* 392 $(M + H)^+$, calc'd for $C_{14}H_{19}INO_4$.

Step 3: **2-((*tert*-Butoxycarbonyl)amino)-2-(4-iodophenyl)acetic acid (35)**. To a solution of **33** (1.1 g, 2.8 mmol) in 1:1 THF/water (20 mL) was added LiOH hydrate (0.35 g, 8 mmol) and the reaction mixture was maintained at ambient temperature. After 1 h, the reaction mixture was concentrated, adjusted to pH = 2 with 2 N HCl, and extracted with DCM. The combined organic extracts were washed with water, dried over anhydrous $Na_2SO_4$, filtered, and concentrated to give

**35** (0.9 g, 94% yield) as a white-colored solid. Mass spectrum, ESI-MS *m/z* 378 $(M + H)^+$, calc'd for $C_{13}H_{17}INO_4$.

Step 4: ***tert*-Butyl 3-((2*R*)-2-(2-amino-2-(4-iodophenyl)acetamido)-2-((3a*S*,4*S*,6*S*,7a*R*)-3a,5,5-trimethylhexahydro-4,6-methanobenzo[d][1,3,2]dioxaborol-2-yl)ethyl)-2-methoxybenzoate (43)**. To a solution of **25** (0.7 g, 1.6 mmol) in THF (6 mL) at −78°C was added dropwise a solution of lithium hexamethyldisilazide (1 M LHMDS/THF, 1.6 mL, 1.6 mmol). Following the addition, the −78°C bath was removed, and the reaction mixture was slowly warmed to ambient temperature. After 18 h, the resulting solution of **26** (*ca.* 1.6 mmol) in THF was used without further purification.

To a mixture of **42** (1.1 g, 3.1 mmol) and HATU (1.4 g, 3.6 mmol) in DMA (6 mL) was added NMM (0.44 mL, 4 mmol) at ambient temperature. After 90 min, the previously prepared solution of **26** (*ca.* 1.6 mmol) in THF was added. After 2.5 h, the reaction mixture was diluted with EtOAc, washed successively with water and brine, dried over anhydrous $Na_2SO_4$, filtered, and concentrated. The crude residue was purified by flash silica gel chromatography (30% EtOAc/hexanes) to provide the coupled product as an off-white-colored solid (0.75 g, 61% yield). Mass spectrum, ESI-MS *m/z* 789 $(M + H)^+$, calc'd for $C_{37}H_{51}BN_2O_8$.

The coupled product (0.75 g, 0.95 mmol) was added to 2 N HCl in diethyl ether (17 mL, 33 mmol) at 0°C and the reaction mixture was warmed to ambient temperature. After 18 h, the reaction mixture was concentrated to give **43** as a tan-colored solid which was used without further purification (0.69 g, quant.). Mass spectrum, ESI-MS *m/z* 689 $(M + H)^+$, calc'd for $C_{32}H_{43}BN_2O_6$.

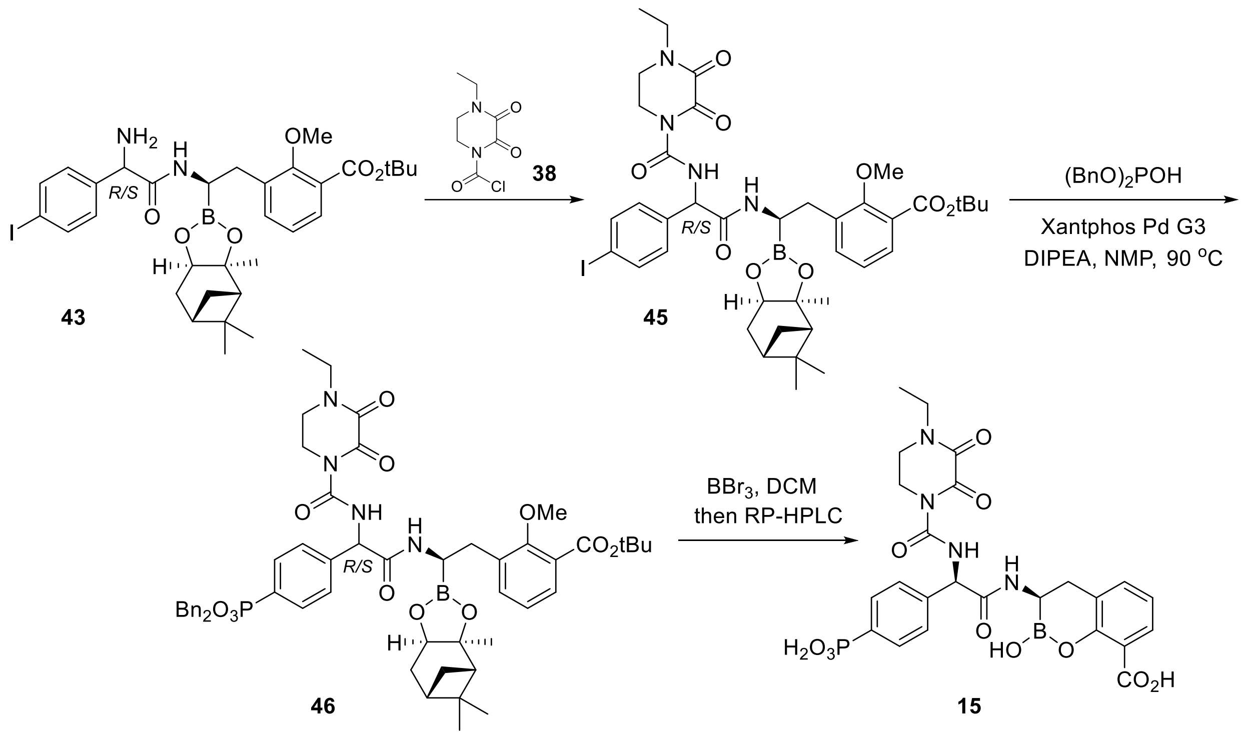

**Scheme S7.** Preparation of phosphonate-containing boro-PBPi **15**.

Step 5: ***tert*-Butyl 3-((2*R*)-2-(2-(4-ethyl-2,3-dioxopiperazine-1-carboxamido)-2-(4-iodophenyl)acetamido)-2-((3a*S*,4*S*,6*S*,7a*R*)-3a,5,5-trimethylhexahydro-4,6-methanobenzo[d]-[1,3,2]dioxaborol-2-yl)ethyl)-2-methoxybenzoate (45).** A solution of crude **43** (0.69 g, 0.9 mmol) in DCM (16 mL) was cooled to 0°C. DIPEA (0.5 mL, 2.8 mmol) was added followed by **38** (0.3 g, 1.4 mmol) and the reaction mixture was warmed to ambient temperature. After 0.5 h, the reaction mixture was poured into water and extracted with DCM. The combined organic extracts were dried over anhydrous $Na_2SO_4$, filtered, and concentrated to give ureido intermediate **45** which was used without further purification. Mass spectrum, ESI-MS *m/z*: 857 $(M + H)^+$, calc'd for $C_{39}H_{51}BIN_4O_9$.

Step 6: ***tert*-Butyl 3-((2*R*)-2-(2-(4-((dibenzyl-l$^3$-oxidaneyl)(l$^1$-oxidaneyl)phosphoryl)phenyl)-2-(4-ethyl-2,3-dioxopiperazine-1-carboxamido)acetamido)-2-((3a*S*,4*S*,6*S*,7a*R*)-3a,5,5-trimethylhexahydro-4,6-methanobenzo[d][1,3,2]dioxaborol-2-yl)ethyl)-2-methoxybenzoate**

**(46)**. To a solution of **45** (0.25 g, 0.3 mmol) in NMP (6 mL) was added DIPEA (0.15 mL, 0.95 mmol), Xantphos Pd G3 (0.03 g, 0.03 mmol), and dibenzyl phosphite (0.14 mL, 0.63 mmol). The reaction mixture was degassed (three times) under argon then warmed to 90°C. After 1 h, the reaction mixture was cooled, diluted with EtOAc, washed with water, dried over anhydrous $Na_2SO_4$, filtered, and concentrated to give **46** as an oil, which was used without further purification. Mass spectrum, ESI-MS *m/z* 991 $(M + H)^+$, calc'd for $C_{53}H_{65}BN_4O_{12}P$.

Step 7: **(*R*)-3-((*R*)-2-(4-ethyl-2,3-dioxopiperazine-1-carboxamido)-2-(4-phosphonophenyl)-acetamido)-2-hydroxy-3,4-dihydro-2H-benzo[e][1,2]oxaborinine-8-carboxylic acid (15)**. To a solution of **46** (0.28 g, 0.3 mmol) in anhydrous DCM (8 mL) at -78°C was slowly added $BBr_3$ (1.0 M in DCM, 2.8 mL, 2.8 mmol) and the reaction mixture was slowly warmed to 0°C. After 2 h, the reaction mixture was quenched with water (2 mL) and methanol (5 mL), concentrated to remove DCM and purified by preparative RP-HPLC [Waters XBridge™ C18 column (5-45% ACN in water containing 0.1% TFA over 10 min; Flow rate: 45 mL/min. Peak 1: *S*,*R*-diastereomer (undesired), RT = 1.71 min; Peak 2: *R*,*R*-diastereomer (desired), RT = 1.79 min]. The product-containing fractions were combined and lyophilized to dryness to provide **15** as a white-colored solid (30 mg, 20% yield). $^1H$ NMR (MeOH-$d_4$): δ 9.36 (d, *J* = 4.4 Hz, 1H), 7.83 (d, *J* = 8.0 Hz, 1H), 7.68 (m, 2H), 7.35 (d, *J* = 7.1 Hz, 1H), 7.08 (m, 1H), 6.96 (m, 2H), 5.56 (d, *J* = 3.5 Hz, 1H), 3.95 (m, 2H), 3.65 (m, 2H), 3.53 (m, 2H), 3.30 (m, 1H), 2.99 (s, 2H), 1.21 (t, *J* = 7.2 Hz, 3H) ppm. Mass spectrum, ESI-MS *m/z* 589 $(M + H)^+$, calc'd for $C_{24}H_{27}BN_4O_{11}P$.

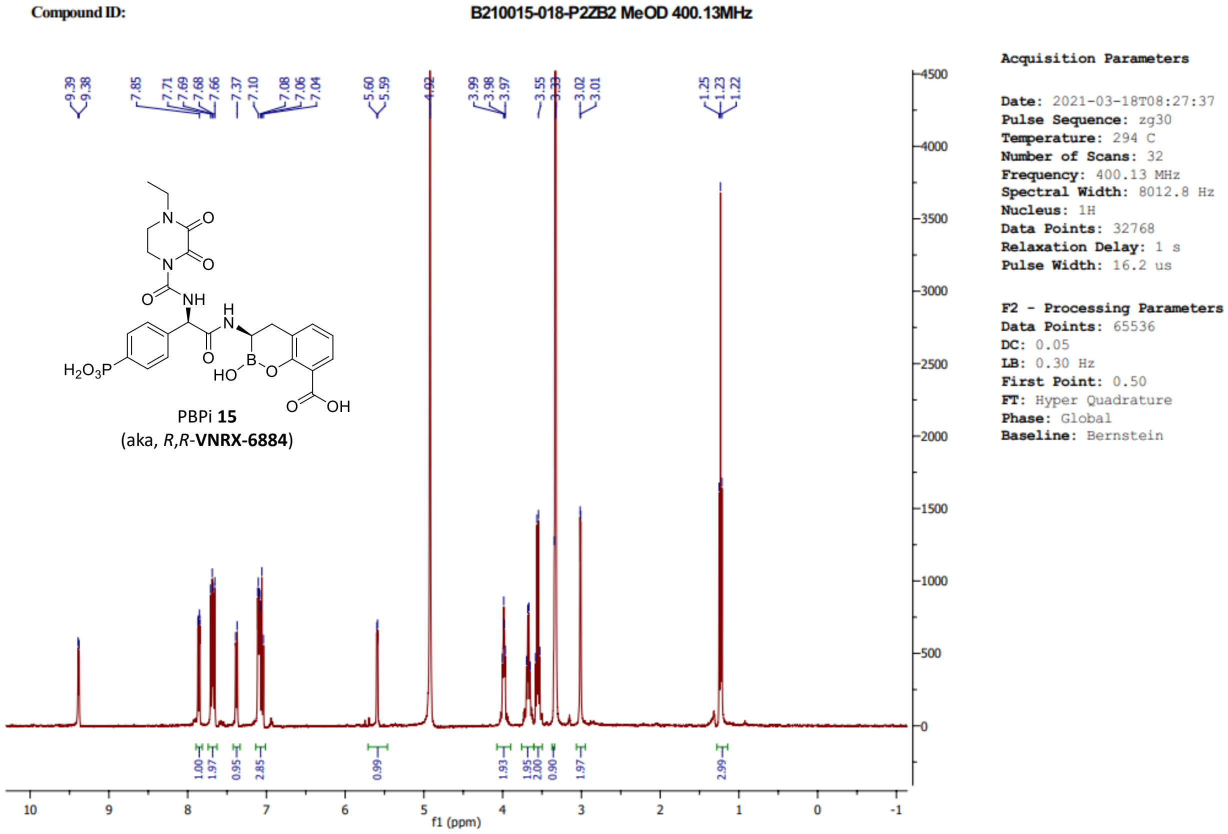

**Figure S13.** The $^{1}$H NMR spectrum of boro-PBPi **15** in MeOH-$d_4$.

```
Print of window 79: MS Spectrum
Data File  : C:\USERS\ZHOUX2\DESKTOP\光化学讲座\VNRX-6884\P2E\B210015-018-P2ZB2-51405.D
Sample Name : B210015-018-P2ZB2
=====================================================================
Acq. Operator   : sysadmin
Acq. Instrument : Agilent LCMS A                  Location : P1-A-01
Injection Date  : 3/18/2021 7:48:51 AM                 Inj :   1
                                                Inj Volume : 1.000 µl
Acq. Method     : D:\DATA\1\METHODS\P100-1000.M
Last changed    : 11/27/2019 10:24:05 AM by sysadmin
Analysis Method : C:\CHEM32\1\METHODS\DEF_LC.M
Last changed    : 11/20/2006 6:14:44 PM
```

MS Spectrum

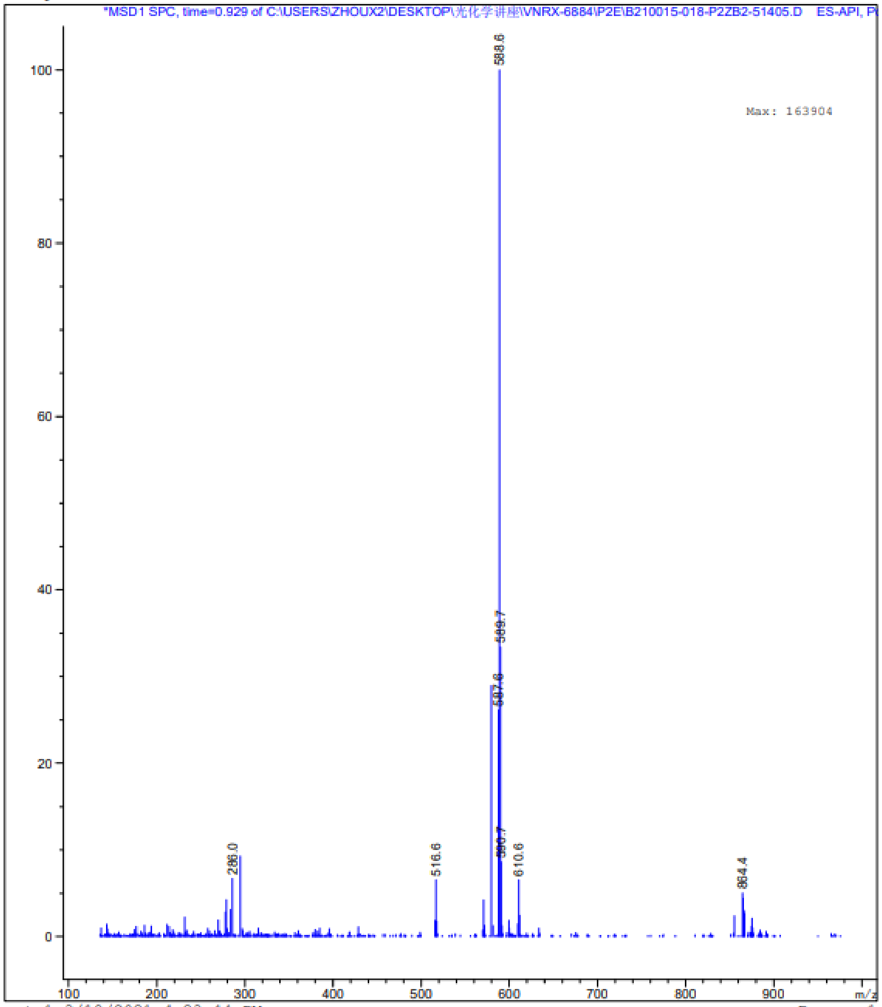

Instrument 1 3/18/2021 4:23:44 PM Page 1 of 1

**Figure S14.** Mass spectral data for boro-PBPi **15**.

Boro-PBPi **16** and **17** were prepared following the general methods outlined for the synthesis of **18**.

**Compound 16: (*R*)-3-((*R*)-2-(4-ethyl-2,3-dioxopiperazine-1-carboxamido)-2-(3-fluoro-4-phosphonophenyl)acetamido)-2-hydroxy-3,4-dihydro-2H-benzo[e][1,2]oxaborinine-8-carboxylic acid.** $^{1}$H NMR (MeOH-$d_4$): δ 9.36 (d, *J* = 4.4 Hz, 1H), 7.87 (d, *J* = 2.8 Hz, 1H), 7.65 (m, 1H), 7.35 (m, 1H), 7.08 (m, 1H), 6.96 (m, 1H), 6.75 (m, 1H), 5.56 (d, *J* = 3.5 Hz, 1H), 3.95 (m, 2H), 3.65 (m, 2H), 3.53 (m, 2H), 3.30 (m, 1H), 2.99 (s, 2H), 1.21 (t, *J* = 7.2 Hz, 3H) ppm. Mass spectrum, ESI-MS *m/z* 607 $(M + H)^+$, calc'd for $C_{24}H_{26}BFN_4O_{11}P$.

**Compound 17: (*R*)-3-((*R*)-2-(4-ethyl-2,3-dioxopiperazine-1-carboxamido)-2-(4-phosphonophenyl)-acetamido)-7-fluoro-2-hydroxy-3,4-dihydro-2H-benzo[e][1,2]oxaborinine-8-carboxylic acid.** $^{1}$H NMR (MeOH-$d_4$): δ 9.44 (m, 1H), 7.76 (m, 2H), 7.27 (m, 2H), 7.12 (m, 1H), 6.67 (m, 1H), 5.55 (m, 1H), 4.08 (m, 1H), 3.87 (m, 1H), 3.78 (m, 1H), 3.58 (m, 3H), 3.30 (m, 1H), 2.87 (s, 2H), 1.21 (t, *J* = 7.2 Hz, 3H) ppm. Mass spectrum, ESI-MS *m/z* 607 $(M + H)^+$, calc'd for $C_{24}H_{26}BFN_4O_{11}P$.

**Compound 18: The preparation of (*R*)-2-hydroxy-3-((*R*)-2-(3-(methylsulfonyl)-2-oxoimidazolidine-1-carboxamido)-2-(4-phosphonophenyl)acetamido)-3,4-dihydro-2H-benzo[e][1,2]oxaborin-ine-8-carboxylic acid:**

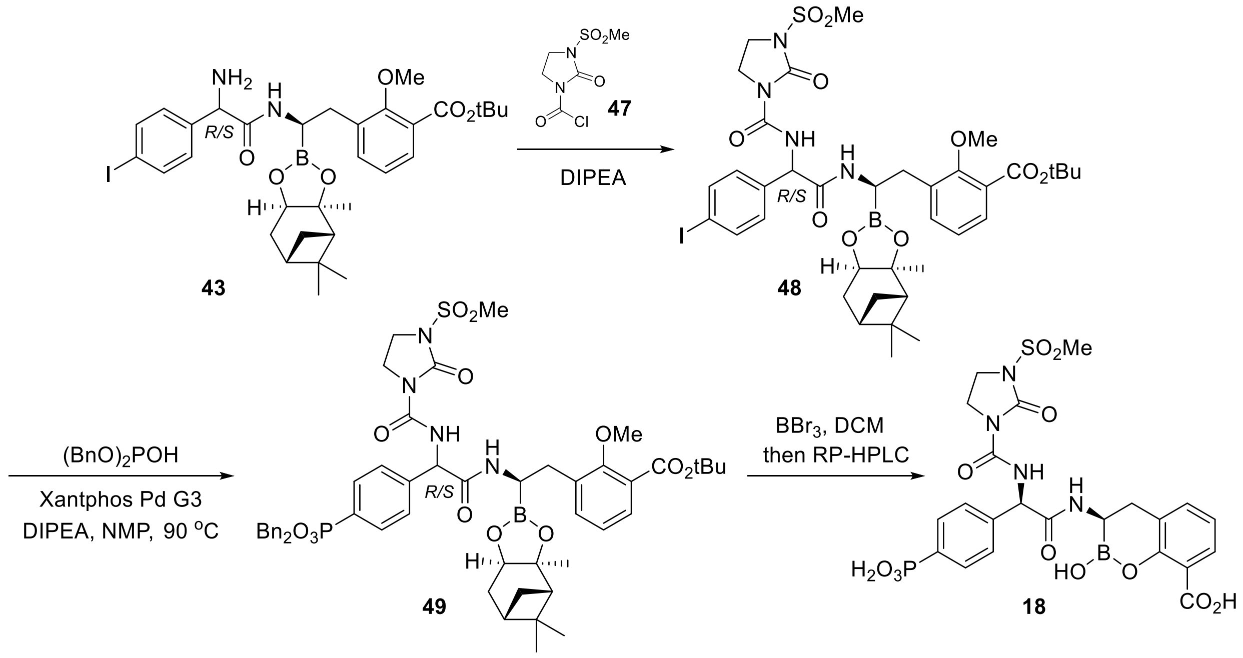

**Scheme S8.** Preparation of phosphonate-containing boro-PBPi **18**.

Step 1: ***tert*-Butyl 3-((2*R*)-2-(2-(4-iodophenyl)-2-(3-(methylsulfonyl)-2-oxoimidazolidine-1-carboxamido)acetamido)-2-((3a*S*,4*S*,6*S*,7a*R*)-3a,5,5-trimethylhexahydro-4,6-methano-benzo[d][1,3,2]dioxaborol-2-yl)ethyl)-2-methoxybenzoate (48).** Synthesized using the general method for the preparation of **15** but utilizing 3-(methylsulfonyl)-2-oxoimidazolidine-1-carbonyl chloride (**47**) in place of **38**. Crude **48** was used without further purification. Mass spectrum, ESI-MS *m/z* 879 $(M + H)^+$, calc'd for $C_{37}H_{49}BIN_4O_{10}S$.

Step 2: ***tert*-Butyl 3-((2*R*)-2-(2-(4-((dibenzyl-l$^3$-oxidaneyl)(l$^1$-oxidaneyl)phosphoryl)phenyl)-2-(3-(methylsulfonyl)-2-oxoimidazolidine-1-carboxamido)acetamido)-2-((3a*S*,4*S*,6*S*,7a*R*)-3a,5,5-trimethylhexahydro-4,6-methanobenzo[d][1,3,2]dioxaborol-2-yl)ethyl)-2-methoxybenzoate (49)**. Synthesized using the general method described for the preparation of

**15**. Crude **49** was used without further purification. Mass spectrum, ESI-MS *m/z* 1013 $(M + H)^+$, calc'd for $C_{51}H_{63}BIN_4O_{13}PS$.

Step 3: ***(R)*-2-Hydroxy-3-((*R*)-2-(3-(methylsulfonyl)-2-oxoimidazolidine-1-carboxamido)-2-(4-phosphonophenyl)acetamido)-3,4-dihydro-2H-benzo[e][1,2]oxaborinine-8-carboxylic acid (18)**. Synthesized and purified using the general methods described for the preparation of **15**. $^1$H NMR (MeOH-$d_4$): δ 8.38 (m, 1H), 7.97 (d, *J* = 10.6 Hz, 1H), 7.66 (m, 2H), 7.36 (m, 1H), 7.05 (m, 3H), 5.59 (m, 1H), 3.91 (m, 2H), 3.72 (m, 2H), 3.34 (m, 1H), 3.30 (s, 3H), 2.99 (s, 2H) ppm. Mass spectrum, ESI-MS *m/z* 611 $(M + H)^+$, calc'd for $C_{22}H_{25}BN_4O_{12}PS$.

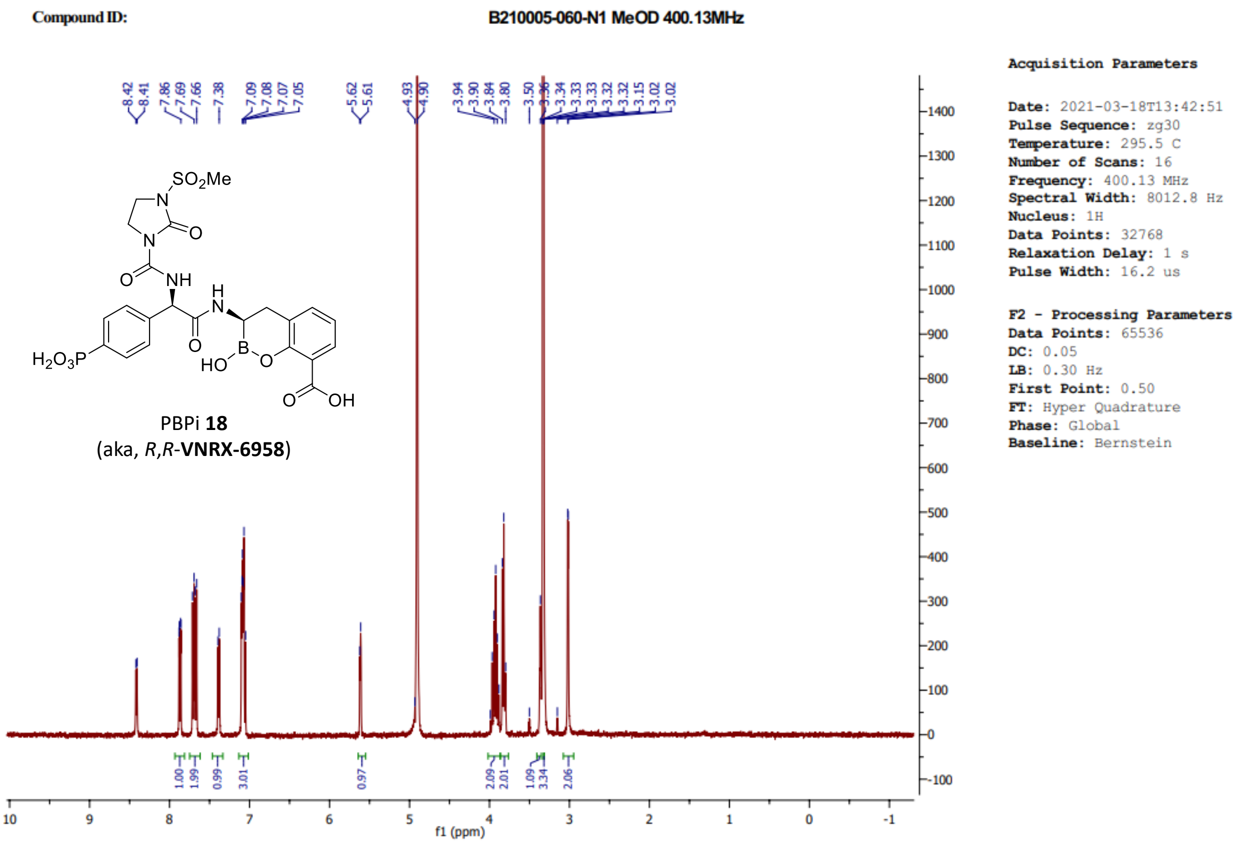

**Figure S15.** The $^1$H NMR spectrum of boro-PBPi **18** in MeOH-$d_4$.

Print of window 79: MS Spectrum
Data File : C:\USERS\ZHOUX2\DESKTOP\光化学讲座\VNRX-6958-P2\B210005-060-PP2-01878.D
Sample Name : B210005-060-PP2
==================================================================

Acq. Operator : admin
Acq. Instrument : LCMS-C Location : P1-D-07
Injection Date : 3/18/2021 1:39:08 PM Inj : 1
Inj Volume : 20.000 µl
Different Inj Volume from Sequence ! Actual Inj Volume : 1.000 µl
Acq. Method : D:\CHEMSTATION\1\METHODS\P100-1000.M
Last changed : 12/21/2020 1:00:41 PM by admin
Analysis Method : C:\CHEM32\1\METHODS\DEF_LC.M
Last changed : 11/20/2006 6:14:44 PM

MS Spectrum

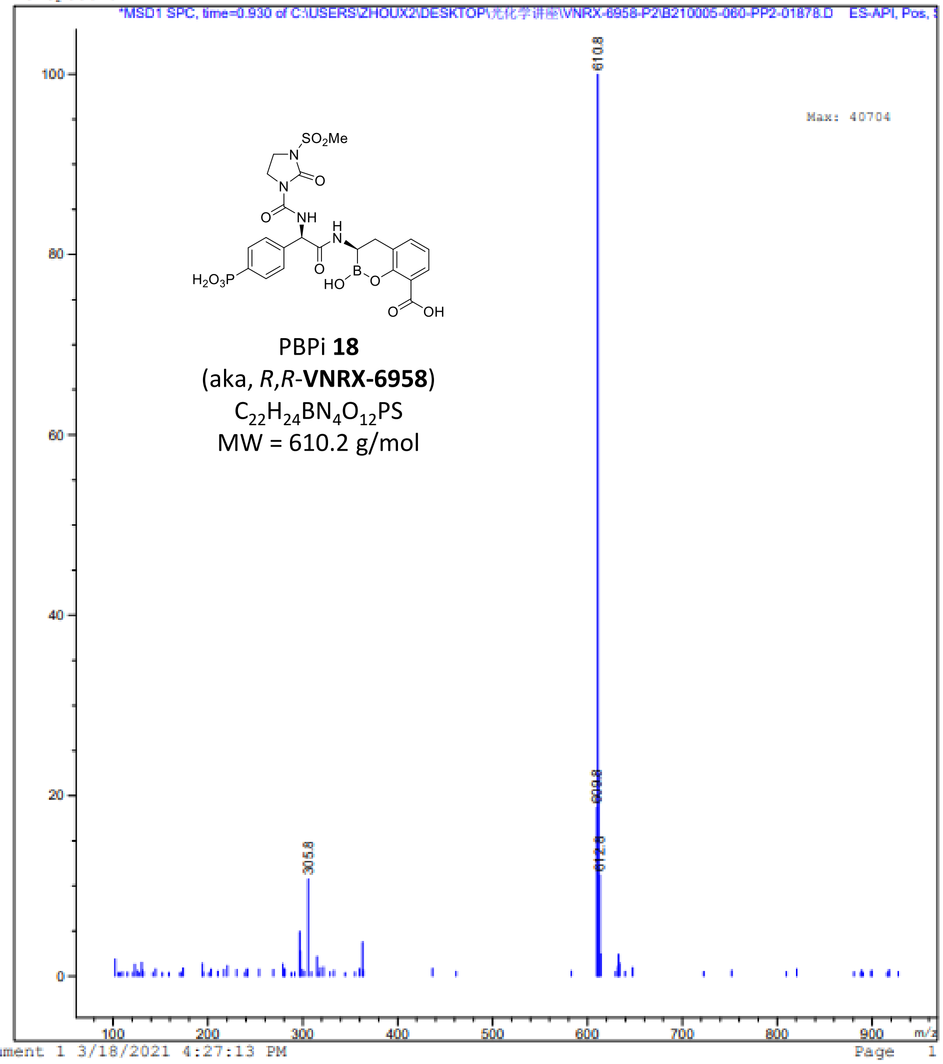

Instrument 1 3/18/2021 4:27:13 PM Page 1 of 1

**Figure S16.** Mass spectral data for boro-PBPi **18**.

**Compound 21 (VNRX-14079): The preparation of (*R*)-7-fluoro-3-((*R*)-2-(3-fluoro-4-phosphonophenyl)-2-(3-(methylsulfonyl)-2-oxoimidazolidine-1-carboxamido)acetamido)-2-hydroxy-3,4-dihydro-2H-benzo[e][1,2]oxaborinine-8-carboxylic acid:**

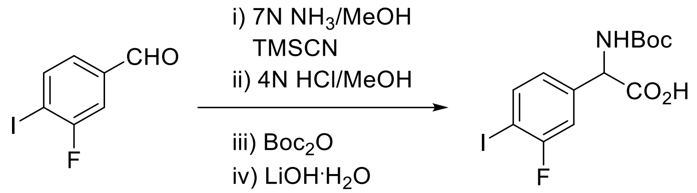

**50**
[1003709-57-6]
$C_7H_4FIO$
MW = 250.0 g/mol

*rac*-**51**
$C_{13}H_{15}FINO_4$
MW = 395.1 g/mol

**Scheme S9.** Preparation of 3-fluoro-4-iodo phenyl glycinate **51**.

**Step 1:** To 3-fluoro-4-iodobenzaldehyde (**50**, 2 g, 8 mmol) at 0°C was added 7N $NH_3$ in MeOH (43 mL) followed by trimethylsilyl cyanide (1.5 mL, 12 mmol, 1.5 eq) and the reaction mixture was warmed to 45°C. After 7 h, the reaction mixture was concentrated *in vacuo* to provide crude *rac*-2-amino-2-(3-fluoro-4-iodophenyl)acetonitrile (2.2 g, quant.) which was used in the next step without further purification. Mass spectrum, *m/z* (ESI) 277 $(M + H)^+$, calc'd for $C_8H_7FIN_2$.

**Step 2:** Crude 2-amino-2-(3-fluoro-4-iodophenyl)acetonitrile (2.2 g, 8 mmol) was dissolved in HCl/dioxane (4 N, 28 mL) and MeOH (28 mL) and warmed to 70°C. After 18 h, the homogeneous reaction mixture was concentrated to provide crude *rac*-methyl 2-amino-2-(3-fluoro-4-iodophenyl)acetate hydrogen chloride (2.8 g, quant.) which was used in the next step without further purification. Mass spectrum, *m/z* (ESI) 310 $(M + H)^+$, calc'd for $C_9H_{10}FINO_2$.

**Step 3:** Crude methyl 2-amino-2-(3-fluoro-4-iodophenyl)acetate hydrogen chloride (2.8 g, 8 mmol) was suspended in THF (30 mL) and cooled at 0°C. Triethylamine (3.3 mL, 24 mmol, 3 eq) was added followed by di-*tert*-butyl dicarbonate (2.6 g, 12 mmol, 1.5 eq) and the reaction mixture was warmed to ambient temperature. After 1 h, the reaction mixture was concentrated and the crude product was purified by flash silica gel chromatography (10% EtOAc/hexanes) to provide *rac*-methyl 2-((*tert*-butoxycarbonyl)amino)-2-(3-fluoro-4-iodophenyl)acetate (1.9 g, 57%) as an off-white-colored solid. Mass spectrum, *m/z* (ESI) 410 (M + H)$^+$, calc'd for $C_{14}H_{18}FINO_4$.

**Step 4:** To a solution of methyl 2-((*tert*-butoxycarbonyl)amino)-2-(3-fluoro-4-iodophenyl)acetate (1.9 g, 4.6 mmol) in 1:1 THF/$H_2O$ (36 mL) was added lithium hydroxide monohydrate (0.57 g, 13.6 mmol, 3 eq) at ambient temperature. After 1 h, the homogeneous solution was concentrated to remove THF. The aqueous solution was made acidic (pH 2) by the dropwise addition of 2N HCl and extracted with $CH_2Cl_2$. The organic extract was washed with water, dried over anhydrous $Na_2SO_4$, filtered, and concentrated to afford *rac*-**51** (1.8 g, quant.) as an off-white-colored solid. Mass spectrum, *m/z* (ESI) 396 (M + H)$^+$, calc'd for $C_{13}H_{16}FINO_4$.

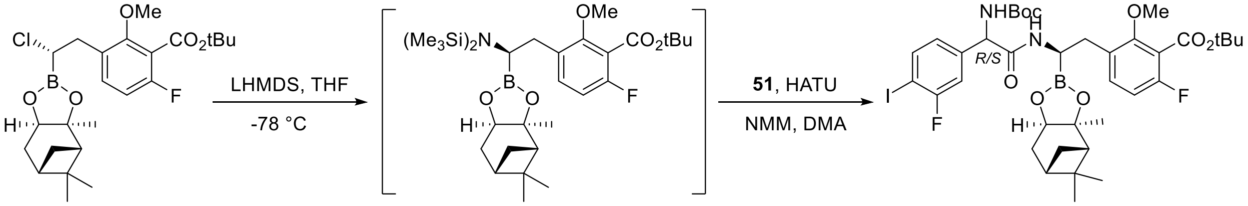

**25a** [1842401-02-8] $C_{24}H_{33}BClFO_5$ MW = 466.7 g/mol

**26a** $C_{30}H_{51}BFNO_5Si_2$ MW = 591.7 g/mol

**52** (~1:1 mixture) $C_{37}H_{48}BF_2IN_2O_8$ MW = 824.5 g/mol

**Scheme S10.** Preparation of key amide-containing intermediate **52**.

**Step 1:** To a solution of **25a** (0.52 g, 1.1 mmol) in THF (4.7 mL) at -78°C was added lithium bistrimethylsilylamide (1 M in THF, 1.2 mL, 1.2 mmol, 1.05 eq). The cold bath was removed and stirring continued at ambient temperature. After 2 h, the resultant solution of **26a** in THF was used without further purification. Mass spectrum, *m/z* (ESI) 592 (M)$^+$, calc'd for $C_{30}H_{51}BFNO_5Si_2$.

**Step 2:** To a mixture of **26a** (0.88 g, 2.22 mmol, 2 eq) and HATU (0.97 g, 2.56 mmol, 2.3 eq) was added DMA (4.7 mL) followed by *N*-methyl-morpholine (0.31 mL, 2.9 mmol, 2.6 eq). The resulting solution was stirred at ambient temperature. After 90 min, a prepared solution of **51** in THF (*ca.* 6 mL) was added and the reaction mixture was stirred at ambient temperature. After 18 h, the reaction mixture was diluted with EtOAc, washed with water, brine, dried over anhydrous $Na_2SO_4$, filtered, and concentrated. The crude product was purified by flash silica gel chromatography (30% EtOAc/hexanes) to give **52** (0.64 g, 70%) as an off-white-colored solid. Mass spectrum, *m/z* (ESI) 825 (M + H)$^+$, calc'd for $C_{37}H_{49}BF_2IN_2O_8$.

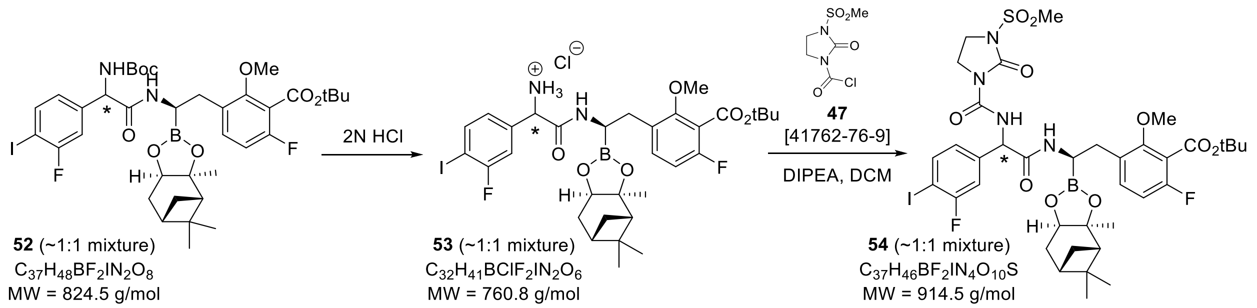

**Scheme S11.** Preparation of iodophenyl intermediate **54**.

**Step 1**: To a round-bottomed flask containing **52** (0.64 g, 0.78 mmol) at 0°C was added pre-cooled (0°C) 2N HCl in diethyl ether (14 mL) and the reaction mixture was slowly warmed to ambient temperature. After 18 h, the reaction mixture was concentrated to provide **53** (0.59 g, quant.) which was used without further purification. Mass spectrum, *m/z* (ESI) 725 (M + H)$^+$, calc'd for $C_{32}H_{41}BF_2IN_2O_6$.

**Step 2**: To a solution of **53** (0.59 g, 0.78 mmol) in $CH_2Cl_2$ (13 mL) at 0°C was added DIPEA (0.41 mL, 2.33 mmol, 3 eq) followed by 3-(methylsulfonyl)-2-oxoimidazolidine-1-carbonyl chloride (**47**) and the reaction mixture was stirred at ambient temperature. After 0.5 h, the reaction mixture was washed with water, dried over anhydrous $Na_2SO_4$, filtered, and concentrated to provide **54** (0.77 g) which was used in the next reaction without further purification. Mass spectrum, m/z (ESI) 915 (M + H)$^+$, calc'd for $C_{37}H_{47}BF_2IN_4O_{10}S$.

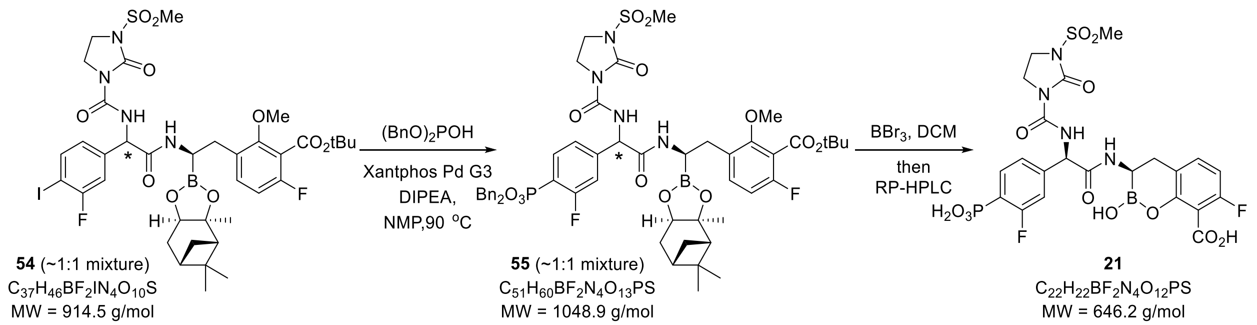

**Scheme S12.** Preparation of boro-PBPi **21**.

**Step 1**: To a solution of **54** (0.77 g, 0.84 mmol) in NMP (15 mL) was added DIPEA (0.44 mL, 2.5 mmol, 3 eq), Xantphos Pd G3 (0.08 g, 0.08 mmol, 10 mol%), and dibenzyl phosphite (0.37 mL, 1.7 mmol, 2 eq) and the reaction mixture was degassed (3×) under argon then warmed to 90°C. After 1 h, the reaction mixture was diluted with EtOAc, washed with water, brine, dried over anhydrous $Na_2SO_4$, filtered, and concentrated to afford **55** (0.74 g) which was used in the next reaction without further purification. Mass spectrum, *m/z* (ESI) 1049 $(M)^+$, calc'd for $C_{51}H_{60}BF_2N_4O_{13}PS$.

**Step 2**: To a solution of **55** (0.74 g, 0.75 mmol) in $CH_2Cl_2$ (20 mL) at -78°C was added $BBr_3$ (1M in DCM, 7.5 mL, 7.5 mmol, 10 eq) and the reaction mixture was warmed to ambient temperature. After 18 h, the reaction mixture was cooled to 0°C, quenched with a solution of water (1.5 mL) and MeOH (0.75 mL) then concentrated. The crude residue (Peak #2 for *R,R*-diastereomer) was purified by reversed-phase HPLC using a Waters™ XBridge C18 column (5-40% ACN in $H_2O$ containing 0.1% TFA over 15 min; Flow rate: 45 mL/min) followed by lyophilization to provide ***R,R*-21** (25 mg) as a white-colored solid. $^1H$ NMR (400 MHz, methanol-*d*$_4$, 27°C) δ 8.47 (d, *J* = 6.9 Hz, 1H), 7.74 (m, 1H), 7.11 (m, 1H), 7.01 (m, 2H), 6.66 (m, 1H), 5.58 (d, *J* = 6.4 Hz, 1H), 3.86 (m, 2H), 3.77 (m, 2H), 3.30 (m, 4H), 2.87 (s, 2H) ppm. Mass spectrum, *m/z* (ESI) 647 $(M + H)^+$, calc'd for $C_{22}H_{23}BF_2N_4O_{12}PS$.

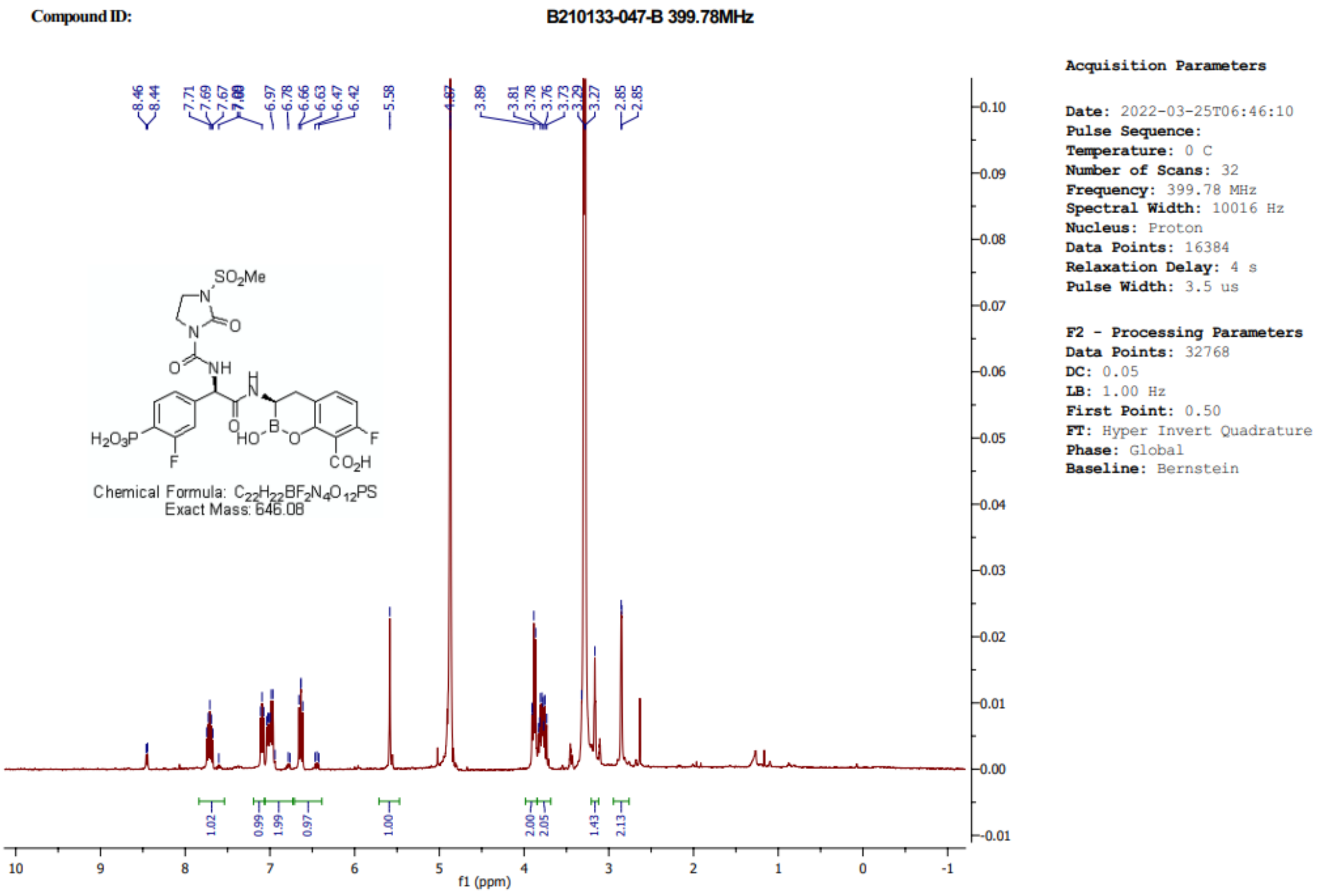

**Figure S17.** The $^{1}$H NMR spectrum of boro-PBPi **21** in MeOH-$d_4$.

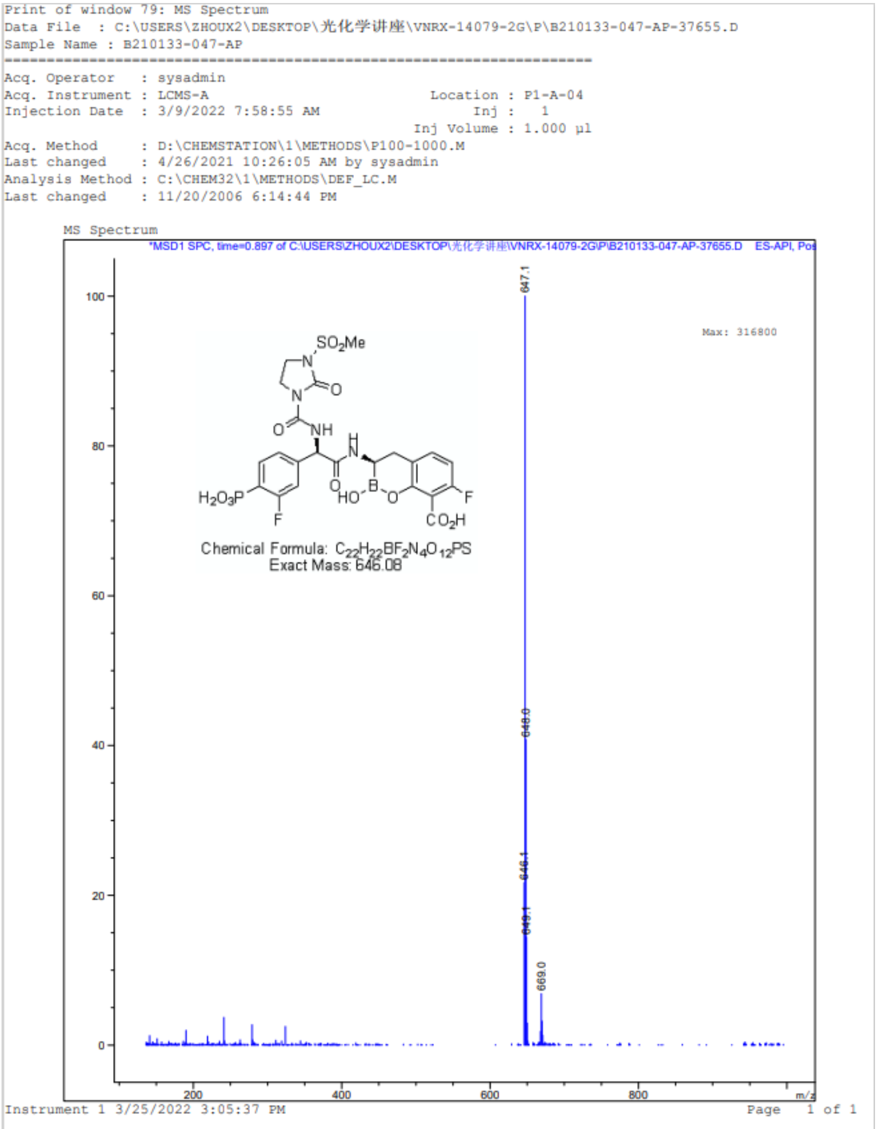

**Figure S18.** Mass spectral data for boro-PBPi **21**.

**Table S1.** Broth microdilution MIC (µg/mL) of ceftriaxone and boro-PBPi against multiple bacterial species.

| **Compound** | ***Ec* ATCC 25922 (QC strain)** | ***Ec* BAS901C* (*lptD4123*)** | ***Ec* D22* (*lpxC101*)** | ***Kp* UMM (KPC-2)** | ***Pa* ATCC 27853 (QC strain)** | ***Pa* ATCC 35151*** | ***Ab* ATCC 19606 (Type strain)** | ***Ng* ATCC 49226 (non-mosaic PBP2)** | ***Ng* H041 (mosaic PBP2)** | ***Sa* ATCC 29213 (MSSA)** |
|---|---|---|---|---|---|---|---|---|---|---|
| **Ceftriaxone** | 0.06 | ≤0.015 | 0.03 | >128 | 16 | 0.06 | 32 | 0.008 | 1 | 4 |
| **1** | 32 | 8 | 16 | 32 | >128 | 32 | >128 | 16 | >128 | 8 |
| **2** | 32 | 1 | 8 | 32 | >128 | 2 | >128 | 4 | 64 | >128 |
| **12** | 32 | 0.06 | 4 | 32 | 32 | 0.25 | 128 | 0.5 | 4 | >128 |
| **15** | 32 | 0.06 | 4 | 32 | 64 | 0.5 | 16 | 0.06 | 0.5 | >128 |
| **18** | 64 | 0.06 | 4 | 64 | >128 | 2 | 16 | 0.12 | 1 | >128 |
| **21** | >128 | 0.5 | 16 | >128 | >128 | 4 | 128 | 0.06 | 0.12 | >128 |

*Hyperpermeable strains.

The description of the bacterial strains used are listed in **Table S12**.

Abbreviations: *Ec*, *Escherichia coli*; *Kp*, *Klebsiella pneumoniae*; *Pa*, *Pseudomonas aeruginosa*; *Ab*, *Acinetobacter baumannii*; *Ng*, *Neisseria gonorrhoeae*; *Sa*, *Staphylococcus aureus*.

**Table S2.** Plasma exposure and pharmacokinetic parameters derived for boro-PBPi **18** following intravenous (IV) and subcutaneous (SC) doses in female BALB/c mice (n = 3/group).

**Boro-PBPi 18, plasma exposure (raw data)**

| Boro-PBPi 18, IV at 3 mg/kg | | | | | | |
|---|---|---|---|---|---|---|
| Time (h) | Calculated Concentration (ng/mL) | | | | | |
| | Mouse #1 | Mouse #2 | Mouse #3 | Mean | SD | CV (%) |
| 0.0833 | 7350 | 7560 | 9100 | 8003 | 956 | 11.9 |
| 0.25 | 3450 | 4200 | 4450 | 4033 | 520 | 12.9 |
| 0.5 | 1390 | 2570 | 2020 | 1993 | 590 | 29.6 |
| 1 | 489 | 633 | 602 | 575 | 76 | 13.2 |
| 2 | 69.3 | 80.4 | 74.6 | 74.8 | 5.6 | 7.43 |
| 4 | 15.1 | 9.07 | 12.0 | 12.1 | 3.0 | 25.0 |
| 8 | 5.44 | 8.79 | 4.72 | 6.32 | 2.17 | 34.4 |
| 24 | 4.08 | 2.91 | BLOQ | 2.33 | NA | NA |

| Boro-PBPi 18, SC at 10 mg/kg | | | | | | |
|---|---|---|---|---|---|---|
| Time (h) | Calculated Concentration (ng/mL) | | | | | |
| | Mouse #4 | Mouse #5 | Mouse #6 | Mean | SD | CV (%) |
| 0.0833 | 12100 | 8150 | 9080 | 9777 | 2065 | 21.1 |
| 0.25 | 20000 | 13700 | 20000 | 17900 | 3637 | 20.3 |
| 0.5 | 13700 | 12700 | 16200 | 14200 | 1803 | 12.7 |
| 1 | 4820 | 4150 | 4680 | 4550 | 353 | 7.77 |
| 2 | 340 | 293 | 364 | 332 | 36 | 10.9 |
| 4 | 21.4 | 22.8 | 36.7 | 27.0 | 8.5 | 31.4 |
| 8 | 19.4 | 17.1 | 15.1 | 17.2 | 2.2 | 12.5 |
| 24 | 6.21 | 2.82 | 7.30 | 5.44 | 2.34 | 42.9 |

| Boro-PBPi 18, SC at 30 mg/kg | | | | | | |
|---|---|---|---|---|---|---|
| Time (h) | Calculated Concentration (ng/mL) | | | | | |
| | Mouse #7 | Mouse #8 | Mouse #9 | Mean | SD | CV (%) |
| 0.0833 | 19100 | 20900 | 20300 | 20100 | 917 | 4.56 |
| 0.25 | 42800 | 45400 | 51800 | 46667 | 4632 | 9.93 |
| 0.5 | 39100 | 36400 | 49600 | 41700 | 6974 | 16.7 |
| 1 | 14100 | 14000 | 19100 | 15733 | 2916 | 18.5 |
| 2 | 1020 | 1130 | 1700 | 1283 | 365 | 28.4 |
| 4 | 85.3 | 80.7 | 101 | 89.0 | 10.6 | 12.0 |
| 8 | 29.5 | 34.3 | 46.8 | 36.9 | 8.9 | 24.2 |
| 24 | 2.87 | 2.76 | 4.85 | 3.49 | 1.18 | 33.7 |

| Boro-PBPi 18, SC at 100 mg/kg | | | | | | |
|---|---|---|---|---|---|---|
| Time (h) | Calculated Concentration (ng/mL) | | | | | |
| | Mouse #10 | Mouse #11 | Mouse #12 | Mean | SD | CV (%) |
| 0.0833 | 88300 | 89600 | 83900 | 87267 | 2987 | 3.42 |
| 0.25 | 157000 | 124000 | 126000 | 135667 | 18502 | 13.6 |
| 0.5 | 120000 | 109000 | 88600 | 105867 | 15933 | 15.0 |
| 1 | 50800 | 44400 | 33000 | 42733 | 9016 | 21.1 |
| 2 | 1370 | 1180 | 1940 | 1497 | 396 | 26.4 |
| 4 | 446 | 409 | 276 | 377 | 89 | 23.7 |
| 8 | 95.4 | 129 | 101 | 108 | 18 | 16.6 |
| 24 | 19.5 | 11.9 | 15.3 | 15.6 | 3.8 | 24.5 |

| Boro-PBPi 18, SC at 300 mg/kg | | | | | | |
|---|---|---|---|---|---|---|
| Time (h) | Calculated Concentration (ng/mL) | | | | | |
| | Mouse #13 | Mouse #14 | Mouse #15 | Mean | SD | CV (%) |
| 0.0833 | 227000 | 207000 | 212000 | 215333 | 10408 | 4.83 |
| 0.25 | 383000 | 413000 | 371000 | 389000 | 21633 | 5.56 |
| 0.5 | 328000 | 388000 | 292000 | 336000 | 48497 | 14.4 |
| 1 | 113000 | 153000 | 129000 | 131667 | 20133 | 15.3 |
| 2 | 8640 | 12400 | 11700 | 10913 | 2000 | 18.3 |
| 4 | 987 | 1130 | 976 | 1031 | 86 | 8.33 |
| 8 | 299 | 327 | 259 | 295 | 34 | 11.6 |
| 24 | 24.7 | 47.5 | 42.2 | 38.1 | 11.9 | 31.3 |

| Boro-PBPi 18, SC at 1000 mg/kg | | | | | | |
|---|---|---|---|---|---|---|
| Time (h) | Calculated Concentration (ng/mL) | | | | | |
| | Mouse #16 | Mouse #17 | Mouse #18 | Mean | SD | CV (%) |
| 0.0833 | 445000 | 465000 | 418000 | 442667 | 23587 | 5.33 |
| 0.25 | 758000 | 777000 | 886000 | 807000 | 69072 | 8.56 |
| 0.5 | 793000 | 782000 | 753000 | 776000 | 20664 | 2.66 |
| 1 | 384000 | 482000 | 468000 | 444667 | 53003 | 11.9 |
| 2 | 41400 | 86900 | 57500 | 61933 | 23072 | 37.3 |
| 4 | 2350 | 4860 | 3200 | 3470 | 1277 | 36.8 |
| 8 | 636 | 821 | 705 | 721 | 93 | 13.0 |
| 24 | 60.6 | 144 | 86.6 | 97.1 | 42.7 | 44.0 |

BLOQ = below LLOQ (lower limit of quantification), LLOQ = 2 ng/mL

**Boro-PBPi 18, pharmacokinetic parameters**

**3 mg/kg IV**

| Animal | $t_{1/2}$ (h) | $C_0$ (ng/mL) | $AUC_{Inf}$ (h*ng/mL) | $V_{ss}$ (L/kg) | CL (mL/min/kg) |
|---|---|---|---|---|---|
| Mouse #1 | 13.4 | 10726 | 3287 | 1.64 | 15.2 |
| Mouse #2 | 11.5 | 10141 | 3988 | 0.910 | 12.5 |
| Mouse #3 | 1.62 | 13010 | 3984 | 0.324 | 12.6 |
| **Mean** | **8.82** | **11292** | **3753** | **0.957** | **13.4** |
| SD | 6.31 | 1516 | 403 | 0.658 | 1.5 |
| CV (%) | 71.5 | 13.4 | 10.8 | 68.7 | 11.5 |

**10 mg/kg SC**

| Animal | $t_{1/2}$ (h) | $T_{max}$ (h) | $C_{max}$ (ng/mL) | $AUC_{inf}$ (ng×h/mL) | $F_{SC}$ (%) |
|---|---|---|---|---|---|
| Mouse #4 | 10.7 | 0.250 | 20000 | 15346 | 123 |
| Mouse #5 | 6.49 | 0.250 | 13700 | 12476 | 100 |
| Mouse #6 | 9.81 | 0.250 | 20000 | 15856 | 127 |
| **Mean** | **9.01** | **0.250** | **17900** | **14559** | **116** |
| SD | 2.24 | 0.000 | 3637 | 1822 | 15 |

**30 mg/kg SC**

| Animal | $t_{1/2}$ (h) | $T_{max}$ (h) | $C_{max}$ (ng/mL) | $AUC_{inf}$ (ng×h/mL) | $F_{SC}$ (%) |
|---|---|---|---|---|---|
| Mouse #7 | 4.26 | 0.250 | 42800 | 38664 | 103 |
| Mouse #8 | 4.19 | 0.250 | 45400 | 38540 | 103 |
| Mouse #9 | 4.65 | 0.250 | 51800 | 49647 | 132 |
| **Mean** | **4.37** | **0.250** | **46667** | **42284** | **113** |
| SD | 0.25 | 0.000 | 4632 | 6377 | 17 |

**100 mg/kg SC**

| Animal | $t_{1/2}$ (h) | $T_{max}$ (h) | $C_{max}$ (ng/mL) | $AUC_{inf}$ (ng×h/mL) | $F_{SC}$ (%) |
|---|---|---|---|---|---|
| Mouse #10 | 4.95 | 0.250 | 157000 | 131491 | 105 |
| Mouse #11 | 4.10 | 0.250 | 124000 | 115663 | 92.5 |
| Mouse #12 | 5.06 | 0.250 | 126000 | 99697 | 79.7 |
| **Mean** | **4.70** | **0.250** | **135667** | **115617** | **92.4** |
| SD | 0.522 | 0.000 | 18502 | 15897 | 12.7 |

**300 mg/kg SC**

| Animal | $t_{1/2}$ (h) | $T_{max}$ (h) | $C_{max}$ (ng/mL) | $AUC_{inf}$ (ng×h/mL) | $F_{SC}$ (%) |
|---|---|---|---|---|---|
| Mouse #13 | 3.93 | 0.250 | 383000 | 335172 | 89.3 |
| Mouse #14 | 4.69 | 0.250 | 413000 | 398135 | 106 |
| Mouse #15 | 4.79 | 0.250 | 371000 | 333745 | 88.9 |
| **Mean** | **4.47** | **0.250** | **389000** | **355684** | **94.8** |
| SD | 0.47 | 0.000 | 21633 | 36771 | 9.8 |

**1,000 mg/kg SC**

| Animal | $t_{1/2}$ (h) | $T_{max}$ (h) | $C_{max}$ (ng/mL) | $AUC_{inf}$ (ng×h/mL) | $F_{SC}$ (%) |
|---|---|---|---|---|---|
| Mouse #16 | 4.02 | 0.500 | 793000 | 875275 | 70.0 |
| Mouse #17 | 4.42 | 0.500 | 782000 | 1029974 | 82.3 |
| Mouse #18 | 4.17 | 0.250 | 886000 | 974336 | 77.9 |
| **Mean** | **4.20** | **0.417** | **820333** | **959862** | **76.7** |
| SD | 0.21 | 0.144 | 57134 | 78358 | 6.3 |

Every $t_{1/2}$ (h) was determined using data from three time points.

**Table S3.** Plasma exposure and pharmacokinetic parameters derived for boro-PBPi **21** following intravenous (IV) and subcutaneous (SC) doses in male CD-1 mice (n = 3/group).

**Boro-PBPi 21, plasma exposure (raw data)**

| Boro-PBPi 21_IV at 3 mg/kg | | | | | | |
|---|---|---|---|---|---|---|
| Time (hr) | Calculated Concentration (ng/mL) | | | | | |
| | Mouse #13 | Mouse #14 | Mouse #15 | Mean | SD | CV (%) |
| 0.0833 | 6860 | 7448 | 6765 | 7024 | 370 | 5.27 |
| 0.25 | 3052 | 3179 | 4367 | 3533 | 725 | 20.5 |
| 0.5 | 2025 | 1465 | 1810 | 1767 | 283 | 16.0 |
| 1 | 418 | 301 | 344 | 354 | 59 | 16.8 |
| 2 | 43.0 | 51.4 | 71.4 | 55.3 | 14.6 | 26.4 |
| 4 | 5.44 | BLOQ | 6.53 | 3.99 | NA | NA |
| 8 | BLOQ | BLOQ | BLOQ | NA | NA | NA |
| 24 | BLOQ | BLOQ | BLOQ | NA | NA | NA |

| Boro-PBPi 21_SC at 10 mg/kg | | | | | | |
|---|---|---|---|---|---|---|
| Time (hr) | Calculated Concentration (ng/mL) | | | | | |
| | Mouse #16 | Mouse #17 | Mouse #18 | Mean | SD | CV (%) |
| 0.25 | 7647 | 8583 | 8886 | 8372 | 646 | 7.71 |
| 0.5 | 7337 | 7917 | 8122 | 7792 | 407 | 5.23 |
| 1 | 4015 | 4607 | 4554 | 4392 | 328 | 7.46 |
| 2 | 1043 | 757 | 537 | 779 | 254 | 32.6 |
| 4 | 67.5 | 42.2 | 31.2 | 47.0 | 18.6 | 39.6 |
| 8 | 4.78 | 4.02 | 3.47 | 4.09 | 0.65 | 16.0 |
| 24 | BLOQ | BLOQ | BLOQ | NA | NA | NA |

| Boro-PBPi 21_SC at 30 mg/kg | | | | | | |
|---|---|---|---|---|---|---|
| Time (hr) | Calculated Concentration (ng/mL) | | | | | |
| | Mouse #19 | Mouse #20 | Mouse #21 | Mean | SD | CV (%) |
| 0.25 | 24680 | 30050 | 24570 | 26433 | 3133 | 11.9 |
| 0.5 | 28860 | 24580 | 22120 | 25187 | 3411 | 13.5 |
| 1 | 11740 | 15600 | 9746 | 12362 | 2976 | 24.1 |
| 2 | 1881 | 2790 | 2236 | 2302 | 458 | 19.9 |
| 4 | 100 | 214 | 130 | 148 | 59 | 39.7 |
| 8 | 20.5 | 20.0 | 13.8 | 18.1 | 3.7 | 20.7 |
| 24 | BLOQ | BLOQ | BLOQ | NA | NA | NA |

| Boro-PBPi 21_SC at 100 mg/kg | | | | | | |
|---|---|---|---|---|---|---|
| Time (hr) | Calculated Concentration (ng/mL) | | | | | |
| | Mouse #22 | Mouse #23 | Mouse #24 | Mean | SD | CV (%) |
| 0.25 | 56450 | 91700 | 49960 | 66037 | 22461 | 34.0 |
| 0.5 | 39850 | 65370 | 53940 | 53053 | 12783 | 24.1 |
| 1 | 43170 | 34310 | 40950 | 39477 | 4610 | 11.7 |
| 2 | 9898 | 9602 | 16710 | 12070 | 4021 | 33.3 |
| 4 | 1040 | 772 | 1083 | 965 | 169 | 17.5 |
| 8 | 252 | 195 | 263 | 237 | 37 | 15.6 |
| 24 | 34.5 | 31.4 | 27.9 | 31.3 | 3.3 | 10.6 |

| Boro-PBPi 21_SC at 300 mg/kg | | | | | | |
|---|---|---|---|---|---|---|
| Time (hr) | Calculated Concentration (ng/mL) | | | | | |
| | Mouse #25 | Mouse #26 | Mouse #27 | Mean | SD | CV (%) |
| 0.25 | 172500 | 199400 | 168400 | 180100 | 16840 | 9.35 |
| 0.5 | 156000 | 185700 | 163000 | 168233 | 15526 | 9.23 |
| 1 | 102000 | 156200 | 112700 | 123633 | 28707 | 23.2 |
| 2 | 43070 | 47650 | 49850 | 46857 | 3459 | 7.38 |
| 4 | 1805 | 4542 | 3557 | 3301 | 1386 | 42.0 |
| 8 | 568 | 778 | 581 | 642 | 118 | 18.3 |
| 24 | 94.9 | 121 | 86.2 | 101 | 18 | 17.9 |

BLOQ = below LLOQ (lower limit of quantification), LLOQ = 2 ng/mL

**Boro-PBPi 21, plasma pharmacokinetic parameters**

**3 mg/kg IV**

| Animal | $t_{1/2}$ (hr)* | $C_0$ (ng/mL) | $AUC_{Inf}$ (hr*ng/mL) | $V_{ss}$ (L/kg) | CL (mL/min/kg) |
|---|---|---|---|---|---|
| Mouse #1 | 0.499 | 10285 | 3069 | 0.348 | 16.3 |
| Mouse #2 | 0.320 | 11400 | 2893 | 0.306 | 17.3 |
| Mouse #3 | 0.532 | 8420 | 3161 | 0.348 | 15.8 |
| **Mean** | **0.450** | **10035** | **3041** | **0.334** | **16.5** |
| SD | 0.114 | 1506 | 136 | 0.024 | 0.7 |
| CV (%) | 25.3 | 15.0 | 4.48 | 7.28 | 4.55 |

**10 mg/kg SC**

| Animal | $t_{1/2}$ (h)* | $T_{max}$ (h) | $C_{max}$ (ng/mL) | $AUC_{inf}$ (ng×h/mL) | $F_{SC}$ (%) |
|---|---|---|---|---|---|
| Mouse #16 | 0.802 | 0.250 | 7647 | 9456 | 93.3 |
| Mouse #17 | 0.833 | 0.250 | 8583 | 9845 | 97.1 |
| Mouse #18 | 0.868 | 0.250 | 8886 | 9593 | 94.6 |
| **Mean** | **0.834** | **0.250** | **8372** | **9631** | **95.0** |
| SD | 0.033 | 0.000 | 646 | 197 | 1.9 |
| CV (%) | 3.95 | 0.00 | 7.71 | 2.05 | 2.05 |

**30 mg/kg SC**

| Animal | $t_{1/2}$ (h)* | $T_{max}$ (h) | $C_{max}$ (ng/mL) | $AUC_{inf}$ (ng×h/mL) | $F_{SC}$ (%) |
|---|---|---|---|---|---|
| Mouse #19 | 0.987 | 0.500 | 28860 | 28990 | 95.3 |
| Mouse #20 | 0.877 | 0.250 | 30050 | 33321 | 109.6 |
| Mouse #21 | 0.859 | 0.250 | 24570 | 25536 | 84.0 |
| **Mean** | **0.908** | **0.333** | **27827** | **29282** | **96.3** |
| SD | 0.069 | 0.144 | 2882 | 3901 | 12.8 |
| CV (%) | 7.65 | 43.30 | 10.36 | 13.32 | 13.32 |

**100 mg/kg SC**

| Animal | t1/2 (h) | Tmax (h) | Cmax (ng/mL) | AUCinf (ng×h/mL) | FSC (%) |
|---|---|---|---|---|---|
| Mouse #22 | 4.412 | 0.250 | 56450 | 82420 | 81.3 |
| Mouse #23 | 4.718 | 0.250 | 91700 | 92300 | 91.1 |
| Mouse #24 | 4.060 | 0.500 | 53940 | 94760 | 93.5 |
| **Mean** | **4.397** | **0.333** | **67363** | **89826** | **88.6** |
| SD | 0.329 | 0.144 | 21114 | 6531 | 6.4 |
| CV (%) | 7.49 | 43.30 | 31.34 | 7.27 | 7.27 |

**300 mg/kg SC**

| Animal | $t_{1/2}$ (h) | $T_{max}$ (h) | $C_{max}$ (ng/mL) | $AUC_{inf}$ (ng×h/mL) | $F_{SC}$ (%) |
|---|---|---|---|---|---|
| Mouse #25 | 5.053 | 0.250 | 172500 | 255277 | 83.9 |
| Mouse #26 | 4.257 | 0.250 | 199400 | 331224 | 108.9 |
| Mouse #27 | 4.152 | 0.250 | 168400 | 280207 | 92.1 |
| **Mean** | **4.487** | **0.250** | **180100** | **288903** | **95.0** |
| SD | 0.493 | 0.000 | 16840 | 38713 | 12.7 |
| CV (%) | 10.98 | 0.00 | 9.35 | 13.40 | 13.40 |

*Calculated half-life. Terminal half-life was not determined as concentrations at later timepoints were below the limit of detection.

**Table S4.** In vivo efficacy data for boro-PBPi **18** in the murine vaginal infection model with WT *N. gonorrhoeae* strain FA1090 (ATCC 700825).

| **Group** | **Treatment** | **Dose (mg/kg) (Route)** | **Animal No.** | **CFU/mL Time post inoculation 2, 26 h** | **Log (CFU/mL) Time post inoculation 2, 26 h** |
|---|---|---|---|---|---|
| 1 | Initial counts, 2 h PI | NA | 1 | 77000 | 4.89 |
| | | | 2 | 34000 | 4.53 |
| | | | 3 | 56000 | 4.75 |
| | | | 4 | 38000 | 4.58 |
| | | | 5 | 37000 | 4.57 |
| | | | Mean | 48400 | 4.66 |
| | | | SEM | 8130 | 0.07 |
| 2 | Vehicle (50 mM PBS pH 6-7) | 10 mL/kg, SC, BID | 1 | 37000 | 4.57 |
| | | | 2 | 171000 | 5.23 |
| | | | 3 | 570000 | 5.76 |
| | | | 4 | 276000 | 5.44 |
| | | | 5 | 390000 | 5.59 |
| | | | Mean | 289000 | 5.32 |
| | | | SEM | 91300 | 0.21 |
| 3 | Ceftriaxone | 1 mg/kg IV, QD | 1 | 10 | 1.00 |
| | | | 2 | <10[a] | <1.00 |
| | | | 3 | <10[a] | <1.00 |
| | | | 4 | <10[a] | <1.00 |
| | | | 5 | 70 | 1.85 |
| | | | Mean | 22 | 1.17* |
| | | | SEM | 12 | 0.17 |
| 4 | Boro-PBPi **18** | 200 mg/kg, SC, BID | 1 | <10[a] | <1.00 |
| | | | 2 | <10[a] | <1.00 |
| | | | 3 | <10[a] | <1.00 |
| | | | 4 | 30 | 1.48 |
| | | | 5 | <10[a] | <1.00 |
| | | | Mean | 14 | 1.10* |
| | | | SEM | 4 | 0.10 |
| 5 | Boro-PBPi **18** | 50 mg/kg, SC, BID | 1 | <10[a] | <1.00 |
| | | | 2 | 120 | 2.08 |
| | | | 3 | <10[a] | <1.00 |
| | | | 4 | 20 | 1.30 |
| | | | 5 | 70 | 1.85 |
| | | | Mean | 46 | 1.45* |
| | | | SEM | 21.6 | 0.22 |
| 6 | Boro-PBPi **18** | 7.5 mg/kg, SC, BID | 1 | 30 | 1.48 |
| | | | 2 | <10[a] | <1.00 |

| Group | Treatment | Dose (mg/kg) (Route) | Animal No. | CFU/mL Time post inoculation 2, 26 h | Log (CFU/mL) Time post inoculation 2, 26 h |
|---|---|---|---|---|---|
| | | | 3 | 20 | 1.30 |
| | | | 4 | 20 | 1.30 |
| | | | 5 | <10[a] | <1.00 |
| | | | Mean | 18 | 1.22* |
| | | | SEM | 3.74 | 0.09 |
| 7 | Boro-PBPi **18** | 0.25 mg/kg, SC, BID | 1 | 33000 | 4.52 |
| | | | 2 | 31000 | 4.49 |
| | | | 3 | 44000 | 4.64 |
| | | | 4 | 82000 | 4.91 |
| | | | 5 | 181000 | 5.26 |
| | | | Mean | 74200 | 4.76* |
| | | | SEM | 28200 | 0.14 |

[a]The bacterial count was below the limit of detection (CFU/mL = 10). A significant difference ($p < 0.05$) compared to the vehicle group as determined by one-way ANOVA is indicated as asterisk (*). The data are plotted in **Fig. S8**.

**Table S5.** In vivo efficacy data and comparison of groups for boro-PBPi **21** in the murine vaginal infection model with ceftriaxone-resistant *N. gonorrhoeae* strain H041.

**Comparison of groups (p-values)**

| Test article | PBS | Ceftriaxone | Boro-PBPi 21 | | |
|---|---|---|---|---|---|
| | **5 mL/kg SC, q8h** | **120 mg/kg SC, q8h** | **10 mg/kg SC, q12h** | **200 mg/kg SC, q24h** | **150 mg/kg SC, q12h** |
| **Ceftriaxone 120 mg/kg SC, q8h** | 0.6324 | - | | | |
| **Boro-PBPi 21 10 mg/kg SC, q12h** | **0.0033** | 0.9889 | - | | |
| **Boro-PBPi 21 200 mg/kg SC, q24h** | **<0.0001** | **0.0059** | 0.9376 | - | |
| **Boro-PBPi 21 150 mg/kg SC, q12h** | **<0.0001** | **0.0033** | 0.6322 | >0.9999 | - |
| **Boro-PBPi 21 150 mg/kg SC, q8h** | **<0.0001** | **0.0008** | 0.2277 | >0.9999 | >0.9999 |

p-values were determined by two-way ANOVA analysis using $\log_{10}$(CFU/mL) values (significant differences [$p < 0.05$] shown in red). The degrees of freedom and the F values calculated from the comparison of groups were $F (5, 54) = 13.34$.

**Raw CFU/mL data**

| **PBS, 5 mL/kg SC, q8h (CFU/mL)** | | | | | | | | | | |
|---|---|---|---|---|---|---|---|---|---|---|
| **Days Post-Treatment** | **Mouse #1** | **Mouse #2** | **Mouse #3** | **Mouse #4** | **Mouse #5** | **Mouse #6** | **Mouse #7** | **Mouse #8** | **Mouse #9** | **Mouse #10** |
| 0 | 5445 | 15100 | 10400 | 34700 | 176000 | 226000 | 100 | 2360000 | 322000 | 5200000 |
| 1 | 177000 | 42000 | 11900 | 243000 | 30950 | 20900 | 444000 | 1280000 | 96000 | 710000 |
| 2 | 420 | 60 | 80 | 20 | 164000 | 3610 | 778000 | 3600000 | 120000 | 456000 |
| 3 | 79500 | 280 | 7500 | 20 | 10900 | 22850 | 1860000 | 134000 | 116000 | 84000 |
| 4 | 50500 | 5890 | 2610 | 20 | 8045 | 20550 | 3500000 | 3350000 | 444000 | 140 |
| 5 | 76500 | 60 | 100 | 20 | 20 | 41500 | 9700000 | 1070000 | 14700 | 100 |
| 6 | 5555 | 20 | 20 | 20 | 20 | 120500 | 7900000 | 163000 | 45000 | 20 |
| 7 | 17500 | 20 | 540 | 20 | 20 | 64500 | 3300000 | 54000 | 980 | 40 |
| 8 | 20 | 20 | 4600 | 20 | 20 | 223000 | 5900000 | 9600 | 2545 | 2630 |

| Ceftriaxone, 120 mg/kg SC, q8h (CFU/mL) | | | | | | | | | | |
|---|---|---|---|---|---|---|---|---|---|---|
| Days Post-Treatment | Mouse #11 | Mouse #12 | Mouse #13 | Mouse #14 | Mouse #15 | Mouse #16 | Mouse #17 | Mouse #18 | Mouse #19 | Mouse #20 |
| 0 | 1200 | 533000 | 3140000 | 200 | 19800 | 1760 | 1220000 | 1240000 | 1430000 | 493000 |
| 1 | 20 | 20 | 40 | 20 | 98500 | 11000 | 2555 | 88000 | 420 | 329000 |
| 2 | 20 | 20 | 3680 | 20 | 20 | 20 | 12550 | 520 | 48300 | 4780 |
| 3 | 20 | 20 | 3140 | 20 | 580 | 20 | 360 | 20 | 2600000 | 120 |
| 4 | 20 | 20 | 160 | 20 | 28100 | 108000 | 5890 | 20 | 2250000 | 1980 |
| 5 | 8090 | 20 | 20 | 20 | 148000 | 164000 | 22500 | 20 | 1340000 | 56000 |
| 6 | 20 | 20 | 20 | 20 | 280 | 986000 | 14100 | 20 | 2250000 | 5780 |
| 7 | 2280 | 20 | 20 | 20 | 72000 | 296000 | 2060 | 20 | 828000 | 80 |
| 8 | 40 | 20 | 20 | 20 | 1480 | 444000 | 8680 | 20 | 1120000 | 7280 |

585

| Boro-PBPi 21, 10 mg/kg SC, q12h (CFU/mL) | | | | | | | | | | |
|---|---|---|---|---|---|---|---|---|---|---|
| Days Post-Treatment | Mouse #21 | Mouse #22 | Mouse #23 | Mouse #24 | Mouse #25 | Mouse #26 | Mouse #27 | Mouse #28 | Mouse #29 | Mouse #30 |
| 0 | 60 | 4650 | 100 | 7230 | 52450 | 1260 | 3720 | 115500 | 70000 | 1400000 |
| 1 | 97000 | 26000 | 18450 | 2630 | 267000 | 720 | 176000 | 80 | 120 | 82000 |
| 2 | 520 | 300 | 560 | 27850 | 1300 | 20 | 1800 | 20 | 20 | 20 |
| 3 | 20 | 100 | 80 | 20 | 640 | 20 | 800 | 20 | 20 | 20 |
| 4 | 20 | 20 | 20 | 20 | 20 | 1280 | 17550 | 20 | 20 | 20 |
| 5 | 20 | 20 | 20 | 20 | 20 | 226000 | 103500 | 20 | 20 | 20 |
| 6 | 20 | 20 | 20 | 20 | 20 | 1400000 | 224000 | 20 | 20 | 20 |
| 7 | 20 | 20 | 20 | 20 | 20 | 220000 | 31000 | 20 | 20 | 20 |
| 8 | 20 | 20 | 20 | 20 | 20 | 118500 | 11600 | 20 | 20 | 20 |

586

| Boro-PBPi 21, 200 mg/kg SC, q24h (CFU/mL) | | | | | | | | | | |
|---|---|---|---|---|---|---|---|---|---|---|
| Days Post-Treatment | Mouse #31 | Mouse #32 | Mouse #33 | Mouse #34 | Mouse #35 | Mouse #36 | Mouse #37 | Mouse #38 | Mouse #39 | Mouse #40 |
| 0 | 40 | 2000 | 3640 | 789000 | 19100 | 651000 | 193000 | 232000 | 184500 | 3900000 |
| 1 | 20 | 20 | 20 | 20 | 1120 | 20 | 1760 | 20 | 140 | 20 |
| 2 | 20 | 20 | 20 | 20 | 20 | 20 | 40 | 20 | 20 | 20 |
| 3 | 20 | 20 | 20 | 20 | 20 | 20 | 20 | 20 | 20 | 20 |
| 4 | 20 | 20 | 20 | 20 | 20 | 20 | 20 | 20 | 20 | 20 |
| 5 | 20 | 20 | 20 | 20 | 20 | 20 | 20 | 20 | 20 | 20 |
| 6 | 20 | 20 | 20 | 20 | 20 | 20 | 20 | 20 | 20 | 20 |
| 7 | 20 | 20 | 20 | 20 | 20 | 20 | 20 | 20 | 20 | 20 |
| 8 | 20 | 20 | 20 | 20 | 20 | 20 | 20 | 20 | 20 | 20 |

587

| Boro-PBPi 21, 150 mg/kg SC, q12h (CFU/mL) | | | | | | | | | | |
|---|---|---|---|---|---|---|---|---|---|---|
| Days Post-Treatment | Mouse #41 | Mouse #42 | Mouse #43 | Mouse #44 | Mouse #45 | Mouse #46 | Mouse #47 | Mouse #48 | Mouse #49 | Mouse #50 |
| 0 | 1320 | 142445 | 1300 | 8330 | 8480 | 2760 | 35700 | 22450 | 8160 | 355000 |
| 1 | 20 | 20 | 20 | 20 | 20 | 20 | 20 | 3840 | 20 | 5500 |
| 2 | 20 | 20 | 20 | 20 | 20 | 20 | 20 | 20 | 20 | 20 |
| 3 | 20 | 20 | 20 | 20 | 20 | 20 | 20 | 20 | 20 | 20 |
| 4 | 20 | 20 | 20 | 20 | 20 | 20 | 20 | 20 | 20 | 20 |
| 5 | 20 | 20 | 20 | 20 | 20 | 20 | 20 | 20 | 20 | 20 |
| 6 | 20 | 20 | 20 | 20 | 20 | 20 | 20 | 20 | 20 | 20 |

| Boro-PBPi 21, 150 mg/kg SC, q12h (CFU/mL) | | | | | | | | | | |
|---|---|---|---|---|---|---|---|---|---|---|
| 7 | 20 | 20 | 20 | 20 | 20 | 20 | 20 | 20 | 20 | 20 |
| 8 | 20 | 20 | 20 | 20 | 20 | 20 | 20 | 20 | 20 | 20 |

| Boro-PBPi 21, 150 mg/kg SC, q8h (CFU/mL) | | | | | | | | | | |
|---|---|---|---|---|---|---|---|---|---|---|
| **Days Post-Treatment** | **Mouse #51** | **Mouse #52** | **Mouse #53** | **Mouse #54** | **Mouse #55** | **Mouse #56** | **Mouse #57** | **Mouse #58** | **Mouse #59** | **Mouse #60** |
| 0 | 540 | 20 | 3210 | 40 | 1322 | 11700 | 4470 | 5000 | 2360 | 2890 |
| 1 | 40 | 20 | 20 | 20 | 20 | 80 | 20 | 20 | 20 | 20 |
| 2 | 20 | 20 | 20 | 20 | 20 | 20 | 20 | 20 | 20 | 20 |
| 3 | 20 | 20 | 20 | 20 | 20 | 20 | 20 | 20 | 20 | 20 |
| 4 | 20 | 20 | 20 | 20 | 20 | 20 | 20 | 20 | 20 | 20 |
| 5 | 20 | 20 | 20 | 20 | 20 | 20 | 20 | 20 | 20 | 20 |
| 6 | 20 | 20 | 20 | 20 | 20 | 20 | 20 | 20 | 20 | 20 |
| 7 | 20 | 20 | 20 | 20 | 20 | 20 | 20 | 20 | 20 | 20 |
| 8 | 20 | 20 | 20 | 20 | 20 | 20 | 20 | 20 | 20 | 20 |

The lower limit of detection was 20 CFU/mL. The values of 20 CFU/mL indicate less than or equal to 20 CFU/mL.

**Table S6.** Percents of polymorphonuclear leukocytes (PMN) in samples from the murine vaginal infection model with ceftriaxone-resistant *N. gonorrhoeae* strain H041.

| PBS (% PMN out of 100 cells) | | | | | | | | | | |
|---|---|---|---|---|---|---|---|---|---|---|
| Days Post-Treatment | Mouse #1 | Mouse #2 | Mouse #3 | Mouse #4 | Mouse #5 | Mouse #6 | Mouse #7 | Mouse #8 | Mouse #9 | Mouse #10 |
| 0 | 6 | 0 | 13 | 3 | 0 | 0 | 0 | 0 | 0 | 4 |
| 1 | 10 | 0 | 41 | 0 | 0 | 0 | 0 | 0 | 0 | 10 |
| 2 | 16 | 0 | 55 | 0 | 0 | 0 | 6 | 0 | 0 | 11 |
| 3 | 21 | 0 | 80 | 0 | 4 | 0 | 4 | 0 | 5 | 21 |
| 4 | 28 | 29 | 46 | 0 | 5 | 0 | 8 | 61 | 21 | 12 |
| 5 | 5 | 12 | 48 | 0 | 0 | 0 | 0 | 58 | 48 | 5 |
| 6 | 3 | 49 | 41 | 0 | 7 | 0 | 0 | 49 | 17 | 30 |
| 7 | 20 | 11 | 28 | 0 | 17 | 4 | 15 | 50 | 26 | 7 |
| 8 | 32 | 9 | 50 | 0 | 1 | 0 | 2 | 8 | 18 | 0 |

| Ceftriaxone, 120 mg/kg SC, q8h (% PMN out of 100 cells) | | | | | | | | | | |
|---|---|---|---|---|---|---|---|---|---|---|
| Days Post-Treatment | Mouse #11 | Mouse #12 | Mouse #13 | Mouse #14 | Mouse #15 | Mouse #16 | Mouse #17 | Mouse #18 | Mouse #19 | Mouse #20 |
| 0 | 0 | 0 | 0 | 0 | 0 | 0 | 0 | 0 | 0 | 0 |
| 1 | 0 | 16 | 4 | 0 | 0 | 0 | 34 | 7 | 0 | 0 |
| 2 | 0 | 21 | 40 | 0 | 0 | 0 | 39 | 57 | 0 | 25 |
| 3 | 19 | 58 | 51 | 0 | 7 | 0 | 48 | 68 | 9 | 31 |
| 4 | 41 | 53 | 45 | 0 | 46 | 3 | 45 | 24 | 14 | 52 |
| 5 | 3 | 65 | 37 | 0 | 14 | 5 | 29 | 20 | 17 | 54 |
| 6 | 36 | 52 | 46 | 0 | 34 | 60 | 36 | 23 | 24 | 39 |
| 7 | 46 | 61 | 67 | 0 | 45 | 32 | 20 | 10 | 26 | 26 |
| 8 | 53 | 55 | 63 | 7 | 46 | 58 | 21 | 13 | 32 | 40 |

| Boro-PBPi 21, 10 mg/kg SC, q12h (% PMN out of 100 cells) | | | | | | | | | | |
|---|---|---|---|---|---|---|---|---|---|---|
| Days Post-Treatment | Mouse #21 | Mouse #22 | Mouse #23 | Mouse #24 | Mouse #25 | Mouse #26 | Mouse #27 | Mouse #28 | Mouse #29 | Mouse #30 |
| 0 | 0 | 3 | 0 | 0 | 0 | 0 | 0 | 0 | 0 | 0 |
| 1 | 0 | 41 | 0 | 0 | 0 | 13 | 0 | 0 | 0 | 19 |
| 2 | 0 | 56 | 0 | 33 | 24 | 4 | 0 | 0 | 0 | 44 |
| 3 | 10 | 76 | 34 | 42 | 17 | 0 | 13 | 0 | 8 | 90 |
| 4 | 20 | 21 | 15 | 45 | 12 | 0 | 26 | 0 | 8 | 71 |
| 5 | 24 | 12 | 20 | 34 | 4 | 0 | 34 | 0 | 13 | 56 |
| 6 | 10 | 70 | 4 | 78 | 13 | 3 | 53 | 0 | 51 | 75 |
| 7 | 21 | 48 | 38 | 37 | 23 | 5 | 43 | 0 | 6 | 80 |
| 8 | 49 | 49 | 33 | 32 | 4 | 4 | 45 | 0 | 8 | 81 |

| Boro-PBPi 21, 200 mg/kg SC, q24h (% PMN out of 100 cells) | | | | | | | | | | |
|---|---|---|---|---|---|---|---|---|---|---|
| Days Post-Treatment | Mouse #31 | Mouse #32 | Mouse #33 | Mouse #34 | Mouse #35 | Mouse #36 | Mouse #37 | Mouse #38 | Mouse #39 | Mouse #40 |
| 0 | 5 | 0 | 0 | 0 | 0 | 0 | 0 | 0 | 0 | 0 |
| 1 | 0 | 0 | 0 | 12 | 0 | 26 | 0 | 12 | 3 | 0 |
| 2 | 0 | 2 | 0 | 33 | 19 | 53 | 6 | 20 | 77 | 1 |
| 3 | 0 | 3 | 0 | 56 | 50 | 57 | 21 | 41 | 78 | 13 |
| 4 | 0 | 7 | 0 | 64 | 59 | 37 | 37 | 10 | 57 | 4 |
| 5 | 0 | 9 | 0 | 30 | 16 | 31 | 43 | 36 | 29 | 4 |
| 6 | 0 | 0 | 0 | 45 | 18 | 20 | 33 | 22 | 56 | 0 |
| 7 | 0 | 0 | 0 | 22 | 0 | 34 | 27 | 39 | 65 | 0 |
| 8 | 0 | 3 | 0 | 44 | 30 | 30 | 0 | 48 | 84 | 0 |

| Boro-PBPi 21, 150 mg/kg SC, q12h (% PMN out of 100 cells) | | | | | | | | | | |
|---|---|---|---|---|---|---|---|---|---|---|
| Days Post-Treatment | Mouse #41 | Mouse #42 | Mouse #43 | Mouse #44 | Mouse #45 | Mouse #46 | Mouse #47 | Mouse #48 | Mouse #49 | Mouse #50 |
| 0 | 0 | 0 | 0 | 0 | 4 | 0 | 0 | 0 | 0 | 0 |
| 1 | 0 | 0 | 0 | 0 | 4 | 25 | 0 | 0 | 0 | 0 |
| 2 | 0 | 0 | 0 | 0 | 5 | 32 | 4 | 1 | 5 | 10 |
| 3 | 0 | 0 | 0 | 0 | 0 | 53 | 27 | 67 | 44 | 29 |
| 4 | 12 | 0 | 0 | 0 | 0 | 57 | 57 | 42 | 31 | 30 |
| 5 | 0 | 0 | 0 | 0 | 0 | 46 | 18 | 40 | 40 | 30 |
| 6 | 0 | 0 | 0 | 13 | 0 | 35 | 20 | 51 | 11 | 20 |
| 7 | 0 | 2 | 0 | 64 | 10 | 32 | 7 | 47 | 34 | 30 |
| 8 | 0 | 23 | 25 | 65 | 13 | 55 | 3 | 43 | 19 | 33 |

| Boro-PBPi 21, 150 mg/kg SC, q8h (% PMN out of 100 cells) | | | | | | | | | | |
|---|---|---|---|---|---|---|---|---|---|---|
| Days Post-Treatment | Mouse #51 | Mouse #52 | Mouse #53 | Mouse #54 | Mouse #55 | Mouse #56 | Mouse #57 | Mouse #58 | Mouse #59 | Mouse #60 |
| 0 | 0 | 0 | 0 | 3 | 0 | 0 | 0 | 2 | 0 | 1 |
| 1 | 0 | 0 | 0 | 0 | 0 | 5 | 6 | 15 | 0 | 0 |
| 2 | 53 | 0 | 0 | 0 | 0 | 71 | 21 | 46 | 2 | 13 |
| 3 | 81 | 0 | 3 | 0 | 0 | 33 | 35 | 52 | 63 | 5 |
| 4 | 61 | 0 | 0 | 0 | 0 | 46 | 35 | 0 | 29 | 23 |
| 5 | 59 | 0 | 0 | 0 | 0 | 17 | 16 | 15 | 50 | 0 |
| 6 | 50 | 3 | 0 | 0 | 0 | 18 | 48 | 50 | 58 | 21 |
| 7 | 50 | 0 | 0 | 0 | 0 | 38 | 50 | 44 | 23 | 0 |
| 8 | 19 | 0 | 0 | 3 | 0 | 0 | 50 | 35 | 30 | 0 |

**Table S7.** *N. gonorrhoeae* strain description and variants in genes associated with β-lactam resistance.

| Strain | Strain information | Source | ***penA*** **allele type (NG-STAR, PBP2)** | PBP1 | MtrR efflux regulator | mtCDE-mtrR intergenic* | PorB1b porin |
|---|---|---|---|---|---|---|---|
| **ATCC 49226** | CLSI QC strain | ATCC | *penA-22* | T375A, S666F | A39T | G120bp→A | Not present, PorB1a type porin |
| **FA19** | Wild type | Robert Nicholas | *penA-15*, reference (WT) | WT | WT | WT | Not present, PorB1a type porin |
| **FA1090** | ATCC 700825 | Ann Jerse | *penA-1* | WT | WT | WT | WT PorB1b |
| **WHO F** | CDC-0901 | CDC | *penA-15* | WT | WT | WT | Not present, PorB1a type porin |
| **WHO G** | CDC-0902 | CDC | *penA-2* | L421P | H105Y | Del_A193bp | Not present, PorB1a type porin |
| **WHO M** | CDC-0905 | CDC | *penA-2* | L421P | G45D | Del_A193bp | 23aa variants (key: G120K, A121D) |
| **MS11** | | Ann Jerse | *penA-22* | L421P | A39T | WT | 16 aa variants (key: G120D, A121D) |
| **WHO L** | CDC-0904 | CDC | *penA-7* | L421P | G45D | G120bp→A | 19 aa variants (key: G120K, A121D) |
| **WHO K** | CDC-0903 | CDC | Mosaic *penA-10* | L421P | G45D | Del_A193bp | 22 aa variants (key: G120K, A121D) |
| **H041** | WHO X, CDC-0912 | Ann Jerse | Mosaic *penA-37* | L421P | H105Y | Del_A193bp | 28 aa variants (key: G120K, A121D) |
| **CDC-0914** | WHO Z, A8806 | CDC | Mosaic *penA-64* | L421P | A40D, T86A | A193bp→C | 21 aa variants (key: G120K, A121D) |
| **WHO Q** | G7944, NCTC 14208 | NCTC | Mosaic *penA-60* | L421P | G45D | Del_A193bp | 13 aa variants (key: G120K, A121D) |
| **CDC-0197** | | CDC | Mosaic *penA-34* | L421P | H105Y | Del_A193bp | 26 aa variants (key: G120K A121N) |
| **F89** | WHO Y, CDC-0913 | Ann Jerse | Mosaic *penA-42* | L421P | H105Y | Del_A193bp | 25 aa variants (key: G120K A121N) |

**Table S8.** PBP2 variants in *N. gonorrhoeae* strains.

| Strain | ***penA*** **allele type (NG-STAR, PBP2)** | Variant No. | A311 | I312 | V316 | D345 ins | T483 | A501 | G542 | G545 | P551 |
|---|---|---|---|---|---|---|---|---|---|---|---|
| **ATCC 49226** | *penA-22* | 11 | - | - | - | Y | - | - | - | - | - |
| **FA19** | *penA-15*, reference (WT) | - | - | - | - | - | - | - | - | - | - |
| **FA1090** | *penA-1* | 2 | - | - | - | Y | - | - | - | - | - |
| **WHO F** | *penA-15* | - | - | - | - | - | - | - | - | - | - |
| **WHO G** | *penA-2* | 5 | - | - | - | Y | - | - | - | - | - |
| **WHO M** | *penA-2* | 5 | - | - | - | Y | - | - | - | - | - |
| **MS11** | *penA-22* | 10 | - | - | - | Y | - | - | - | - | - |
| **WHO L** | *penA-7* | 7 | - | - | - | Y | - | V | S | - | - |
| **WHO K** | Mosaic *penA-10* | 59 | - | M | T | - | - | - | - | S | - |
| **H041** | Mosaic *penA-37* | 61 | V | M | P | - | S | - | - | S | V |
| **CDC-0914** | Mosaic *penA-64* | 57 | V | M | T | - | S | - | - | S | V |
| **WHO Q** | Mosaic *penA-60* | 49 | V | M | T | - | S | - | - | S | V |
| **CDC-0197** | Mosaic *penA-34* | 51 | - | M | T | - | - | - | - | S | - |
| **F89** | Mosaic *penA-42* | 52 | - | M | T | - | - | P | - | S | - |

**Table S9.** Boro-PBPi activity against zoliflodacin-resistant mutants generated in strain WHO K.

| Strain | 15 | 18 | 21 | CRO | ZOLI | GEPO | CIP | AZM | WGS result |
|---|---|---|---|---|---|---|---|---|---|
| **WHO K Parent** | 0.5 | 0.5 | 0.25 | 0.03 | 0.12 | 0.5 | 64 | 0.25 | - |
| **WHO K Colony 3** | 0.5 | 0.25 | 0.12 | 0.03 | 4 | 0.25 | 32 | 0.06 | GyrB D429N |
| **WHO K Colony 7** | 0.25 | 0.25 | 0.12 | 0.03 | 4 | 0.25 | 32 | 0.06 | GyrB D429N |
| **WHO K Colony 10** | 0.5 | 0.25 | 0.12 | 0.06 | 4 | 0.25 | 64 | 0.06 | GyrB D429N |
| **WHO K Colony 13** | 0.5 | 0.25 | 0.12 | 0.03 | 4 | 0.25 | 32 | 0.12 | GyrB D429N |

Broth MIC (µg/mL)

Abbreviations: CRO, ceftriaxone; ZOLI, zoliflodacin; GEPO, gepotidacin; AZM, azithromycin; CIP, ciprofloxacin; PEN G, penicillin G; WGS, whole genome sequence.

**Table S10.** Agar MIC (µg/mL) against each of 44 CDC *N. gonorrhoeae* strains.

| Strain ID | *penA* allele | 12 | 15 | 18 | 21 | CRO | ZOLI | GEPO | AZM | CIP | PEN G |
|---|---|---|---|---|---|---|---|---|---|---|---|
| **CDC-0166** | *penA*-34 mosaic | 2 | 0.5 | 0.5 | 0.12 | 0.06 | 0.12 | 0.5 | 0.5 | 16 | 2 |
| **CDC-0167** | *penA*-2 | 0.06 | 0.016 | 0.016 | 0.016 | 0.004 | 0.06 | 0.25 | 8 | ≤0.004 | 0.12 |
| **CDC-0168** | *penA*-34 mosaic | 2 | 0.12 | 0.12 | 0.06 | 0.06 | 0.06 | 0.25 | 0.25 | 16 | 1 |
| **CDC-0169** | *penA*-34 mosaic | 4 | 0.25 | 0.25 | 0.12 | 0.06 | 0.12 | 0.5 | 0.5 | 16 | 1 |
| **CDC-0170** | *penA*-34 mosaic | 2 | 0.25 | 0.25 | 0.06 | 0.06 | 0.12 | 0.5 | 0.25 | 16 | 1 |
| **CDC-0171** | *penA*-34 mosaic | 2 | 0.12 | 0.12 | 0.06 | 0.06 | 0.06 | 0.5 | 0.25 | 16 | 2 |
| **CDC-0172** | *penA*-34 mosaic | 2 | 0.25 | 0.5 | 0.12 | 0.06 | 0.06 | 0.5 | 0.25 | 16 | 1 |
| **CDC-0173** | *penA*-34 mosaic | 4 | 0.25 | 0.5 | 0.12 | 0.06 | 0.12 | 0.5 | 0.5 | 16 | 1 |
| **CDC-0174** | *penA*-34 mosaic | 4 | 0.5 | 0.5 | 0.12 | 0.12 | 0.12 | 0.5 | 0.5 | 16 | 2 |
| **CDC-0175** | *penA*-2 | 0.06 | 0.016 | 0.016 | 0.016 | 0.004 | 0.12 | 0.5 | 4 | ≤0.004 | 0.12 |
| **CDC-0176** | *penA*-34 mosaic | 2 | 0.25 | 0.25 | 0.06 | 0.06 | 0.12 | 1 | 0.25 | 16 | 2 |
| **CDC-0177** | *penA*-2 | 0.25 | 0.03 | 0.03 | 0.03 | 0.008 | 0.25 | 1 | 1 | 0.016 | 0.5 |
| **CDC-0178** | *penA*-34 mosaic | 4 | 0.5 | 0.5 | 0.06 | 0.06 | 0.12 | 0.25 | 0.5 | 16 | 2 |
| **CDC-0179** | *penA*-2 | 0.12 | 0.016 | 0.016 | 0.016 | 0.004 | 0.06 | 0.25 | 8 | ≤0.004 | 0.12 |
| **CDC-0180** | *penA*-34 mosaic | 4 | 0.5 | 0.5 | 0.12 | 0.12 | 0.12 | 0.25 | 0.25 | 16 | 2 |
| **CDC-0182** | *penA*-34 mosaic | 4 | 0.5 | 0.5 | 0.12 | 0.12 | 0.12 | 1 | 0.5 | 16 | 2 |
| **CDC-0183** | *penA*-34 mosaic | 4 | 0.25 | 0.25 | 0.06 | 0.06 | 0.12 | 0.5 | 1 | 32 | 4 |
| **CDC-0184** | *penA*-34 mosaic | 2 | 0.25 | 0.25 | 0.06 | 0.06 | 0.12 | 1 | 0.25 | 16 | 2 |
| **CDC-0185** | *penA*-34 mosaic | 2 | 0.25 | 0.5 | 0.06 | 0.12 | 0.12 | 1 | 0.25 | 16 | 2 |
| **CDC-0186** | *penA*-34 mosaic | 4 | 0.5 | 0.5 | 0.12 | 0.12 | 0.06 | 0.25 | 0.25 | 16 | 2 |
| **CDC-0187** | *penA*-34 mosaic | 1 | 0.12 | 0.12 | 0.06 | 0.03 | 0.25 | 1 | 1 | 8 | 0.5 |
| **CDC-0188** | *penA*-34 mosaic | 2 | 0.25 | 0.25 | 0.06 | 0.06 | 0.12 | 0.5 | 0.25 | 16 | 2 |
| **CDC-0189** | *penA*-34 mosaic | 2 | 0.25 | 0.25 | 0.06 | 0.06 | 0.12 | 1 | 0.25 | 16 | 2 |
| **CDC-0190** | *penA*-34 mosaic | 4 | 0.5 | 0.5 | 0.12 | 0.12 | 0.06 | 1 | 0.25 | 16 | 1 |
| **CDC-0191** | *penA*-34 mosaic | 2 | 0.5 | 0.5 | 0.06 | 0.12 | 0.12 | 1 | 0.5 | 16 | 4 |
| **CDC-0192** | *penA*-34 mosaic | 2 | 0.25 | 0.5 | 0.06 | 0.06 | 0.12 | 0.5 | 0.5 | 16 | 2 |

| Strain ID | *penA* allele | 12 | 15 | 18 | 21 | CRO | ZOLI | GEPO | AZM | CIP | PEN G |
|---|---|---|---|---|---|---|---|---|---|---|---|
| **CDC-0193** | *penA*-9 (*penA*-2 + P551L) | 2 | 0.5 | 0.5 | 0.12 | 0.12 | 0.25 | 1 | 1 | 0.016 | 2 |
| **CDC-0195** | *penA*-34 mosaic | 4 | 0.25 | 0.25 | 0.12 | 0.06 | 0.12 | 1 | 0.5 | 16 | 2 |
| **CDC-0196** | *penA*-34 mosaic | 2 | 0.5 | 0.5 | 0.12 | 0.12 | 0.12 | 0.5 | 0.5 | 16 | 2 |
| **CDC-0197** | *penA*-34 mosaic | 2 | 0.25 | 0.25 | 0.06 | 0.03 | 0.06 | 0.25 | 4.1 | 16 | 1 |
| **CDC-0198** | *penA*-34 mosaic | 2 | 0.25 | 0.25 | 0.06 | 0.06 | 0.12 | 0.5 | 0.5 | 2 | 2 |
| **CDC-0199** | *penA*-9 (*penA*-2 + P551L) | 1 | 0.25 | 0.25 | 0.12 | 0.06 | 0.25 | 0.5 | 0.5 | 1 | 1 |
| **CDC-0200** | *penA*-34 mosaic | 8 | 0.5 | 0.5 | 0.25 | 0.12 | 0.12 | 0.5 | 0.5 | 8 | 2 |
| **CDC-0201** | *penA*-34 mosaic | 2 | 0.25 | 0.25 | 0.06 | 0.06 | 0.12 | 0.5 | 0.25 | 2 | 2 |
| **CDC-0203** | *penA*-34 mosaic | 4 | 0.5 | 0.5 | 0.12 | 0.06 | 0.12 | 0.5 | 0.25 | 4 | 2 |
| **CDC-0204** | *penA*-34 mosaic | 2 | 0.5 | 0.25 | 0.06 | 0.25 | 0.12 | 1 | 0.25 | 2 | 2 |
| **CDC-0206** | *penA*-34 mosaic | 2 | 0.5 | 0.5 | 0.06 | 0.06 | 0.12 | 1 | 0.25 | 2 | 1 |
| **CDC-0207** | *penA*-34 mosaic | 2 | 0.5 | 0.5 | 0.12 | 0.12 | 0.12 | 0.5 | 0.5 | 2 | 1 |
| **CDC-0208** | *penA*-34 mosaic | 2 | 0.25 | 0.25 | 0.06 | 0.06 | 0.12 | 0.5 | 0.5 | 2 | 2 |
| **CDC-0209** | *penA*-34 mosaic | 2 | 0.25 | 0.25 | 0.06 | 0.06 | 0.12 | 1 | 0.5 | 2 | 2 |
| **CDC-0210** | *penA*-34 mosaic | 4 | 0.25 | 0.5 | 0.12 | 0.06 | 0.12 | 1 | 0.25 | 4 | 1 |
| **CDC-0212** | *penA*-34 mosaic | 4 | 0.5 | 0.5 | 0.12 | 0.06 | 0.12 | 1 | 0.5 | 16 | 2 |
| **CDC-0213** | *penA*-34 mosaic | 4 | 0.25 | 0.5 | 0.06 | 0.06 | 0.12 | 0.5 | 0.12 | 16 | 1 |
| **CDC-0214** | *penA*-34 mosaic | 4 | 0.25 | 0.25 | 0.12 | 0.12 | 0.12 | 0.5 | 0.25 | 16 | 2 |
| **Summary** | | **12** | **15** | **18** | **21** | **CRO** | **ZOLI** | **GEPO** | **AZM** | **CIP** | **PEN G** |
| **MIC$_{50}$** | | 2 | 0.25 | 0.25 | 0.06 | 0.06 | 0.12 | 0.5 | 0.5 | 16 | 2 |
| **MIC$_{90}$** | | 4 | 0.5 | 0.5 | 0.12 | 0.12 | 0.12 | 1 | 1 | 16 | 2 |
| **MIC$_{100}$** | | 8 | 0.5 | 0.5 | 0.25 | 0.25 | 0.25 | 1 | 8 | 32 | 4 |

Abbreviations: CRO, ceftriaxone; ZOLI, zoliflodacin; GEPO, gepotidacin; AZM, azithromycin; CIP, ciprofloxacin; PEN G, penicillin G.

**Table S11**. Frequency of Resistance (FoR) data of boro-PBPi **21** and ceftriaxone.

| Compound | Concentration | ATCC 49226 | H041 | WHO Q |
|---|---|---|---|---|
| **Boro-PBPi 21** | **4× MIC** | $<1.56 \times 10^{-9}$ | $<1.79 \times 10^{-9}$ | $<4.17 \times 10^{-9}$ |
| **Boro-PBPi 21** | **16× MIC** | $<1.56 \times 10^{-9}$ | $<1.79 \times 10^{-9}$ | $<4.17 \times 10^{-9}$ |
| **Ceftriaxone** | **4× MIC** | $<1.56 \times 10^{-9}$ | $<1.79 \times 10^{-9}$ | $<4.17 \times 10^{-9}$ |
| **Ceftriaxone** | **16× MIC** | $<1.56 \times 10^{-9}$ | $<1.79 \times 10^{-9}$ | $<4.17 \times 10^{-9}$ |

**Table S12.** Safety and selectivity data for boro-PBPi **21**.

**a. Plasma protein binding (mouse and human)**

Mouse plasma protein binding data:

| Compound | Free (%) | Bound (%) | | Stability (%) | | Recovery (%) | |
|---|---|---|---|---|---|---|---|
| | Replicate | Replicate | Mean | Replicate | Mean | Replicate | Mean |
| **Boro-PBPi 21** | 82.68 | 17.32 | 18.97 | 94.58 | 95.47 | 93.11 | 93.13 |
| | 79.38 | 20.62 | | 96.36 | | 93.15 | |
| **Warfarin** | 3.23 | 96.77 | 96.62 | 96.21 | 98.11 | 89.90 | 89.02 |
| | 3.52 | 96.48 | | 100.00 | | 88.15 | |

Human plasma protein binding data:

| Compound | Free (%) | Bound (%) | | Stability (%) | | Recovery (%) | |
|---|---|---|---|---|---|---|---|
| | Replicate | Replicate | Mean | Replicate | Mean | Replicate | Mean |
| **Boro-PBPi 21** | 59.98 | 40.02 | 39.46 | 89.63 | 88.56 | 83.69 | 86.45 |
| | 61.09 | 38.91 | | 87.49 | | 89.21 | |
| **Warfarin** | 1.14 | 98.86 | 98.79 | 99.65 | 97.53 | 89.13 | 86.98 |
| | 1.29 | 98.71 | | 95.41 | | 84.82 | |

The plasma protein binding assays were performed by BioDuro-Sundia using the equilibrium dialysis method by incubating compound at 5 µM with plasma (mouse or human) in one compartment and phosphate buffer in the other compartment separated by a dialysis membrane (12–14K MWCO) for 5 h at 37°C with shaking. The amount of compound in each chamber compartment (buffer and plasma) was measured using reversed-phase high performance liquid chromatography/tandem mass spectrometry methods that were customized for each compound. The data were generated by BioDuro-Sundia.

**b. Cytotoxicity**

| Compound | Concentration tested (µg/mL) | MRC-5 MTS readout ($OD_{490}$ value) | HeLa MTS readout ($OD_{490}$ value) | 3T3 MTS readout ($OD_{490}$ value) |
|---|---|---|---|---|
| **Boro-PBPi 21** | **256** | 0.439 | 0.505 | 0.365 |
| | **81** | 0.588 | 0.455 | 0.348 |
| | **25.6** | 0.621 | 0.442 | 0.354 |
| | **8.1** | 0.511 | 0.396 | 0.338 |
| **Chlorohexidine (positive control)** | **256** | 0.115 | 0.161 | 0.191 |
| | **81** | 0.103 | 0.205 | 0.103 |
| | **25.6** | 0.083 | 0.146 | 0.256 |
| | **8.1** | 0.528 | 0.077 | 0.480 |
| **DMSO control (average ± standard deviation)** | - | 0.457 ± 0.094 | 0.411 ± 0.077 | 0.381 ± 0.021 |
| **Medium only (no cells) (average ± standard deviation)** | - | 0.050 ± 0.001 | 0.051 ± 0.002 | 0.056 ± 0.005 |

The half maximal cytotoxic concentration ($CC_{50}$) of each compound against three human cell lines (MRC-5 [ATCC CCL-171], HeLa [ATCC CCL-2], and 3T3-Swiss albino [ATCC CCL-92]) was measured by performing an MTS-based cell proliferation assay where actively dividing cells were used to maximize assay sensitivity. Procedurally, 180 µL of cells were seeded in cell-culture-treated 96-well plates (Celltreat, Pepperell, MA) at a density of 2–3.5 × $10^4$ cells/mL and incubated overnight at 37°C in a humidified, 5% $CO_2$ atmosphere. The next day, compound was diluted in PBS with 6% DMSO in a 0.5 $log_{10}$ dilution scheme at 10× the final compound concentration array. Compound titrations (in 20 µL volume) were added to each well of the cell culture plates to affect a final 10-fold dilution ranging from 8.1–256 µg/mL for boro-PBPi **21**. The final DMSO concentration in all assays was 0.6%. After an additional 72 h of incubation (when cells reached 90–100% confluency), CellTiter 96® $AQ_{ueous}$ solution (Promega, Fitchburg, WI) was added at 20 µL per well and plates were incubated an additional 2–4 hours at 37°C, 5% $CO_2$. The optical density of the wells was read on a Biotek Cytation 3 plate reader at 490 nm. Background signal (wells receiving Cell Titer reagent but containing no cells) was subtracted from all wells on the plate. If the cell absorbance was ≥50% when compared to the DMSO control wells, the compound was considered to have no cytotoxicity and the $CC_{50}$ was recorded as greater than the highest concentration tested. If the cell absorbance was <50% when compared to the cell control wells, percent growth versus $log_{10}$ drug concentration was plotted and the concentrations at which 50% of the cells survived compared to growth control ($CC_{50}$) were calculated using a sigmoidal four parameter curve fit in GraphPad Prism.

**c. Hemolysis**

| Compound | Concentration tested | OD541 value | OD541 background (DMSO/water only) | % Hemolysis |
|---|---|---|---|---|
| **Boro-PBPi 21** | 1 mg/mL | 0.058 | 0.079 | -0.99 |
| **Water** | - | 2.203 | 0.000 | 100.00 |

Human red blood cells (Rockland Immunochemicals, Inc., #R4070050) were diluted in PBS to obtain a target blood cell concentration of $5 \times 10^8$ erythrocytes /mL. Desired test compound stock solutions were prepared in 1.2 mL PBS alongside appropriate controls. The diluted blood cells (300 µL) were added to each tube and gently inverted to mix. Tubes were incubated for 1 h at 37°C, gently inverted to mix, and centrifuged at 11,000 × *g* for 5 min. The OD at 541 nm of the supernatant was measured for each desired test sample and control and the % hemolysis was calculated. In vitro hemolysis values of <10% are considered nonhemolytic[3].

**d. Mitochondrial toxicity**

| Compound | Media | $IC_{50}$ (µM) | Fold shift (Glu/Gal) |
|---|---|---|---|
| **Antimycin** | Glucose | >25 | >3 |
| | Galactose | 0.00205 | |
| **Staurosporine** | Glucose | 0.80270 | 0.70 |
| | Galactose | 1.15300 | |
| **Boro-PBPi 21** | Glucose | >100 | Not applicable |
| | Galactose | >100 | |

The assay to monitor mitochondrial toxicity was performed by BioDuro-Sundia. Briefly, SKOV-3 cells (ATCC HTB-77) were seeded and cultured in media containing either glucose or galactose overnight. Test compounds [Boro-PBPi **21**, antimycin, and staurosporine] were added at 6 concentrations (threefold serial dilution from 100 μM to 0.4 μM) and incubated with the cells for 24 h. Cell viability was measured by the CellTiter-Glo® method.

#### e. Chromosomal aberrations (micronucleus assay)

#### Micronucleus in CHO-K1 cells without S9

| Compound | Concentration (μM) | Scored Cells (Number of Binucleated Cells) | | % Micronucleated Cells | | % Cytotoxicity CBPI Index | | t-Test p-value | Test Result |
|---|---|---|---|---|---|---|---|---|---|
| **Boro-PBPi 21** | 300 | 13216 | 13916 | 1.97% | 2.34% | 3.77 | 4.46 | 0.4730 | - |
| | 100.000 | 14274 | 14221 | 2.23% | 2.01% | 2.38 | 3.36 | 0.5003 | - |
| | 33.333 | 13899 | 14177 | 2.26% | 2.14% | 0.52 | 2.63 | 0.3105 | - |
| | 11.111 | 13607 | 14643 | 2.49% | 2.19% | 0.37 | 1.78 | 0.1847 | - |
| | 3.704 | 13839 | 14309 | 2.23% | 1.86% | 1.94 | 1.99 | 0.7625 | - |
| | 1.235 | 14225 | 14642 | 1.98% | 1.96% | -0.72 | 2.32 | 0.9905 | - |
| | 0.412 | 14176 | 14076 | 2.19% | 2.14% | 0.78 | 2.19 | 0.3625 | - |
| | 0.137 | 14301 | 14562 | 2.17% | 1.94% | -0.64 | 1.15 | 0.6981 | - |

| Compound | Concentration (μM) | Scored Cells | % Micronucleated Cells | % Cytotoxicity CBPI Index | Test Result |
|---|---|---|---|---|---|
| **Untreated control** | - | 14574 | 2.15% | 3.24 | NA |
| | - | 14693 | 2.06% | -2.17 | |
| | - | 14998 | 1.69% | -1.08 | |
| **Mitomycin C** | 1.85 | 4277 | 13.82% | 62.34 | + |
| | 1.85 | 4744 | 13.76% | 61.80 | |
| | 1.85 | 5006 | 13.82% | 58.95 | |

#### Micronucleus in CHO-K1 cells with S9

| Compound | Concentration (μM) | Scored Cells (Number of Binucleated Cells) | | % Micronucleated Cells | | % Cytotoxicity CBPI Index | | t-Test p-value | Test Result |
|---|---|---|---|---|---|---|---|---|---|
| **Boro-PBPi 21** | 300 | 4759 | 5332 | 3.99% | 3.68% | 4.33 | 6.52 | 0.3734 | - |
| | 100.000 | 4429 | 5334 | 5.08% | 4.48% | 12.30 | 4.88 | 0.0244 | - |
| | 33.333 | 4437 | 3433 | 4.28% | 6.00% | 12.98 | 18.25 | 0.1056 | - |
| | 11.111 | 4650 | 3116 | 3.78% | 5.55% | 12.81 | 34.47 | 0.2223 | - |

| | | | | | | | | | |
|---|---|---|---|---|---|---|---|---|---|
| | 3.704 | 3577 | 3584 | 4.75% | 5.69% | 16.77 | 29.55 | 0.0254 | - |
| | 1.235 | 3222 | 3953 | 5.12% | 5.89% | 24.93 | 26.16 | 0.0106 | - |
| | 0.412 | 4351 | 3370 | 3.40% | 5.93% | 13.50 | 24.78 | 0.3570 | - |
| | 0.137 | 3913 | 3684 | 4.75% | 6.30% | 20.22 | 28.65 | 0.0501 | - |

| Compound | Concentration (μM) | Scored Cells | % Micronucleated Cells | % Cytotoxicity CBPI Index | Test Result |
|---|---|---|---|---|---|
| **Untreated control + S9** | - | 5453 | 3.70% | 5.91 | NA |
| | - | 5604 | 3.39% | -3.43 | |
| | - | 5538 | 3.79% | -2.48 | |
| **Mitomycin C** | 22.375 | 1669 | 13.48% | 56.08 | + |
| | 22.375 | 1868 | 10.76% | 60.22 | |
| | 22.375 | 1709 | 10.94% | 48.35 | |

The micronucleus assay to monitor genotoxicity of boro-PBPi **21** was performed by BioDuro-Sundia as described previously[4]. Briefly, CHO-K1 cells (ATCC CCL-61)) were treated with **21** at concentrations up to 300 μM with and without incubation with liver S9 fraction, followed by incubation for 24 h. The cells were fixed by 4% paraformaldehyde and stained with 4,6-diamidino-2-phenylindole (DAPI). The micronucleus rate with CHO cells was evaluated using a High Throughput Screening system. As positive controls, CHO-K1 cells were incubated with cyclophosphamide with S9 and mitomycin C without S9, where chromosomal damage and micronucleated cells were observed relative to untreated controls.

#### f. CYP450 inhibition data

| CYP450 enzyme | Substrate used | % Enzyme Activity (Relative to DMSO controls) | | Positive control compound used | Positive control $IC_{50}$ (μM) |
|---|---|---|---|---|---|
| | | Boro-PBPi 21 at 30 μM | | | |
| | | Data 1 | Data 2 | | |
| **CYP1A2** | Vivid™ EOMCC | 96.16 | 97.67 | **Furafylline** | 1.70 |
| **CYP2C19** | Vivid™ EOMCC | 64.27 | 66.47 | **Ketoconazole** | 2.96 |
| **CYP2D6** | Vivid™ EOMCC | 98.28 | 103.14 | **Ketoconazole** | 9.59 |
| **CYP3A4** | Vivid™ BOMCC | 100.61 | 102.47 | **Ketoconazole** | 0.0052 |

| CYP450 enzyme | Substrate used | % Enzyme Activity (Relative to DMSO controls) Boro-PBPi 21 at 30 µM | | Positive control compound used | Positive control $IC_{50}$ (µM) |
|---|---|---|---|---|---|
| | | Data 1 | Data 2 | | |
| **CYP2E1** | Vivid™ EOMCC | 92.83 | 95.60 | **Tranylcypromine** | 91.9 |
| **CYP2B6** | Vivid™ BOMCC | 107.41 | 109.00 | **Miconazole** | 0.036 |
| **CYP2C9** | Vivid™ OOMR | 96.60 | 95.24 | **Ketoconazole** | 3.03 |
| **CYP2C8** | Vivid™ DBOMF | 101.96 | 103.27 | **Miconazole** | 3.76 |

Inhibition of CYP450 enzymes was monitored with boro-PBPi **21** at 30 µM in fluorescence-based assays using Vivid™ substrates, relative to DMSO controls, at Reaction Biology (Malvern, PA). The $IC_{50}$s of Positive control compounds were also determined to validate the assays. The assays were performed at Reaction Biology (Malvern, PA).

**g. hERG binding data**

| Compound | % tracer binding | | Positive control $IC_{50}$ (µM) |
|---|---|---|---|
| | Data 1 | Data 2 | |
| **DMSO** | 102.20 | 95.48 | ND |
| **Boro-PBPi 21 at 30 µM** | 88.32 | 94.99 | ND |
| **E4031 (positive control)** | NA | NA | 0.019 |

Binding to hERG in the membrane for boro-PBPi **21** was monitored in the fluorescence-based assay using 1 nM Predictor™ hERG Tracer Red and 1X Predictor™ hERG membrane. Fluorescence was measured using excitation and emission at 531 nm and 595 nm, respectively. The assay was performed at Reaction Biology (Malvern, PA).

**h. Protease inhibition data**

| Target: | % Enzyme Activity (relative to DMSO controls) | Control compound |
|---|---|---|

| | Boro-PBPi 21 at 30 μM | | | Control Compound $IC_{50}$ (M) |
|---|---|---|---|---|
| | **Data 1** | **Data 2** | | |
| **Chymotrypsin** | 82.12 | 82.35 | **Chymostatin** | 7.27E-10 |
| **Thrombin a** | 89.45 | 91.98 | **Gabexate mesylate** | 1.49E-06 |
| **Trypsin** | 94.04 | 90.85 | **Gabexate mesylate** | 2.40E-08 |

The protease activities were monitored as a time-course measurement of the increase in fluorescence signal from fluorescently labeled peptide substrate, and initial linear portion of slope (signal/min) was analyzed. The assays were performed at Reaction Biology (Malvern, PA).

**Table S13.** Bacterial strains used.

| Species | Strain ID | Strain information | Source/reference |
|---|---|---|---|
| *N. gonorrhoeae* | ATCC 49226 | CLSI QC strain | ATCC |
| *N. gonorrhoeae* | FA19 | Wild type, penicillin-susceptible laboratory strain | Robert Nicholas[5] |
| *N. gonorrhoeae* | FA1090 | ATCC 700825 | Ann Jerse[6] |
| *N. gonorrhoeae* | WHO F | CDC-0901 | CDC[7] |
| *N. gonorrhoeae* | WHO G | CDC-0902 | CDC[7] |
| *N. gonorrhoeae* | WHO M | CDC-0905 | CDC[7] |
| *N. gonorrhoeae* | MS11 | | Ann Jerse[8] |
| *N. gonorrhoeae* | WHO L | CDC-0904 | CDC[7] |
| *N. gonorrhoeae* | WHO K | CDC-0903 | CDC[7] |
| *N. gonorrhoeae* | H041 | WHO X, CDC-0912 | Ann Jerse |
| *N. gonorrhoeae* | CDC-0914 | WHO Z, A8806 | CDC |
| *N. gonorrhoeae* | WHO Q | G7944, NCTC 14208 | NCTC[9] |
| *N. gonorrhoeae* | CDC-0197 | | CDC |
| *N. gonorrhoeae* | F89 | WHO Y, CDC-0913 | Ann Jerse[10] |
| *N. gonorrhoeae* | H041($STR^R$) | H041 streptomycin-resistance, used in the in vivo efficacy study | Ann Jerse[2] |
| *N. gonorrhoeae* | CDC set | 44 CDC strains | CDC |
| *Staphylococcus aureus* | ATCC 29213 | Methicillin-sensitive | ATCC |
| *Escherichia coli* | ATCC 25922 | CLSI QC strain | ATCC |
| *E. coli* | BAS901C | MC4100 Δ*lamB106 zab*::Tn5 *lptD4123*, drug-susceptible | Eric Brown[11, 12] |
| *Klebsiella pneumoniae* | UMM3 | KPC-2 | Jean-Denis Docquier[13] |
| *Pseudomonas aeruginosa* | ATCC 27853 | CLSI QC strain | ATCC |
| *P. aeruginosa* | ATCC 35151 | *lptD4123*, drug-susceptible | ATCC |
| *Acinetobacter baumannii* | ATCC 19606 | Type strain | ATCC |

ATCC, the American Type Culture Collection; CDC, the Centers for Disease Control and Prevention; NCTC, the National Collection of Type Cultures; CLSI, the Clinical Laboratory Standard Institute.

**Table S14.** Crystallographic data collection and model refinement statistics.

| | **tPBP2$^{35/02}$-12** | **tPBP2$^{35/02}$-15** |
|---|---|---|
| **Data collection** | | |
| Resolution range | 38.7 – 2.6 (2.7-2.6) | 38.9-1.89 (1.96-1.89) |
| Space group | $P\ 2_1 2_1 2_1$ | $P\ 2_1 2_1 2_1$ |
| Unit cell *a, b, c* (Å) | | 50.6, 60.7, 110.3 |
| Total reflections | 76,034 (7,191) | 197,257 (19,553) |
| Unique reflections | 10,571 (975) | 27,662 (2,643) |
| Multiplicity | 7.2 (7.4) | 7.1 (7.4) |
| Completeness (%) | 98.9 (94.3) | 99.3 (96.1) |
| Mean I/sigma(I) | 14.1 (4.2) | 15.9 (3.1) |
| R-merge | 0.199 ((0.713) | 0.109 (0.708) |
| R-pim | 0.080 (0.283) | 0.044 (0.276) |
| $CC_{1/2}$ | 0.988 (0.831) | 0.998 (0.839) |
| **Refinement** | | |
| R-factor (%) | 17.8 | 17.4 |
| R-work (%) | 17.5 (22.7) | 17.3 (21.3) |
| R-free (%) | 23.2 (29.4) | 20.5 (26.3) |
| No. of non-hydrogen protein atoms | 2,506 | 2,471 |
| No. of ligand atoms | 39 | 40 |
| No. of waters | 19 | 122 |
| RMSDs from ideal stereochemistry: | | |
| RMS (bonds) | 0.008 | 0.009 |
| RMS (angles) | 1.57 | 1.58 |
| Ramachandran plot: | | |
| Core (%) | 92.0 | 92.7 |
| Allowed (%) | 8.0 | 6.5 |
| Generously allowed (%) | 0.0 | 0.7 |
| Outliers (%) | 0.0 | 0.0 |
| B-factors: | | |
| Mean B-factor (all atoms) (Å$^2$) | 29.9 | 26.0 |
| Macromolecules (Å$^2$) | 29.7 | 25.8 |
| Ligands (Å$^2$) | 42.0 | 30.8 |
| Solvent (Å$^2$) | 19.0 | 29.5 |
| PDB code | 9MD0 | 9MCZ |

**Table S15.** Liquid chromatography parameters in the final bioanalytical method for boro-PBPi **21**.

| Parameter | Value | |
|---|---|---|
| **Ion Mode** | Positive | |
| **Mobile Phase A** | $H_2O$ + 0.5% formic acid | |
| **Mobile Phase B** | ACN + 0.5% formic acid | |
| **Weak needle wash** | 90/10 $H_2O$/MeOH + 0.02%$NH_4OH$ (pH 10) | |
| **Strong needle wash** | 10/90 $H_2O$/ACN | |
| **Seal wash** | 90/10 $H_2O$/ACN | |
| **Column** | Waters Acquity Premier BEH C18 50*2.1mm, 1.7 µm | |
| **Column Temperature (°C)** | 25 | |
| **Auto-sampler Temperature (°C)** | 5 | |
| **Injection volume (µL)** | 7.5 | |
| **Gradient** | | |
| **Time (min)** | Flow (mL/min) | %MPB |
| **0** | 0.5 | 15 |
| **0.2** | 0.5 | 15 |
| **0.9** | 0.5 | 30 |
| **1.6** | 0.5 | 95 |
| **2** | 0.5 | 95 |
| **2.01** | 0.5 | 15 |
| **2.5** | 0.5 | 15 |

**Table S16.** MS/MS setting in the final bioanalytical method for boro-PBPi **21**.

| | |
|---|---|
| Source temperature (°C) | 150 |
| Desolvation Temperature (°C) | 500 |
| Cone Gas flow (L/Hr) | 60 |
| Desolvation gas flow (L/min) | 1000 |

| **Compound** | **Q1(m/z)** | **Q3(m/z)** | **Collision Energy(V)** | **Cone Voltage (V)** |
|---|---|---|---|---|
| **Boro-PBPi 21** | 647.04 | 629.1 | 16 | 20 |
| Levofloxacin (IS) | 362.2 | 261.2 | 20 | 26 |

IS = internal standard

**Table S17.** Validation summary for boro-PBPi **21** bioanalytical method in rat plasma.

| Parameter | Acceptance criteria | Pass/Fail |
|---|---|---|
| **Linearity** | $R^2$ value ≥ 0.98 | Passed |
| **Sensitivity** | ± 20% bias (with less than 20% CV) | Passed |
| **Selectivity** | LLOQ level: ± 20% bias<br>Blank: analyte signal ≤ 20% of the average analyte signal in LLOQ samples<br>DB: analyte signal ≤ 20% of the average analyte signal in LLOQ samples; IS signal ≤ 5% the average IS signal in LLOQ samples. | Passed |
| **Accuracy and precision** | ± 15% bias (with less than 15% CV) | Passed |
| **Dilution factor** | ± 15% bias (with less than 15% CV) | Passed for 10X |
| **Benchtop stability** | ± 15% bias (with less than 15% CV) | Passed for 3.5 hours at RT |
| **Freeze/Thaw stability** | ± 15% bias (with less than 15% CV) | Passed for 4 cycles between -80°C and RT |
| **Reinjection stability** | ± 15% bias (with less than 15% CV) | Passed for 49 hours at autosampler temperature |
| **Hemolysis** | ± 15% bias (with 15% CV) | Passed for 2% hemolyzed plasma |
| **Interference to IS from analyte** | ≤ 5% of the average IS signal | 0% |
| **Matrix effect** | | 1.11 (enhancement) |
| **Recovery** | | 45.6% |

Blank: samples with no analyte but IS was added.

Double blank: samples with no analyte and IS.
